# The first crested duck genome reveals clues to genetic compensation and crest cushion formation

**DOI:** 10.1101/2021.07.25.452189

**Authors:** Guobin Chang, Xiaoya Yuan, Qixin Guo, Hao Bai, Xiaofang Cao, Meng Liu, Zhixiu Wang, Bichun Li, Shasha Wang, Yong Jiang, Zhiquan Wang, Yang Zhang, Qi Xu, Qianqian Song, Rui Pan, Shenghan Zheng, Lingling Qiu, Tiantian Gu, Xinsheng Wu, Yulin Bi, Zhengfeng Cao, Yu Zhang, Yang Chen, Hong Li, Jianfeng Liu, Wangcheng Dai, Guohong Chen

## Abstract

The Chinese crested (CC) duck is a unique indigenous waterfowl breed with a phenotypic crest trait that affects its high survival rate. Therefore, the CC duck is an ideal model to investigate the genetic compensation response to maintain genetic stability. In the present study, we first generated a chromosome-level genome of CC ducks. Comparative genomics revealed genes related to tissue repair, immune function, and tumors were under strong positive selection, which suggested that these adaptive changes might enhance cancer resistance and immune response to maintain the genetic stability of CC ducks. We sub-assembled a Chinese spot-billed duck genome and detected genome-assembled structure variants among three ducks. Functional analysis revealed that a large number of structural variants were related to the immune system, which strongly suggests the occurrence of genetic compensation in the anti-tumor and immune systems to further support the survival of CC ducks. Moreover, we confirmed that the CC duck originated from the mallard ducks. Finally, we revealed the physiological and genetic basis of crest traits and identified a causative mutation in *TAS2R40* that leads to crest formation. Overall, the findings of this study provide new insights into the role of genetic compensation in adaptive evolution.

## Introduction

Organisms have developed dynamic buffer systems during evolution to maintain normal development in the presence of certain genetic mutations[1–4]. Organisms adapt to their environments by genomic fine-tuning during their evolution. Recently, the genetic compensation response (GCR), a new mechanism supporting genomic robustness, was found in zebrafish [5, 6], mice [7] and rockcress [8] by gene-knockout mutations. In a sense, the organism developed a lethal phenotype caused by harmful mutations, or in these instances resulting ’similar to gene knockout. Under the action of long-term natural selection and artificial selection, the GCR causes a series of genetic compensation mutations, thereby promoting genetic stability to maintain the organism. Over time, compensation mutations may lead to a series of phenotypic changes that offset the lethal phenotype to maintain the population.

The Chinese crested (CC) duck is a unique breed with complex feather-protruding traits that are collectively termed the crest. Feather crests are widely distributed in birds (such as cockatoos, grey-crowned cranes, and great-crested grebes), although there are significant differences in shape and physiological mechanisms. Almost all birds with crest traits exhibited a distinct crown formed by prominent feathers. The crest cushion of the CC duck consists of soft tissue protuberances covered by feathers and skin. While the presence of a crest does not affect survival in most crested birds, crested ducks are an exception. Previous studies of ‘Hochbrutflugenten’ (HBTcr) ducks, which are crested duck breeds in Germany, have shown that crested ducks have high pre- and postnatal mortality, exhibiting motor incoordination in the wild due to incomplete skull closure [9–11]. Although the phenotype composition of the crest cushion and the fertilization rate in HBTcr and CC ducks were similar, the survival rate of CC ducks was significantly higher (more than 95%) after birth with good motor coordination. Therefore, the formation mechanism of the crest cushion and the genomic compensation for the effect of the crest cushion on the CC duck has gathered considerable interest in CC duck research. However, resolving this issue has proven challenging because the crest trait is phenotypically complex. Nevertheless, the CC duck is an ideal example to help explain the function of the GCR in maintaining genetic stability. Specifically, genome assembly may be the best solution to address these issues. However, the genomic resources for duck are limited, with published genome sequences limited to Peking duck, mallard duck, and Shaoxing duck in the NCBI database [12–14]. In addition, these genomes cannot reveal the basis of crested cushion formation at the genomic level.

To explore the physiological and genetic basis behind the formation of crest cushions, we first assembled a high-quality CC duck genome and a Chinese spot-billed duck (Csp-b duck; *Anas zonorhyncha*) genome. These genomes were compared to those of other wild and domesticated ducks to investigate shifts in structural variants and genes under adaptive evolution. Our results provide valuable insights for understanding the role of the GCR in adaptive evolution and provide a valuable genomic resource for future genome-wide analyses of economically important traits in poultry.

## Results

### Genome assembly of the CC duck

A 28-week-old female CC duck was selected for genome sequencing and assembly. The genome size was estimated to be 1.26 Gb based on the *k*-mer distribution (Figure S1 and Table S1). To generate a high-quality reference genome for CC duck, a total of 85.06 Gb (∼75.97x) PacBio long reads were assembled using FALCON v0.7 [15] and this assembly was then polished using Quiver (smrtlink v6.0.1) [16]. Thereafter, 10× Genomics (∼79.15x) was used to connect contigs into super-scaffolds with the software fragScaff [17], which resulted in a 1.13 assembly (CC_duck_v1.0) with an N50 contig size of 3.24 Mb and an N50 scaffold size of 7.61 Mb (Table S3). Approximately 88.65 Gb (∼79.15x) Illumina paired-end reads were used to polish the assembly with Pilon v1.18. Using the high linkage genetics map, 1,216 scaffolds were anchored and oriented onto 37 autosome chromosomes using CHROMONMER [18] (Figure S3). The remaining scaffolds were organized into the CC duck Z chromosomes based on their sequence similarity with the Z chromosomes of the published duck genome (CAU_duck1.0) by MUMmer v3.23. The final assembly yielded an N50 scaffold size of 73.74 Mb and an N50 contig size of 3.24 Mb, and ∼94.10% of the assembly genome was anchored onto the 38 chromosomes (i.e., 37 autosome chromosomes and one Z chromosome) (Figure 1 and Table S4). To assess the quality and integrity of the genome assembly, short paired-end reads were aligned with the assembly. Overall, 96.68% of the paired-end reads could be mapped to the genome, suggesting high integrity of our assembly genome (Table S5). Benchmarking Universal Single Copy Orthologs (BUSCO) [19] showed that 97.7% (2,527/2,586) of vertebrate single-copy orthologous genes were captured in our assembly, which was comparable or even better than that in published duck genomes (Table S6). A total of 17,425 protein-coding genes were predicted in the CC duck genome by combining *de novo*, homology-based, and RNA-sequence gene prediction methods (Table S7). In addition, to help explore the origin and adaptive evolution of CC ducks, we assembled another duck genome (Csp-b duck) with ∼85.81 × paired-end reads using SOAPdenovo (Supplementary Note 8 and Table S32) [20]. Finally, we generated a 1.10 assembly with an N50 scaffold size of 675.96 kb (Table S8) and predicted 15,278 protein-coding genes based on homologous comparison approaches (Table S9).

**Figure 1.**
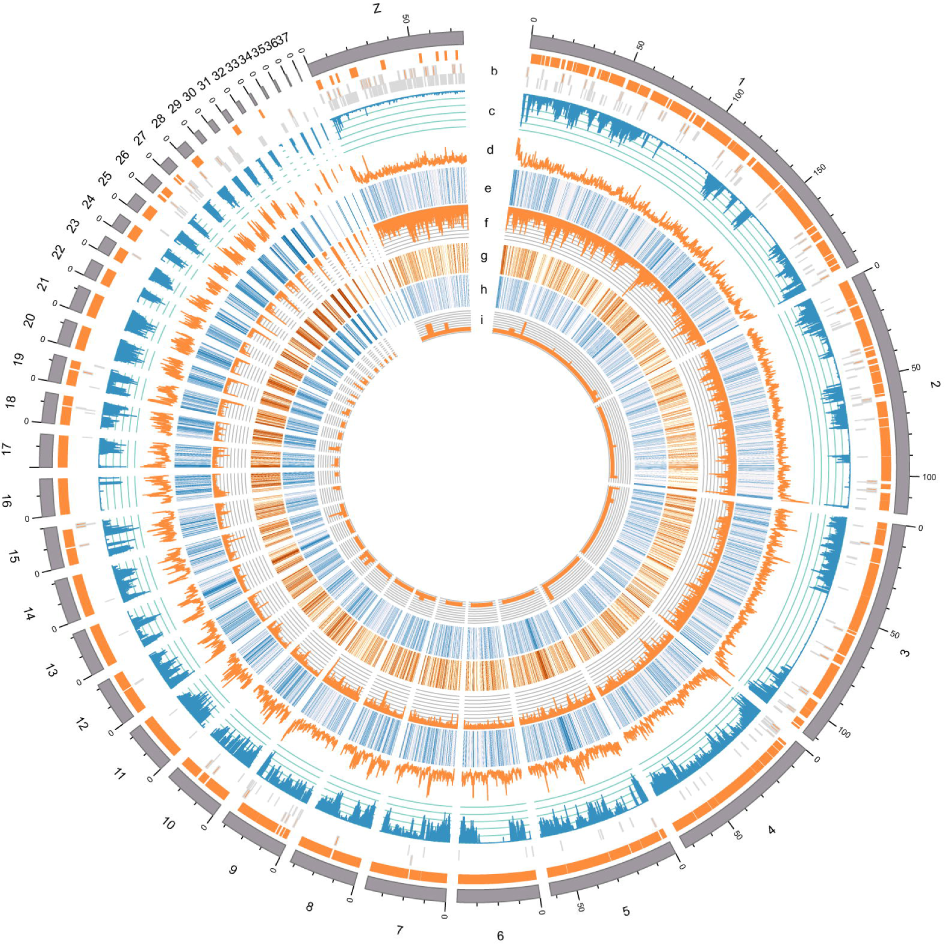
Landscape and linkage groups of the Chinese crested duck genome. The circles illustrate from the outside to inside: a) chromosome ideograms for CC duck; b) schematic depicting assembly contig lengths for CC duck chromosomes. The first track of orange rectangles represent contigs >1 Mb, the second track correspond to contigs ≤1 Mb and > 500 kb, and the third track represent the other contigs; c) distribution of SNP density (non-overlapping, window size, 500 kb) in each chromosome; d) distribution of GC content (non-overlapping, window size, 500 kb); e) distribution of gene content (non-overlapping, window size, 500 kb); f) distribution of repeat content (non-overlapping, window size, 500 kb); g) heat map showing the expression level of genes in crested-head tissue, represented by FPKM values; h) heat map showing the expression level of gene in skin tissue, represented by FPKM values; i) distribution of miRNA number (non-overlapping, window size, 500 kb).

### Historical population structure reveals the origin of the CC duck

The CC duck is a unique domesticated duck breed with a crest cushion in China. According to historical records, the first documented origin of the CC duck can be traced back to the early Ming Dynasty in China (A.D. 1368) and may have been present earlier (Figure S4). To explore the origin of the crest cushion, we obtained data from three wild duck breeds (mallard duck from Ningxia Province (MDN), mallard duck from Zhejiang Province (MDZ), and Csp-b) and two domesticated duck breeds (Pekin duck and CC duck) (Table S10). After excluding linked SNP loci that could potentially bias clustering results, we built a neighbor-joining (NJ) tree using 39 samples. The NJ tree assigned all samples to three major groups (the wild duck, Pekin duck, and CC duck groups) (Figure 2a). These clustered results were also supported by principal component analysis (PCA) (Figure 2b). Additionally, we used FRAPPE to explore the genetic composition of each group after initially removing potential bias caused by missing loci [21]. The sample clusters were evaluated using an ad hoc statistic (Δ*K*). The domesticated ducks were separated from the wild type ducks when the cluster number *K* was set to 2. The ΔK value reached its maximum at *K* = 3, indicating the uppermost structural level. At the same time, the MDZ was also separated from the MDN. At *K* = 4, the clusters revealed that the Pekin duck shared gene flow with the CC duck (Figure 2c). The results of the NJ tree, PCA, and FRAPPE indicated that there was gene flow between the Pekin and CC ducks. We inferred that the CC duck might have been domesticated independently from the MDZ.

**Figure 2.**
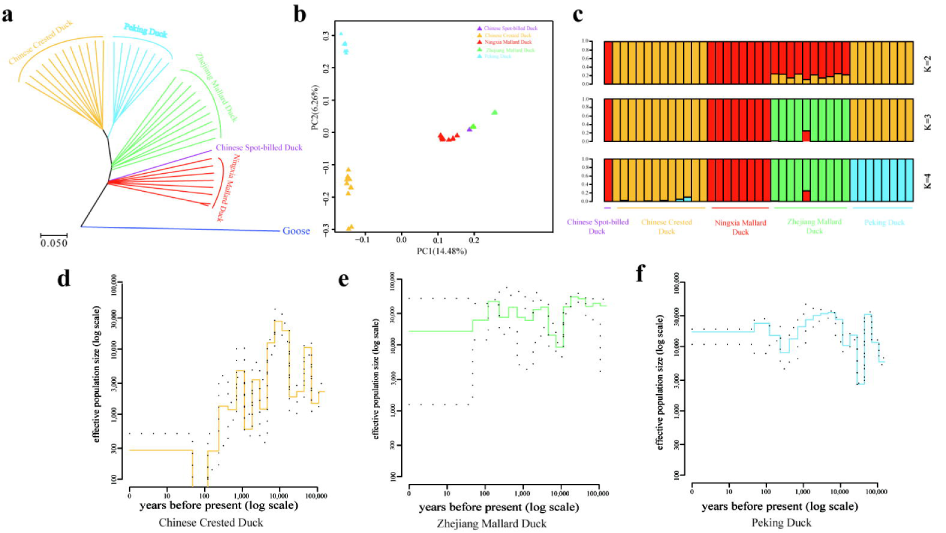
Population structure, genomic landscape of duck divergence, and population size estimate. a) The principal components analysis of the duck samples. b) Phylogenetic tree (neighbor-joining tree with 1,000 bootstraps) of all samples inferred from the whole-genome tag SNPs with geese (*Anser cygnoides*) as an outgroup. c) Population structure of all individuals (*K* = 2, 3, and 4). The population origin of each individual is indicated on the x-axis. Each individual is represented by a bar that is segmented into colors based on the ancestry proportions given the assumption of *K* populations. d–f) The recent effective population size (*Ne*) for CC duck (d), MDZ (e), and Pekin duck (f) were inferred using PopSizeABC v2.1. A 90% confidence interval is indicated by dotted lines.

Broad-scale population collection and management of CC ducks are critical for population recovery. Such efforts are challenging because the historical population scale of CC ducks is unclear. To infer the ancient demographic history of the CC ducks, the PopSizeABC method, which is based on approximate Bayesian computation, was used to predict the effective population size (*N_e_*) of CC ducks, Pekin ducks, and mallard ducks over the past 100,000 years. Over this period, we found that the population size of CC (Figure 2d), mallard (Figure 2e), and Pekin ducks (Figure 2f) varied significantly in the degree of fluctuations followed by a short period of relative population stability before the near extinction of the CC duck in the past 100 years. The recent demographic pattern implies that the CC duck experienced a population increase over the past 70 years through human protection beginning in the 20^th^ century.

### Gene evolution related to the GCR and crested trait formation of CC duck

To reveal the genomic signatures of the GCR in the adaptive evolution of the CC duck, its genome and 13 other published species genomes (Table S11) were selected for gene family clustering analysis using OrthoMCL software [22], which identified 19,605 gene families including 3,089 single-copy gene families. Based on 3,089 single-copy genes, we constructed a phylogenetic tree and estimated the divergence times of these 14 species. Phylogenomic analysis showed that the CC duck diverged from goose ∼23.3 million years ago (Mya)—slightly earlier than the previous molecular-based estimate of 20.8 Mya for ducks and geese [23] (Figure 3a). Interestingly, we found that the crest cushion also exists in every branch of the gray-crowned crane, great-crested grebe, hoatzin, and little egret. Other species in the phylogenetic tree also possess a crest, such as crested cockatoo, crested pigeons, and emperor penguins, but the type and function of their crests may vary. Therefore, we considered that the crest might be widespread in all types of birds, although the crested characters were removed or preserved in some birds under natural selection or human intervention.

**Figure 3.**
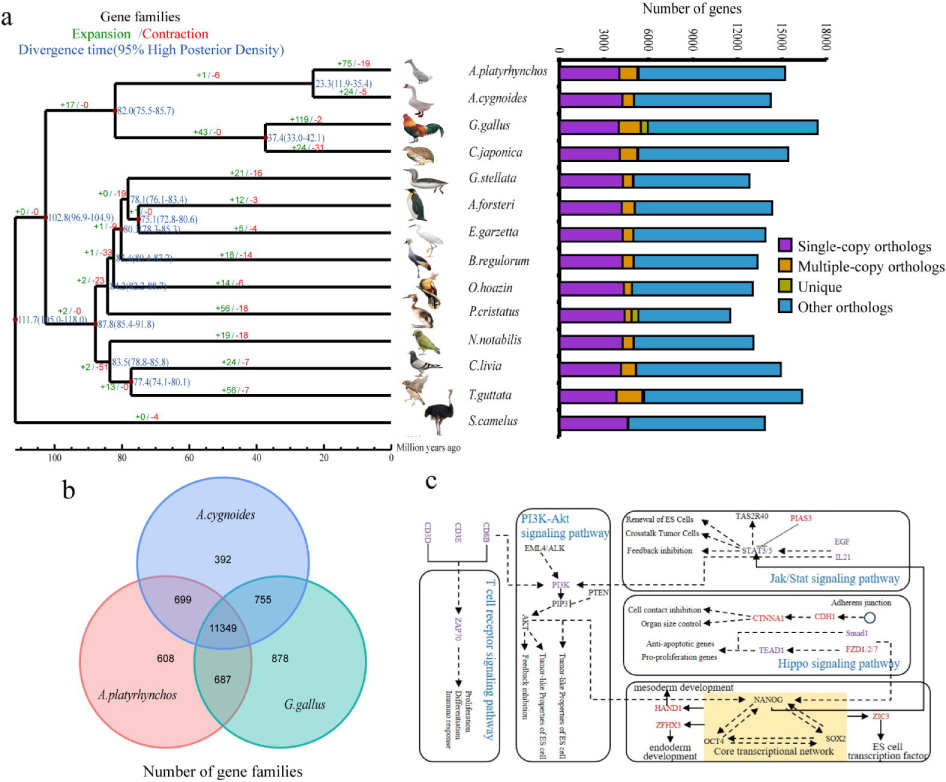
Comparative genomics reveals clues to genetic compensation of CC duck. (a) Phylogenetic tree constructed using single-copy orthologs. The estimated divergence time is shown in the middle of the branch (blue). The expansion gene (green) and contraction gene (red) above the branch. The gene family cluster is shown to the right of the phylogenetic tree. (b) Lineage-specific genes (LSGs) of CC duck identified by comparison with chicken, Pekin duck and geese. (c) Proposed signaling pathways for the protective mechanism and crest formation mechanism in CC duck. The positively selected genes in CC duck are indicated in purple, and expanded genes are indicated in red.

### Gene family evolution

Gene family expansion and contraction were examined using CAFE software [24] . Compared to the most recent common ancestor (MRCA), we identified 75 expanded gene families and 19 contracted gene families in CC ducks (Figure 3a). Furthermore, these expanded gene families were mainly enriched in gene ontology (GO) terms including cell adhesion, intracellular non-membrane-bound organelles, Wnt-activated receptor activity, and interleukin-1 receptor binding. KEGG enrichment analyses predicted that these genes were involved in the Hippo signaling pathway, cell adhesion molecules, gap junctions, and signaling pathways regulating the pluripotency of stem cells. We speculated that these expanded genes in CC ducks may potentially participate in special phenotypic evolution, such as that of the crest cushion (Tables S11–S14). In addition, the tripartite motif containing 39 (*TRIM39*) and 7 (*TRIM7*), which are parts of the contracted genes, have been implicated in the immune system [23, 25]. The TRIM gene family has been shown to be involved in some tumor mechanisms due to E3-ubiquitin ligase activity [26]. These results might provide the basis for phenotypic plasticity and compensate for the effect of the crest cushion. Compared with the chicken and goose genomes, we identified 608 gene families specific to CC duck, many of which were involved in cell adhesion molecules, focal adhesion, and calcium ion binding, especially tissue repair and tumor formation pathways (Table S15–S16), suggesting that these genes might play an essential role in cancer development, diffusion, and tissue repair. Collectively, we considered that human intervention led to adaptive evolution in the protective mechanisms of certain species.

### Positive selection of genes involved in the anti-tumor response

Positive selection has undoubtedly played an important role in the evolution of animals, especially in maintaining some endangered species. We identified 479 positive selection genes (PSGs) in the CC duck lineage. Functional enrichment analyses showed that these PSGs were significantly associated with genomic stability and tumor formation and were assigned terms including mismatch repair, cellular response to DNA damage stimulus, DNA double-strand break repair telomere maintenance *via* telomerase, and cancer, which may be the underlying basis for the crest cushion (Tables S17–S18). Importantly, we found that several key genes were under positive selection, such as epidermal growth factor (*EGF*), phosphatidylinositol-4,5-bisphosphate 3-kinase catalytic subunit delta (*PIK3CD*), and phosphoinositide-3-kinase regulatory subunit 4 (*PIK3R4*), which are involved in PD-L1 expression and the PD-1 checkpoint pathway, providing further evidence that the evolution of the crest cushion relies on tumor formation. Furthermore, we found that the Fraser extracellular matrix complex subunit 1 (*FRAS1*) gene was also under positive selection in CC ducks, which provides evidence that *FRAS1* is associated with hair curliness [27]. In addition, certain genes (e.g., golgin, RAB6-interacting (*GORAB*), Fas cell surface death receptor (*FAS*), etc.) in the p53 pathway were also positively selected. These findings suggest that CC ducks might have enhanced *GORAB* and reduced mouse double minute 2 homolog (*MDM2*) expression during evolution, thereby promoting p53 escape and activating the apoptosis pathway [28]. We also found some proto-oncogenes under strong positive selection in CC ducks, such as key genes in the PI3K-Akt signaling pathway (*PIK3CD*, *PIK3R4*, collagen type VI alpha 1 chain (*COL6A1*), *EGF*, laminin subunit alpha 1 (*LAMA1*), and von Willebrand factor (*VWF*); Figure 3c). These results suggest that genetic complementation mutations might have occurred at the genomic level. The effect of this compensation variation was amplified by artificial protection, allowing the CC duck to continue to survive or even expand its population.

In addition, to identified the expression level of PSGs during the crested cushion development, we compared the crest region and adjacent frontal skin tissues at each important embryo development stage to identify differentially expressed genes (DEGs) (Figure S5). For the crested cushion, we identified 176, 207, 203, 233, 296, and 401 DEGs in each developmental stage, respectively. Based on the KEGG enrichment analysis, we found that almost all DEGs were enriched in fatty acid biosynthesis and metabolism, tumor formation and anti-tumor response, tissue repair, and neural cell development pathways. Furthermore, the positive selection genes, such as connective tissue growth factor (*CTGF*), fatty acid synthase (*FASN*), homeobox D10 (*HOXD10*), syndecan 3 (*SDC3*), and four and a half LIM domains 2 (*FHL2*), were DEGs between the crest region and adjacent frontal skin tissues, which were enriched in osteoclast differentiation, cell adhesion molecules, fatty acid biosynthesis and metabolism, and microRNA in cancer pathways. Overall, positive selection analysis and gene expression results suggested that crest cushion formation was largely related to neural cells, skin tissue, bone, and fatty tissue.

### Genomic signatures reveal the domestication of CC ducks

To identify genomic regions influenced by the domestication of CC ducks, we compared the genomes of MDZ and CC duck populations representing different geographic regions using the cross-population extended haplotype homozygosity (XP-EHH) [29]. We identified 1,151 (XP-EHH score > 4.409, Z-test *P* < 0.01) putative selective sweeps in the CC duck genome compared to MDZ (Figure 4a). To further identify genome-wide signatures of domestication selection, we calculated the fixation index (*F*_ST_) values between MDZ and CC ducks. In total, we identified 919 putative selective sweeps of CC ducks compared to MDZ (*F*_ST_ > 0.504, top 0.01) (Figure 4b). As genomic regions targeted by artificial selection may be expected to have decreased levels of genetic variation, we also measured and plotted nucleotide diversity (π) along their genomes. Selecting the windows with the top 1% diversity ratios, i.e., low diversity in the two mallard ducks but high diversity in the CC ducks, we found 1,023 potential artificial selection windows by the CC duck compared to the MDZ (Figure 4c). Combining the results of the three methods (*F*_ST,_ π, and XP-EHH), we obtained 51 putative selective regions covering 30 genes in the CC duck domestication process (Table S19). Among these genes, we found that dynamin 3 (*DNM3*), which is an activator of p53, was under selection. *DNM3* is a member of the dynamin family, which possesses mechanochemical properties involved in actin-membrane processes, is predominantly expressed in the brain, and is associated with Sézary’s syndrome, which is a lymphoproliferative disorder. Nanog homeobox (*NANOG*), a gene under positive selection in CC ducks, is a key factor in the specification of early embryonic pluripotent cells. If *NANOG* is ablated *in vivo*, it will directly affect the fate determination of embryonic stem cells. In addition, a previous study suggested that *NANOG* inhibits apoptosis and promotes cell cycle arrest mainly *via* p53 regulation. In addition, transient receptor potential cation channel subfamily V member 5 (*TRPV5*), which is the key gene regulating the homeostatic balance of calcium, is also under positive selection in CC ducks. The function of these genes under positive selection during CC duck domestication suggests that regulatory elements may also play a role in the GCR of the crested trait formation.

**Figure 4.**
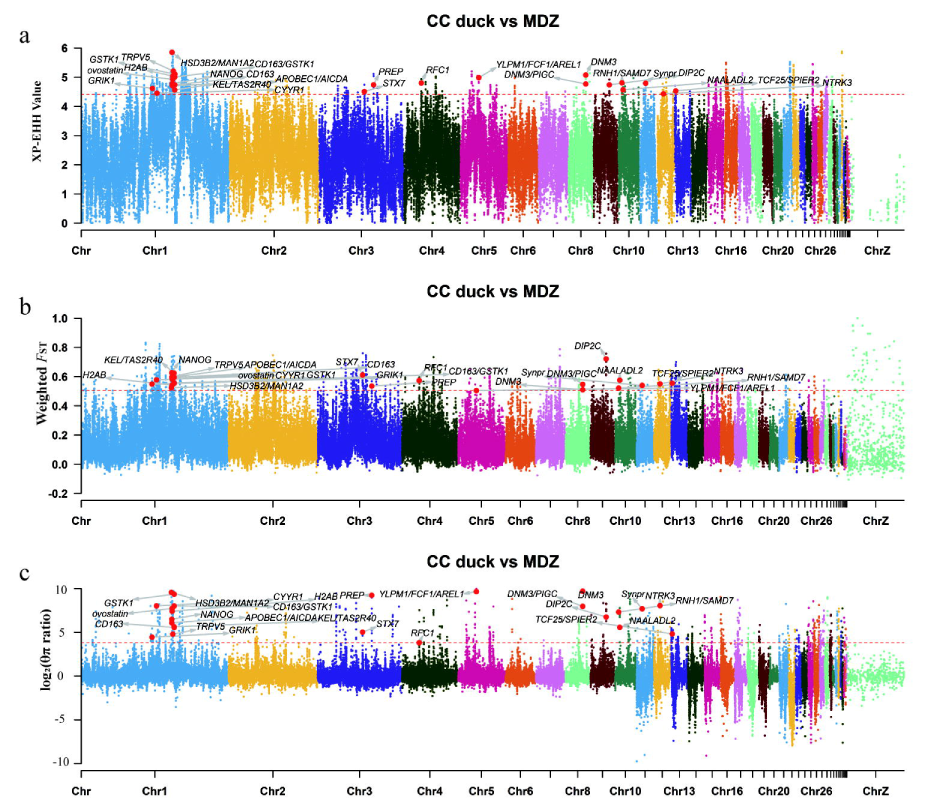
Genomic selection signatures during CC duck domestication. Distribution of cross-population extended haplotype homozygosity (XP-EHH) test (a), population differentiation (*F*_ST_) (b), and log_2_(θπ) ratio (c) between CC duck and MDZ duck using a 20 kb sliding window and 10 kb step; the dotted line represents the significant threshold (*F*_ST_: top 1%; XP-EHH value: Z-test *P* < 0.01; and log_2_(θπ) ratio).

### Structural variation detection reveals the essentials of genome adaptive evolution and genetic compensation

Genome-level evolution and structural variant accumulation provide an impetus for the adaptive genome evolution of species. Genomic structural variation can have a pronounced phenotypic impact, disrupting gene function and modifying gene dosage, whereas some large structural variations can lead to large body mutations, including neurodevelopmental disorders and unique trait formation. The CC duck has specific phenotypic traits in the crest cushion and immune levels compared to the Pekin duck and Csp-b ducks. To explore the reasons for these differences at the genomic level, we used the same approach as that of Li et al. [30]. We identified 9,369 structural variants (SVs), including 1,935 insertions, 4,118 deletions, and 3,316 inversions in the CC duck assembly. These SVs correspond to 71.91% (6,737/9,369) of the previous SVs from Illumina short-read genome sequencing, and 28.09% (2,632/9,369) of the SVs were novel. We found 1,541 species-specific genes to be embedded or almost completely contained (>50% overlap of gene length) in the missing sequences of the Pekin duck assembly. We explored functional enrichment for the SVs and species-specific genes from CC ducks using the ClusterProfiler package of R v4.1.0 packages[31] and the GO terms revealed by the clustering tool REVIGO (Figure S6–S8) [32]. We identified 35 GO terms that were significantly overrepresented (false discovery rate (FDR) < 0.05) in more than one gene (Table S20). Notably, there were some GO terms related to tissue repair, including cell adhesion, homophilic cell adhesion, and cell communication. These functions may be related to the unique crest traits of the CC ducks. The candidate genes contained several genes related to the immune system and signal transduction, which may have played important roles in the sub-phenotype of the crest trait in CC duck, including: ephrin type-A receptor 1 (*EPHA1*), a key factor required for angiogenesis and regulating cell proliferation; RUNX family transcription factor 2 (*RUNX2*), mutations of which have been found to be associated with the bone development disorder cleidocranial dysplasia; and taste 2 receptor member 40 (*TAS2R40*), which plays a role in the perception of bitterness. Interestingly, some gene families related to animal domestication have appeared as structural variants, such as the SLC superfamily of solute carriers and taste 2 receptors.

Similarly, we also identified putative SVs in the Csp-b duck genome assembly by comparison with the CC duck genome, and identified 2,694 insertions, 3,991 deletions, 609 inversions, and 421 species-specific genes. Functional enrichment among these SV-related genes and species-specific genes from Csp-b ducks was determined using GO analysis and pathway analysis (Table S21). These 74 functional categories were statistical significant (*P* < 0.05), and the regulation of small GTPase-mediated signal transduction was ranked as the top category in the GO biological process. We also calculated *K_a_/K_s_* ratios by comparing Csp-b ducks to chicken (Figure S9) and Zhedong goose (*A. cygnoides*) (Figure S10) lineages to account for rapid genome evolution. We found that genes with elevated *K_a_/K_s_* values in Csp-b ducks were significantly enriched for these functions (FDR, *q* < 0.01). Furthermore, these functional GO terms overlapped with the SV-related GO terms by 20.27% (15/74) in Csp-b duck-Zhedong goose pairs and 12.16% (9/74) in Csp-b duck-gallus pairs. We further examined the overlapping GO terms for both pairs, and there were seven categories associated with energy metabolism, the nervous system, and signal transduction in the Csp-b duck. We speculate that these seven functional categories might contribute to the duck habitat environment–adapted phenotype.

### The physiological and genetic basis of crest traits

The crest, which is an interesting phenotypic trait, appears in most bird species worldwide. However, CC ducks are unique duck breeds with bulbous feathers and skin protuberances in China. To fully reveal the physiological basis of crest cushion formation, we investigated the development of the parieto-occipital region of the CC duck during embryo development by microscopy. The results showed a protuberance at the cranial crest of the E4 duck embryo (Figure 5h and Figure S11). Therefore, we speculated that epidermal hyperplasia generated pressure on the adjacent skull cartilage tissue in the fontanelle during the development period, which led to the appearance of perforations in the parieto-occipital region during the cartilage ossification process (Figure S12). Coincidentally, preadipocytes began to differentiate into fat cells. To compensate for the decrease in brain pressure caused by the perforation, different volumes of fat were deposited between the brain and cerebellum (Figure S13). However, spherical feathers are only used to protect against fragile epidermal hyperplasia. Thus, the formation of the crest cushion is the result of several consecutive coincidences during the development of the skull, scalp, and feathers. Protuberance may be the most fundamental cause of crest formation, and the sub-phenotype of the crested trait was therefore attributed to phenotypic compensation in response to the crested cushion. Furthermore, we found that the inheritance patterns of the crest trait conformed to Mendel’s genetic laws in the F_2_ generation of 707□CC ducks ×□CV ducks (crest: crestless□=□541:166, χ^2^_df = 1_□=□0.35).

**Figure 5.**
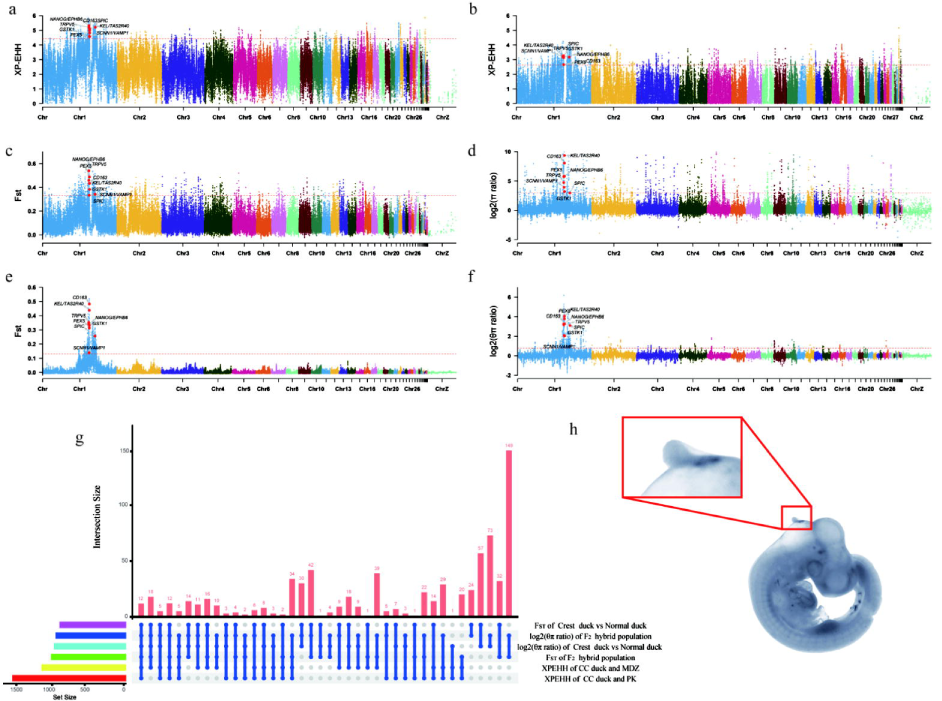
Selective-sweep analysis of the crest cushion of the CC duck. XP-EHH values for CC duck compared to MDZ (a) and MDN (b). c–d) Manhattan plot of *F*_ST_ and log_2_( ) ratio of CC duck domestication. e–f) Manhattan plot of *F*_ST_ and log_2_(θπ) ratio for selection of crest duck in F_2_ hybrid. g) Upset plot representing overlaps between above selective analysis methods. h) The embryo of CC duck and a full image of the crest cushion.

To identify the genetic basis of crest traits, we performed genome-wide selection tests in CC ducks compared to Pekin ducks and MDZ, which represent phenotypes for several traits that are relevant for the crest trait of CC ducks. We calculated the global XP-EHH among the CC duck compared to the Pekin duck and MDZ using a 20 kb sliding window and a shift of 10 kb across the CC duck genome, and 1,561 and 1,156 putatively selected genomic regions were identified (Figure 5a–b). Additionally, we identified 289 selected regions shared by the two-pair comparison group. In an additional analysis involving the *F*_ST_ and log_2_(θπ) ratio using 12 crested ducks and 27 normal ducks, we identified 902 and 980 selective regions, respectively (Figure 5c–d). Combining the results of the selective sweep analysis of the above four methods, we identified 26 shared selective regions and spanned 18 candidate genes that we speculated to be associated with crest traits (Table S22). Additional *F*_ST_ and genetic diversity analysis of the F_2_ hybrid population identified 1,165 and 997 special selection regions with the top 1% global *F*_ST_ (Figure 5e) and log_2_(θπ) ratio values (Figure 5f). Combining the results of between- and within-population selective sweep analysis, we identified 12 selective windows that may be significantly related to the crested trait (Figure 5g and Table S23). Annotation of the 13 genes putatively influenced by the crested trait revealed functions associated with the sub-phenotypic crested trait.

To fine-map regions identified using selective sweep methodologies and search for direct evidence of genotype-phenotype associations, we performed genome-wide association analysis (GWAS) for crest traits with informative phenotypic records. Using a panel of samples from the F_2_ hybrid from high-quality SNPs as well as the mixed model, which involved a variance component approach to correct the population structure, we identified two significant signals (harboring 4,914 SNPs) that were associated with the crest cushion trait with a threshold of −log_10_(*P*-value)□=□8.38 (Figure 6a–b). SNPs in the 12 candidate divergent regions (CDRs) associated with crest cushion formation showed extensive linkage disequilibrium (LD). The peak position was located between the *KEL* and *TAS2R40* genes. Furthermore, we found that the genotype frequencies of the related sites in *TAS2R40* and *NANOG* almost separated the crested ducks and normal ducks in the F_2_ population (Figure S14). Therefore, we consider that *TAS2R40*, *KEL*, and *NANOG* might be candidate genes for crest cushions based on the selective sweep and GWAS co-localization criteria (Figure 6c–e). To detect the candidate SNPs, we used Sanger sequencing and genotyping of 30 CC ducks and 75 normal ducks from three duck breeds. We found that the genotype of the 5’UTR of *TAS2R40* (123272114_c. G78A) (Figure 6f), the first intron (123248845_c. G7127C) of *KEL* (Figure S15), and the fourth exon (120130992_c. G577A_p. V193M and 120131265_c. G850T: p. A284S) of *NANOG* (Figure S16) could separate the CC duck from other 15 non-crested ducks. Among them, only the 123272114_c. G78A of *TAS2R40* showed a 100% heterozygous genotype and homozygous mutant gene frequency, while other SNP loci showed a percentage of more than 80%. In particular, this SNP exhibited a significant *P*-value and could account for 54.68% of the explained phenotypic variation in MLM. Importantly, the tissue expression profile of *TAS2R40* at 56 days of age showed that *TAS2R40* was hardly expressed in the cerebellum, thigh muscle, and breast muscle. The relative expression in the crested cushion and abdominal fat was significantly higher than that in other tissues (*P* < 0.01) (Figure 6g). The results revealed that the mutation in the 5’UTR of *TAS2R40* specifically affected the expression level of *TAS2R40* in the crested tissue of CC ducks. Subsequently, luciferase assays showed that the relative luciferase activity of *TAS2R40* 5’UTR-MT was significantly lower than that of *TAS2R40* 5’UTR-WT (*P* < 0.01) (Figure 6h). A series of results showed that the G > A mutation in the transcription region affected the regulatory effect and reduced its expression in the crested tissue. Combining the above results, we speculated that the SNP in the 5’UTR of *TAS2R40* was a causative mutation of the crested cushion.

**Figure 6.**
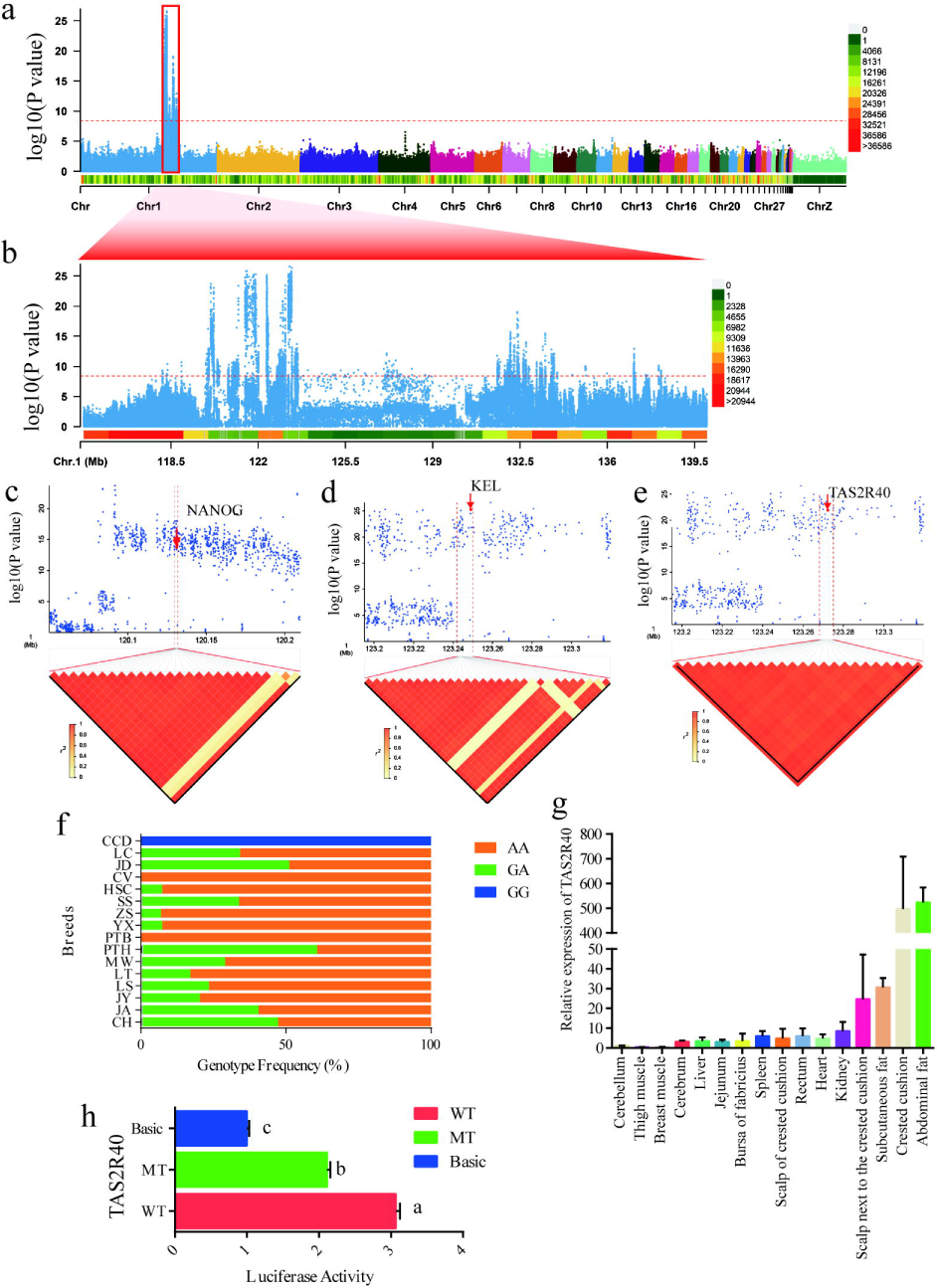
Candidate genes identified for the crest cushion in crested ducks. a) GWAS of the crest cushion, including 63 crested ducks and 211 normal ducks. The red horizontal dashed lines indicate that the Bonferroni significance threshold of the GWAS was 8.39 (0.05/total SNPs). b) Magnified view of the chromosome 1 peak in (a). c–e) Zoom of 0.2 Mb and 0.1 Mb of candidate SNPs for *NANOG*, *KEL*, and *TAS2R40*. The LD heatmap (d) is depicted in red. f) Genotype frequency of *TAS2R40* (123272114_c. G78A). The CCD represent the Chinese creste duck; LC represent the Lianchen duck; JD represent the Jingding duck; HSC represent the Taiwanese Brown Vegetable Duck; SS represent the Sansui duck; ZS represent the Zhongshanma duck; YX represent the Youxianma duck; PTB represent the Putian white duck; PTH represent the Putian black duck; MW represent the Mawang duck; LT represent the mallard; LS represent the Longshencui duck; JY represent the Jingyunma duck; JA represent the Ji’an read feather duck and CH represent the Chaohu duck. g) Relative expression of *TAS2R40* gene in 16 tissues of the CC duck. h) The luciferase activities were detected after transfection of *TAS2R40* 5’UTR-MT (mutant type) vector and *TAS2R40* 5’UTR-WT (wild type) vector. Statistical significance given as different lowercase letters.

## Discussion

Since the first duck draft genome was published in *Nature Genetics* in 2013 [13], the origin, evolution, domestication, and selection of ducks have been revealed. More importantly, a series of characteristic traits and phenotypes, such as disease resistance, body size, plumage color, and egg color have been gradually discovered [12, 33], providing deep insights into genotype-phenotype associations in animal molecular breeding and germplasm conservation.

During species evolution, directional artificial selection and non-directional natural selection can cause genetic diversity in animals. Adaptive evolution allows animals to acquire certain protective mechanisms that allow the species to continue. Based on the phenotype analysis, we explained the mechanism of crest cushion occurrence at an anatomical level and found that the crest cushion might affect the survival of the CC duck. Theoretically, natural selection promotes the spread of mutations and removes harmful ones. However, it is not completely effective, and all populations harbor genetic variants with deleterious effects. Human intervention in speciation preservation might maintain the inheritance of harmful mutations and promote the accumulation of beneficial mutations. The results presented herein provide evidence of human intervention leading to genome protection and the evolutionary maintenance of species. Considering the structural variants, genome evolution-related genes, and gene content enrichment among various birds, there is evidence for genome protection and evolutionary maintenance of species that complement one another. The CC ducks had a greater proportion of genes under adaptive evolution with functions related to tissue repair than the other two ducks.

Crest cushions or crest crowns are conspicuous and diverse features of almost all bird lineages with feather crests and are unique among almost all bird species [34]. The most obvious difference between the Chinese crested duck and other existing crested birds is that the crested tissue of the crested duck affects the embryonic development of the crested duck and can even lead to embryonic death. Our results indicate that the crest cushion is caused by the proliferation of relevant cells in the parieto-occipital region during the embryonic stage. This process generates downward pressure, resulting in incomplete closure of the cartilage and, in some crested ducks, likely leading to brain overflow and death as a result of exaggerated crest cushion size. This finding demonstrates that the root cause of crested cushion formation is the rapid proliferation of cells in the parieto-occipital region. Furthermore, we observed that even if some crested duck embryos have a hole in the cartilage, the mortality rate of crested ducks is very low if the scalp heals and adipose tissue compensate for the insufficient brain pressure (Figure 7). Based on the above results, we propose that the healing of the scalp and the presence of adipose tissue may act as a phenotypic compensation mechanism for crested tissue to reduce the mortality of crested ducks. To reveal the protein basis, we generated a high-quality chromosome-level CC duck genome. Compared to the recently reported duck genome[12–14, 33], the evaluation result of BUSSCO was better than that of other duck genomes. Furthermore, we compared the CC duck genome to other bird genomes and identified some genes related to tumorigenesis. Simultaneously, some immune-related genes in the crested duck genome have also undergone positive selection due to the presence of holes that can cause brain exposure, which is more important for CC duck survival. In addition, we believe that the composition of these phenotypes may be the physiological basis of crested formation under stronger positive conditions. We also identified the genetic basis of crest trait formation and phenotypic composition by inter- and intra-population selective sweep analysis. We found that 12 CDRs harboring 13 genes were strongly selected in CC ducks. Based on GWAS and experimental evidence, we confirmed that *TAS2R40* may be the most fundamental cause of mortality. We speculate that the 5’UTR mutation of *TAS2R40* may affect the expression of *TAS2R40*, leading to abnormal expansion of certain ectodermal cells in the early embryonic development stage, forming the initial protruding tissue of the crested head, leading to the occurrence of the crested trait. In addition, ephrin type-b receptor 2 (*EPHB2*), which belongs to the same gene family as ephrin type-b receptor 6 (*EPHB6*) identified here, has proven to be related to the inverse growth of the crest feathers of crested pigeons [35]. Thus, *EPHB6* may control cranial crest feathers to grow clockwise, forming a spherical crested feather phenotype in the crested duck. Thus, the crested duck can form a protective tissue on fragile crested tissue. NANOG, which is involved in the development of neural crest stem cells, has been shown to play a role in the pathogenesis of many cancers by regulating cell proliferation, invasion, and metastasis [36, 37]. Therefore, we suggest that *NANOG* could be a key gene in DNA damage repair and the GCR in CC ducks.

**Figure 7.**
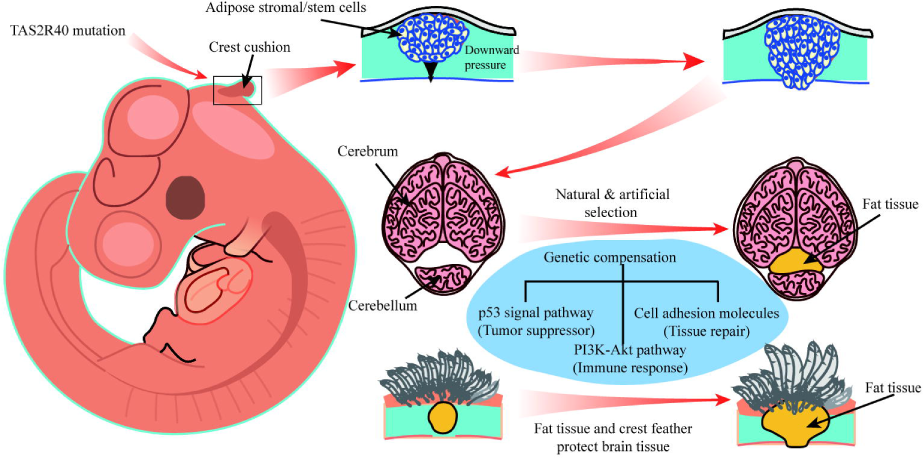
Diagram of crested cushion formation and genetics compensation of CC duck. The causative mutation of *TAS2R40* leads to the proliferation of cells in the parieto-occipital region during embryonic stage. As these cell continue to proliferate, downward pressure is generated on the chondrocyte layer in the parieto-occipital area, forming irregular holes, causing incomplete skull closure, and reducing intracranial pressure. Through a long process of natural and artificial selection, some phenotypic compensation has occurred in the crested tissue. For example, an adipose tissue is formed between the cerebrum and the cerebellum to supplement the lost intracranial pressure. Simultaneously, the scalp in the crested area gradually thickens, accompanied by the increase of crested tissue, the change of the polarity, and rapid growth of the top feathers, which protects the leaking brain tissue due to the presence of holes. The most fundamental reason for phenotypic compensation is the selection of pathways, such as cancer suppression (p53 pathway), immunity (PI3K-Akt pathway), and tissue repair (cell adhesion molecules) in the evolution process, which leads to genetic compensation.

Previous studies have shown that all domesticated ducks originate from mallard ducks [12, 33]. However, according to the distribution map of the mallard, we found that the mallard exists in two regions of China: the northern and southern group. However, our data suggest that the CC duck originated from mallards in Zhejiang Province, China, and provided important findings on the history of the CC duck. In recent decades, the CC duck has become endangered, but it has quickly recovered in response to conservation efforts. Our analyses identified 30 candidate genes in the genomic regions under selection in the CC duck domestication process, with most of these genes related to neuron development, response to stress, and response to wounding. Therefore, CC ducks represent a critical example of evolutionary adaptation and genetic compensation.

By comparing the CC duck genome with those of 13 other bird species, we shed new light on how CC ducks likely evolved *via* the GCR mechanism and propose this breed as a model for studying GCR by natural selection. We found that the four main biological processes were likely co-enriched. The first process involves tumorigenesis and suppression, such as the p53 pathway, PD-L1 expression and PD-1 checkpoint pathway, cellular response to DNA damage stimulus, etc. Based on our observations, we suggest that the root cause of crested head formation may be the short-term rapid proliferation of cranial neural crest cells (similar to local neoplasia). However, with the evolution of the CC duck, the crested duck has evolved a control system that can prevent cells from continuing to proliferate rapidly. Second, we enriched some pathways related to tissue repair, such as cell adhesion molecules and focal adhesions. These genes may control scalp and cartilage healing to prevent encephalocele formation. Third, we identified the genes related to fat synthesis and metabolism. We suspect that the main role of these genes in the brain is the formation of adipose tissue, which is used to compensate for the loss of missing skull intracranial pressure, thus ensuring that crested ducks maintain normal levels of brain pressure. Fourth, due to the existence of crested tissue, the immune system has also undergone a certain degree of positive selection, such as the immune-related genes enriched in the PI3K-Akt pathway (Figure 7). In short, the compensatory evolution of a series of genes caused by the occurrence of crested traits has allowed crested ducks to survive and even stabilize the population. Other genes may have evolved due to the accompanying mutations caused by crested traits, incurring a GCR and protecting the survival of crested ducks. However, the regulatory relationship of these genes in the mechanism of crest cushion formation remains unclear, and with advances in cell biology, this problem will be gradually solved in the future.

## Conclusions

In the present study, we revealed the genetic mechanisms underlying the evolutionary, developmental, and histological origins of the crest trait of CC ducks and provided insights into the molecular mechanisms of the GCR and its relevance to cancer resistance. The identified genes and their specific mutations provide a starting point for future functional studies of crest cushion development, genetic compensation mechanisms, oncogenesis, and tumor defense.

## Materials and methods

### Ethical approval

All experiments with ducks were performed in accordance with the Regulations on the Administration of Experimental Animals issued by the Ministry of Science and Technology (Beijing, China) in 1988 (last modified in 2001). The experimental protocols were approved by the Animal Care and Use Committee of Yangzhou University (YZUDWSY2017-11-07). All efforts were made to minimize animal discomfort and suffering.

### Sample preparation and sequencing

A 28-week-old female CC duck from Zhenjiang Tiancheng Agricultural Science and Technology (Zhenjiang, Jiangsu, China) was used for genome sequencing and assembly. High-quality genomic DNA was extracted from the blood tissue using a standard phenol/chloroform protocol [38]. A paired-end Illumina sequence library with an insert size of 350 bp and a 10× Genomics linked-read library was constructed and sequenced on the Illumina HiSeq X Ten platform (San Diego, CA, USA). A PacBio library was constructed and sequenced using the PacBio Sequel platform (Menlo Park, CA, USA). RNA-seq libraries for eight tissues (crested tissue, spleen, ovary, liver, duodenum, skin, pectoral, and blood) were constructed and sequenced using Illumina HiSeq4000. Clean reads were assembled using Trinity for gene prediction. In addition, a 28-week-old female Csp-b duck was used for genome sequencing and assembly. Short-insert (250 bp and 350 bp) paired-end libraries and large-insert (2 kb and 5 kb) mate-pair libraries were constructed and sequenced on the Illumina HiSeq4000.

### Genome size estimation, assembly, and quality assessment

The genome size of the CC duck genome was estimated based on the *k*-mer distribution using high-quality paired-end reads. Contig assembly of CC duck was assembled with PacBio reads using FALCON v0.7 [12]. This assembly was polished using Quiver [13] with the default parameters. 10× Genomics was then used to connect contigs to super-scaffolds using fragScaff software [14]. Subsequently, Illumina paired-end reads were used to correct for any errors using Pilon v1.18 [39]. Finally, the scaffolds were anchored and oriented on chromosomes using CHROMONMER v1.07 [18]. A detailed description of the genetic linkage map construction and chromosome anchoring is presented in the supplementary materials and methods. To estimate the quality of the final assembly, short paired-end reads were aligned onto the CC duck genome using the Burrows-Wheeler aligner (BWA) with the parameters of ‘-k 32 -w 10 -B 3 -O 11 -E 4’. BUSCO [16] was used to assess completeness.

### Genome annotation

Homology-based and *de novo* predictions were combined to identify repetitive sequences in the CC duck genome. RepeatMasker and RepeatProteinMask (both available from http://www.repeatmasker.org) were used for homologous repeat detection to run against RepBase [40], LTR_FINDER [41], RepeatModeler and RepeatScout [42] were used to construct a *de novo* repeat library with default settings. Using the *de novo* library, RepeatMasker was run on the CC duck genome. Tandem repeats were identified using TRF v4.07b [43].

Gene prediction was performed using homology-based prediction, *ab initio* prediction, and transcriptome-based prediction. Protein sequences of *Anser cygnoides*, *Aptenodytes forsteri*, *Anas platyrhynchos domestica*, *Coturnix japonica*, *Columba livia*, *Egretta garzetta*, *Gallus gallus*, *Homo sapiens*, *Nestor notabilis*, *Struthio camelus*, and *Taeniopygia guttata* were aligned against the CC duck genome using TBLASTN [44]. The blast hits were then conjoined by Solar software [45] and GeneWise [46] was used to predict accurate spliced alignments. For *ab initio* prediction, Augustus [47], Genscan [48], Geneid [49], GlimmerHMM [50] and SNAP [51] were used to predict genes in the repeat-masked genome. RNA-seq data from eight tissues were aligned to the genome using Tophat and Cufflinks [52] to predict gene structures. All predicted genes from the three approaches were integrated using the EvidenceModeler (EVM) [53]. Functional annotation of the predicted genes was carried out using BLASTP against the public databases: To obtain gene functional annotations, KEGG [54], SwissProt [55] and NR databases [56]. InterProScan [57] was used to identify domains by searching the InterPro and GO [58] databases. *Comparative genomic analyses*

In total, 14 species, including *Anser cygnoides domesticus*, *Aptenodytes forsteri*, *Balearica regulorum*, *Coturnix japonica*, *Columba livia*, CC duck, *Egretta garzetta*, *Gallus gallus*, *Gavia stellata*, *Nestor notabilis*, *Opisthocomus hoazin*, *Podiceps cristatus*, *Struthio camelus,* and *Taeniopygia guttata*, were used for gene family analysis. The longest transcripts of each gene (>30 amino acids) were retained when a gene had multiple splicing isoforms. ‘All-against-all’ BLAST v2.2.26 (e-value <= 1e-7) [44] was used to determine the similarities between the retained genes. OrthoMCL software [22] was used to define the orthologous groups in the above species with the parameter of ‘-inflation 1.5’. The phylogenetic tree was reconstructed using single-copy orthologs from gene family analysis. Multiple alignments were performed using MUSCLE [59]. The protein alignments were transformed back to CDS alignments, and then the alignments were concatenated to a super alignment matrix. We constructed a maximum-likelihood phylogenetic tree using RAxML [60]. The mcmctree program from PAML was used for divergence time estimation with eight calibration points from the TimeTree website [61], and the calibration points are provided in Table S24. We determined the expansion and contraction of orthologous gene families using CAFÉ v1.6 [62] based on a random birth and death model to model gene gain and loss over a phylogeny.

To identify PSGs, all single-copy gene families of five species, including CC duck, *Anser cygnoides domesticus*, *G. gallus*, *P. cristatus*, and *A. platyrhynchos*, were used for analysis. Protein-coding sequences were aligned with MUSCLE [63] and the branch-site model of CODEML in PAML was used to identify PSGs by setting the CC duck as the foreground branch. *P*-values were calculated using the chi-square test and corrected by the FDR method. Sequence quality and alignment errors have certain influences on the test, so the PSGs with low alignment quality were filtered using the following criteria: (1) FDR > 0.05; (2) presence of gaps near three amino acids around the positively selected sites in the five species. In addition, the kaka_calculator was used to calculate the *K*_a_/*K_s_* ratio [64].

### Structural variation detection

We built pairwise local genome alignments between the CC duck and two other duck genome assemblies (i.e., the Pekin and Csp-b duck) using LASTZ v1.04.00 [39] with the parameters of ‘*M* = 254, *K* = 4,500, *L* = 3,000, *Y* = 15,000, *E* = 150, *H* = 2,000, *O* = 600, and *T* = 2’. The genomes used for pairwise alignments were soft-masked for repeats using the RepeatMasker software. Then we used “axtChain” to build the co-linear alignment chains and used “chainNet” to to obtain nets from a set of chains with the default parameters. The “netSyntenic” command was used to add the co-linear information to the nets. The “netToAxt” and “axtSort” were used to convert the net-format to axt-format and change the order of the sequences, respectively. Subsequently, we obtained the best hit for each single location by the utility “axtBest” [65].

Structural variant detection was performed based on the best alignment hits with gapped extension, which indicated insertions or deletions. In addition, short paired-end reads of the Pekin and Csp-b duck genomes were aligned onto the CC duck genome by BWA [42]. Based on the depth of the reads, we validated our structural variation results. Deletions with average depth less than half of the average depth of the whole reference genome, and insertions with average depth over half of the average depth of the whole assembly. The software source code is available from Li et al. [26].

### RNA sequencing and transcriptomic analysis

Total RNA was extracted from the crest region and adjacent frontal skin tissues of CC ducks and scalps of Cherry Valley ducks using RNAiso Plus reagent (code no. 9109; Takara, Dalian, China) according to the manufacturer’s instructions, and 3 µg per sample was used as the input material for RNA sample preparations. The PCR products were purified using an AMPure XP system, and library quality was assessed using an Agilent Bioanalyzer 2100 system. After cluster generation, the library was sequenced using an Illumina HiSeq platform at Novogene Biotechnology (Beijing, China), and 125/150 bp paired-end reads were generated. The quality of the RNA sequences was checked using FastQC, while sequence adapters and low-quality reads (read quality <[30) were removed using Trimmomatic v0.36 with TRAILING:20 and SLIDING WINDOW: 4:15 as parameters. The remaining high-quality RNA-seq clean reads were aligned to the corresponding CC duck genome using HISAT2 v2.1.0 with default parameters. FeatureCounts v1.5.0-p3 (parameters: -Q 10 -B -C) was used to count the transcript reads, and StringTie was used to quantify the gene expression levels (in fragments per kilobase of transcript per million mapped reads; FPKM) in the detected tissue based on the corresponding transcript annotation. DEGs were identified using negative binomial generalized linear models implemented in DESeq2 v1.20.0. Genes with a *P* <[0.05, and |log2 (fold change (FC)) |[≥ 1 were considered DEGs. Hierarchical clustering analysis was performed to determine the variability and repeatability of the samples, and a volcano plot was used to visualize the DEG distribution.

### Historical population size estimation

The recent demographic history was inferred from the trends in the *N_e_* changes using PopSizeABC v2.1 [66] with default parameters set for the duck population (mutation rate of 7.54 × 10^-8^ and recombination rate of 1.6 × 10^-8^, minor allele count threshold for allele frequency spectrum (AFS) and identity-by-state (IBS) statistics computation = 4, minor allele count threshold for LD statistics computation = 4, and size of each segment = 2,000,000) and 1,000 simulated datasets. Summary statistics were extracted using the same parameters, with the tolerance set to 0.05, as recommended.

### Alignment and variation calling

A total of 308 samples from GWAS were aligned to the CC duck genome using BWA [67] (settings: mem -t 4 -k 32 -M -R). The sample alignment rates were between 96–98.00%. The average coverage depth for the reference genome (excluding the *N* region) was between 9.34–15.74×, and 4X base coverage (≥4) was greater than 82.64%. All the population structure analysis samples were aligned to the CC duck genome using BWA (settings: mem -t 4 -k 32 -M -R), and the sample alignment rate was between 94–98.42%. The average coverage depth for the reference genome (excluding the *N* region) was 6.00X and 17.66X. Variant calling was performed for all samples using the Genome Analysis Toolkit (GATK) v 3.7 [68] with the UnifiedGenotyper method. The SNPs were filtered using Perl script. After filtering, the GWAS sample retained 12.6 Mb of SNPs (filter conditions: only two alleles; single-sample quality = 5; single-sample depth: 5∼75; total-sample quality = 20; total-sample depth: 308∼1,000,000; maximum missing rate□of individuals and site = 0.1; and a minor allele frequency = 0.05), and the population genetics analysis retained 5.4 Mb of SNPs (filter conditions: only two alleles; single-sample quality = 5; single-sample depth: 3∼75; total-sample quality = 20; total-sample depth: 39∼1,000,000; maximum missing rate□of individuals and site = 0.1; and a minor allele frequency = 0.05).

### Population structure analysis

To clarify the phylogenetic relationship from a genome-wide perspective, an individual-based NJ tree was constructed based on the p-distance using TreeBeST v1.9.2 [69]; the bootstrap value parameter was 1,000. PCA was performed based on all the SNPs using GCTA v1.24.2 [70]. The population genetics structure was examined using an expectation maximization algorithm, as implemented in the program FRAPPE v1.1 [21]. In the population genetics structure analysis, we filtered 5,425,458 SNPs from 36,611,493 SNPs that were filtered by GATK (filter conditions: minor allele frequency = 0.05, maximum missing rate[of individuals and site = 0.1, single-sample depth = 3, and single sample quality = 5). The number of assumed genetic clusters *K* ranged from two to seven, with 10,000 iterations for each run. We compared the patterns of LD using high-quality SNPs. To estimate LD decay, the degree of the LD coefficient (r^2^) between pairwise SNPs was calculated using Haploview v4.2, and R v4.1.0 was used to plot LD decay [71]. The program parameters were set as ‘-n -dprime -minMAF 0.05.’ The average r^2^ value was calculated for pairwise markers in a 500-kb window and averaged across the whole genome. We found differences in the rate of decay and level of the LD value that reflected variations in the population demographic history and *Ne* among breeds/populations.

We estimated the ancestry of each individual using the genome-wide unlinked SNP dataset, and the model-based assignment software FRAPPE [21] was used to quantify the genome-wide admixture between the wild duck, Pekin, and CC duck populations. FRAPPE was run for each possible group number (*K* = 2 to 4) with default parameters to estimate the parameter standard errors used to determine the optimal group number (*K*).

### Selective-sweep analysis

To identify the putative selective sweep regions, we used the high *F*_ST_ value [72], high differences in genetic diversity (π log2 ratio), and XP-EHH [29]. We calculated the *F*_ST_ and π log2 ratio value in 20-kb sliding windows with 10-kb steps along the autosomes using VCFtools [73] for further analyses, where *F*_ST_ was employed for comparisons among the CC, Pekin, and Csp-b ducks were used for comparisons between crested and normal ducks in the F_2_ population. We then filtered out any windows that had fewer than 20 SNPs. The top 1% of windows or regions with the highest reduction in nucleotide diversity (ROD) values represented the extreme ends of the distributions.

### Genome-wide association study

A case-control GWAS was conducted, including 63 crested ducks (case) and 211 normal ducks (control), involving a total of 12.6 Mb of SNPs. After filtering with PLINK v1.90 [74] (“--geno 0.1 --hwe 1e-05 --maf 0.05 --mind 0.1”), 63 crested ducks and 211 normal ducks with a total of 12.2 Mb of SNPs were used for the subsequent association study. An MLM program, Efficient Mixed-Model Association eXpedited (EMMAX) (beta version) [75], was used to carry out the GWAS. To minimize false positives, the population structure was considered using the top 20 PCA, which was estimated using PLINK. For the F_2_ population, the top 20 PCA values (eigenvectors) were set as fixed effects in the mixed model. The BN kinship matrix was set as a random effect to control for family effects. We defined the whole-genome significance cutoff as the Bonferroni test threshold, which was set as 0.05/total effective SNPs. The GWAS threshold for the crest-cushion was 8.38. Manhattan plots and QQ plots of GWAS were produced using the qqman package in R v4.1.0 [76].

### GO and pathway enrichment analyses using DAVID and clusterprofiler

The Database for Annotation, Visualization, and Integrated Discovery (DAVID) v6.8 (https://david.ncifcrf.gov) [77] and clusterprofiler[31] were used to perform GO enrichment and KEGG pathway analyses. The Bonferroni method, which is a method of R/stats package, was used to adjust the *P*-value in GO enrichment and KEGG pathway analysis.

### Experimental validation

PCR was performed to determine the candidate genes. Oligo 6 was used for primer design, and the primers and annealing are shown in Supplementary Table S25. Primers for RT-qPCR (Table S25) were designed using the Oligo 6. *TAS2R40* was used to measure the expression levels. Three CC ducks of 56 days of age were slaughtered by stunning and exsanguination. Tissue samples, including the cerebellum, thigh muscle, breast muscle, cerebrum, liver, jejunum, bursa of Fabricius, spleen, scalp of crested cushion, rectum, heart, kidney, scalp next to the crested cushion, subcutaneous fat, crested cushion, and abdominal fat (50–100 mg) were rapidly collected, snap-frozen in liquid nitrogen, and stored at –80 °C. Glyceraldehyde 3-phosphate dehydrogenase (*GAPDH*) was used as an endogenous control. The collected data were analyzed using the 2^-ΔΔCt^ method. Fragments of the 5’UTR of *TAS2R40* were cloned and inserted between the NheI and XhoI restriction sites of the pGL-Basic 3.0 vector. Luciferase activity was measured 36 h after transfection using the dual-luciferase reporter system (Promega, Madison, WI, USA). Firefly luciferase activity was normalized to Renilla luciferase activity.

## Data availability

The genome assembly and all of the re-sequencing data were deposited in the BIG Data Center (http://bigd.big.ac.cn/) under BioProject accession PRJCA001785.

## Author’s statement

GBC and GHC conceived the project and designed the study. QXG, ML, XFC, HB, and HL performed the bioinformatics analysis. GBC, GHC, XYY, and SSW constructed the F_2_ population. GBC, XYY, SSW, ZXW, QX, YZ, QQS, RP, SHZ, LLQ, TTG, XSW, YLB, ZFC, YZ, YC, and WCD collected the F_2_ population phenotype data. XYY, HB, and YJ prepared the DNA and RNA, and performed laboratory experiments. GHC, GBC, XYY, QXG, ML, XFC, and HB wrote the manuscript. BCL, ZQW, and JFL have revised the manuscript. All authors read and approved the final manuscript.

## Competing interests

The authors have declared no competing interests.

## Acknowledgments

This project was supported by the China Agriculture Research System (CARS-42), the Jiangsu Agricultural Technology System (JATS[2020]435), and the Jiangsu Agricultural Science and Technology Innovation Fund (CX[18]1004). We are deeply grateful to all the donors who participated in this program. We thank Prof. Lizhi Lu from Zhejiang Academy of Agricultural Sciences and Prof. Lujiang Qu from China Agricultural University for providing the NGS data of mallard duck, Prof. Zhuocheng Hou from China Agricultural University for providing the genome annotation file of mallard duck; Prof. Yunzeng Zhang and Prof. Duonan Yu from Yangzhou University for their suggestions on the data analyses and manuscript writing, and Prof. Zhiqiang Du from Northeast China Agricultural University for helpful suggestions on the F_2_ population design.

