## Supplementary material for "The first crested duck genome reveals clues to genetic compensation and crest cushion formation": Supplementary materials and methods.docx

Supplementary Information

Guobin Chang1,3,†, Xiaoya Yuan1,†，Qixin Guo1,†，Hao Bai3,†, Xiaofang Cao2,†, Meng Liu2,†, Zhixiu Wang1, Bichun Li1, Shasha Wang1, Yong Jiang1, Zhiquan Wang4, Yang Zhang1, Qi Xu1, Qianqian Song1, Rui Pan1, Shenghan Zheng1, Lingling Qiu1, Tiantian Gu1, Xinsheng Wu1, Yulin Bi1, Zhengfeng Cao1, Yu Zhang1, Yang Chen1, Hong Li2, Jianfeng Liu5, Wangcheng Dai6, and Guohong Chen1,3,*

^1^Key Laboratory of Animal Genetics and Breeding and Molecular Design of Jiangsu Province, College of Animal Science and Technology, Yangzhou University, Yangzhou, China

^2^Novogene Bioinformatics Institute, Beijing, China

^3^Joint International Research Laboratory of Agriculture and Agri-Product Safety, the Ministry of Education of China, Institutes of Agricultural Science and Technology Development, Yangzhou University, Yangzhou, China

^4^Department of Agricultural, Food, and Nutritional Sciences, University of Alberta, Edmonton, AB, Canada

^5^College of Animal Science and Technology, China Agricultural University, Beijing, China

^6^Zhenjiang Tiancheng Agricultural Science and Technology Co.,Ltd, Zhenjiang, Jiangsu, China

†These authors contributed equally to this work.

*Correspondence:

**Content:**

**Supplementary Materials and Methods**

**1. Samples information**

For the genome assembly and annotation, the genomic DNA (gDNA) of a 28-week-old female Chinese crested (CC) duck and a 28-week-old female Chinese spotted-bill (Csp-b) duck was extracted from whole blood and isolated using a traditional phenol-chloroform protocol. The CC duck came from our experimental base in Zhenjiang Tiancheng Agricultural Science and Technology Co., Ltd. (Zhenjiang, Jiangsu, China), and the Csp-b duck was obtained from Jiangsu Yancheng Wetland National Nature Reserve, Rare Birds in China. Additionally, we sampled a total of five duck breeds, including two populations of domesticated ducks and three populations of mallards from different geographic regions. The domesticated ducks were one crested duck population, i.e., CC duck (n = 12) and other domesticated population, i.e., Pekin duck (PK duck; n = 8, download from NCBI, BioProject accession number: PRJNA419832, BioSample ID: SAMN08099603-SAMN08099610). The mallard ducks were three wild breeds duck, i.e., mallards from Zhejiang Province (MDZ; n = 10), Ningxia Province (MDN; n = 8, download from NCBI, BioProject accession number: PRJNA419832, BioSample ID: SAMN08099581-SAMN08099588) and Csp-b duck from Jiangsu Province (n = 1). In addition, a goose breed (Anser cygnoides domesticus, Acyg, BioProject accession number: PRJNA183603, BioSample ID: SAMN01830643) was used for phylogenetic analysis in the present study. A complete description of the appearance, characteristics, and sample distributions of the breeds in this study is presented in Table S21.

**2. Crest phenotype observation**

To analyze the occurrence and composition of crest traits, magnetic resonance imaging (MRI) equipment (Discovery MR750w3.0T) was used to detect the morphological and histological influence on crest trait in CC duck. To help analyze the crested trait, we used intramuscular (IM) anesthetics (Phenobarbitol sodium) to help for fixation the CC duck. And the special coil performed to improve the signal to noise ratio. For examination, the anaesthetized ducks were limited in ventral recumbency in a plastic pan and put in the middle of the magnetic field on a sliding carriage. The ducks were imaged in a 3.0 Tesla whole-body magnet. During the measurement, a multi-slice sequence of measuring blocks was generated, including 15 sagittal adjacent slices of 1.8 mm thickness. A repetition time of 1,927.7 ms for sagittal images and an echo time of 29.2 ms was set. And, the result of MRI shown that the relate tissue of crest cushion extends from the crested tissue to between the brain and the cerebellum (Figure S13). And then, with a skull specimen to observed the skull change of CC duck compare to normal duck. To make the skull specimen, large muscles were peeled away and removed, and the remaining muscles and fats were treated with a 10% neutral protease solution in a constant temperature water bath at 70 °C for 24 hours. The skull specimen was dried and fixed with quick-drying glue. Base on the skull specimen, we found a perforation on the region of posterior fontanelle. And crested ducks with different sizes of the crested cushion have different size perforations (Figure S12). Moreover, to further explore the histological formation mechanism of the crested trait, we performed continuous microscopic (Figure S11), anatomical and sections of crested raised tissue to observations of duck embryo development and analysis the sub-phenotypic of the crest trait.

**3. Genetics linkage mapping population establishing**

To aid in genome assembly, a full-sib family was established by mating a pair of a CC duck and a Cherry valley (CV) duck which showed a large genetic distance. The CC duck is a small-sized breed with high feather crests, a black beak and shank, and white plumage. In contrast, the CV duck is a large-sized breed with a yellow bill and white plumage but no crested phenotype on the cranial crest. In the F_1_ generation, we collected more than 20 ducks, and the ratio of male to female ducks was relatively balanced. Interestingly, no crested duck appeared in the F_1_ generation, and the phenotype of F_1_ individuals was uniform in all families. Although genetic performances differed within the generation of reciprocal crosses to a certain extent, the F_1_ generation was generally heterozygous and consistent. The F_2_ generation was produced from the natural mating of F_1_ hybrids. In the F_2_ generation, we collected and sampled the venous blood of 177 ducks through the sub-wing vein blood collection method for subsequent resequencing.

**4. F_2_ hybrid population construction**

To determine candidate loci or genes of several traits of interest, we constructed an F_2_ segregating population in the present study. The F_1_ generation was produced from orthogonal crosses of CC duck and CV duck. In the orthogonal experiment, 86 CC ducks and 13 CV ducks were randomly selected to be divided into seven families. The offspring number in the F_1_ generation reached more than 500 individuals. Moreover, there is no duck with crest trait in the F_1_ generation. The ratio of male to female ducks was relatively consistent. The F_2_ generation was produced from the natural mating of F_1_ hybrids, and the mating was internally limited to the orthogonal experiments. In building the families, we considered and complied with the following principles: (1) the ratio of males to females was 1:3, (2) males and females in the same family were not from the same nest, and (3) female ducks within a family were not half-siblings. To avoid half-siblings, we designed a mating system. The F_2_ generation was composed of almost 2000 ducks that displayed segregation of various genetic characteristics, including the crest cushion, feather color, bill color, shank color and body weight. When the ducklings hatched, the ducks were weighed each week after hatching. At three weeks after hatching, all members of the F_2_ generation were moved from the duckling house to a designed individual shed and were raised to the age of six weeks. We performed a slaughter experiment of more than 800 ducks and measured a series of traits, including crest cushion, bill, plumage and shank colors, and body weight. In all families, we found that the crest trait followed the recessive inheritance of Mendel's law of separation after recoding. Furthermore, samples of blood from all F_0_, F_1_ and F_2_ ducks were obtained for DNA extraction and biochemical examination. Tissues were sampled for RNA or protein extraction and used in the transcriptome analyses. High-quality genomic DNA (gDNA) was then extracted using a traditional phenol-chloroform protocol. The purity and concentration of the gDNA samples were measured using a NanoDrop ND-2000 spectrophotometer (Thermo Scientific, MA, USA) and agarose gel electrophoresis. The final concentration was adjusted to 50 ng/μL, and gDNA samples with an A260/280 ratio of 1.8-2.0 were finally submitted for sequencing.

**5. Whole-genome sequencing**

***5.1 Sample collection and nucleic acid preparation***

For the genome assembly and annotation, the gDNA of a 28-week-old female CC duck, which came from our experimental base in Zhenjiang Tiancheng Agricultural Science and Technology Co., Ltd. (Zhenjiang, Jiangsu, China), was extracted from whole blood. gDNA was isolated using a traditional phenol-chloroform protocol.

***5.2 Library construction and sequencing***

For the preparation of the single-molecule real-time (SMAT) DNA template, the high-molecular-weight (HMW) genomic DNA was divided into large fragments (50 kbp average) by ultrasonication and then end-repaired according to the manufacturer’s instructions (Pacific Biosciences). The blunt hairpins and sequencing adaptor were ligated to the DNA fragments; DNA sequencing polymerases were bound to the SMRTbell templates. The library was quantified using a Qubit 4 Fluorometer (Invitrogen, USA). Then, PacBio SEQUEL platform was used for sequencing. Finally, we obtained 85.09 Gb (over 75 coverage) of long subreads data in total and used them for the sub-step genome assembly.

10X Genomics library construction: An automated micro-fluidic system allows the combination of the functionalized gel beads and high molecular weight DNA (HMW gDNA) together with oil to form a ‘gel bead in emulsion (GEM)’. Each GEM contains ~10 molecules of HMW gDNA and primers with unique barcodes and P5 sequencing adapters. After PCR amplification, P7 sequencing adapters were added for Illumina sequencing. Data processing: Firstly, 16 bp barcode sequences and 7 bp random sequences were trimmed on the reads 1. Then the unqualified paired reads were removed as well. Finally, we obtained 113.19 Gb raw bases with the 10X Genomics sequencing approach.

Two paired-end sequencing libraries with 350 bp of insert sizes were constructed according to Illumina’s protocol (Illumina, San Diego, CA, USA). Genomic DNA molecules were fragmented, end-paired and ligated to the adaptor. The ligated fragments were fractionated on agarose gels and purified by PCR amplification to produce sequencing libraries. A total of 89.16 Gb sequencing data were generated from Illumina’s paired-end sequencing and raw sequence data generated by Illumina platform were filtered according to the following criteria: (a) filtered reads with adapters; (b) filtered reads with N bases more than 10%; (c) filtered reads with low-quality bases (≤5) more than 50% of total length. If any member of the paired reads was classified as low quality, both pairs were discarded. After filtering, 88.65 Gb clean bases were obtained for *de novo* genome assembly.

Additionally, eight tissues, namely, the crested tissue, spleen, ovary, liver, duodenum, skin, pectoral, and blood, were sampled for genome annotation. Subsequently, all samples were subjected to RNA extraction using an RNAiso Pure RNA Isolation Kit (TaKaRa, Japan), which was followed by DNaseI treatment. A NanoVue Plus spectrophotometer (GE Healthcare, NJ, USA) was used to assess the concentration and quality of the extracted RNAs. All RNA samples were sequenced by Illumina HiSeq 4,000 to generate paired-end reads of 150 bp.

***5.3 Csp-b duck collection and sequencing***

We sequenced the genome of a 28-week-old female Csp-b duck using Illumina sequencing technology. Two short-insert (250 bp and 350 bp) and two long-insert (2 kb and 5 kb) DNA libraries were paired-end sequenced on the Illumina HiSeq 4000 platform (Table S26). Finally, 108.98 Gb raw data were generated, after removing the low-quality bases and paired reads with the Illumina adaptor sequence using SolexaQA++ 6 (version v.3.1.7.1). After filtering, 106.44 Gb clean bases were obtained for *de novo* genome assembly.

***5.4 Genome re-sequencing of all F_2_ generation samples***

The GWAS and genetic map analysis contained a total of 487 DNA samples, of which 308 were analyzed by GWAS and 179 (177 F_2_ individual and two parents) were used for genetic mapping. Two paired-end sequencing libraries with 350 bp of insert sizes were constructed according to Illumina’s protocol (Illumina, San Diego, CA, USA). All libraries were sequenced on the Illumina NovaSeq platform to an average clean read sequence coverage of ×11.60 and ×8.66 for the GWAS populations and the genetic map F_2_ individuals, respectively.

**6. *De novo* assembly and genetic map construction of the CC duck**

***6.1 De novo assembly of CC duck genome***

Three types of libraries of Illumina sequencing data, Pacbio sequencing data and 10X genomic sequencing data were used in different assembly stages separately. The Pacbio sequencing and 10X genomic sequencing were used for contig and primary scaffold assembly, and the genetic linkage map was used for chromosome-level scaffolding (Figure S5).

Before the CC duck genome assembly, we first surveyed the CC duck genome. In the genome survey stage, paired reads with “N” sites more than 10% of paired reads and low-quality (Q < 5) more than 20% of this paired reads were filtered out from the Illumina library. The pair reads containing the Illumina adaptor sequence were also filtered. Using Jellyfish[1], the frequency of 17-mers in the Illumina clean data was calculated with a 1 bp sliding window using the established method [2] and obeyed the theoretical Poisson distribution. Finally, the proportion of heterozygosity in the CC duck genome was evaluated as 0.55%, and the genome size was estimated as 1,257 Mb, with a repeat content of 40.61%.

The contig assembly of the duck genome was carried out using the FALCON (version 0.7) assembler [3], followed by one round of polishing with Quiver [4]. FALCON implements a hierarchical assembly process, which includes these steps: 1) subread error correction through aligning all reads to each other using daligner; the overlap data were then processed to generate error-corrected consensus reads; 2) second round of overlap detection using error-corrected reads; 3) construction of a directed string graph from overlap data; 4) resolving the contig path from the string graph. After FALCON assembly, the genome was polished by Quiver. Initial assembly of the PacBio data alone resulted in a total length of 1,115.45 Mb, with a contig N50 (the minimum length of contigs accounting for half of the haploid genome size) of 2.66 Mb. The fragScaff (version 140324.1) [5] software was mainly used for 10X Genomics Scaffolds Extending. The procedures are as follows: Linked-reads generated using the 10X Genomics library were aligned to the consensus sequence of PaBio assembly result to obtain the super-scaffold using BOWTIE v2.2 [6]. With the actual distance of consensus sequence increased, the number of linked-reads that support its connection will decrease. The consensus sequence without the linked-reads support will be filtered, and only the consensus sequence with the linked-reads support will be used for the subsequent assembly. FragScaff was used to generate 1,120.27 Mb scaffolds with a scaffold N50 length of 7.61 Mb. Finally, we used PBJelly [7] for gap filling and Illumina-derived short reads to correct any remaining errors by Pilon [8]. These processes yielded a final draft duck genome assembly (CC_duck _v1.0) which contained 2,027 contigs with a total length and contig N50 length of 1126.23 Mb and 3.24 Mb, respectively.

***6.2 Genetics linkage map construct***

***6.2.1 Variants calling and Genotyping***

BWA (Burrows-Wheeler Aligner) [9] was used to align the clean reads of each sample against the CC duck scaffold genome (settings: mem -t 4 -k 32 –M -R). Alignment files were converted to BAM files using SAMtools software [10] (settings: –bS –t). If multiple read pairs had identical external coordinates, only the pair with the highest mapping quality was retained. The sample alignment rate was between 97.27% and 98.03%. The average coverage depth for the reference genome (excluding the N region) was between 6.74X and 12.98X, and the 4X coverage (at least four) of the base was above 67.17%. The comparison results were normal and could be used for subsequent mutation detection and related analysis. The Genome Analysis Toolkit (GATK, version v3.7) [11] was used in variant calling for all samples with the UnifiedGenotyper method. SNP was filtered by the Perl script. A total of 7,301,165 parental polymorphism markers were obtained in this project, which were classified into eight segregation patterns (ab×cd, ef×eg, hk×hk, lm× ll, nn × np, aa × bb, ab × cc and cc × ab). For the F_2_ population, the "aa×bb" segregation patterns were choosing for genetic mapping, and the number of polymorphic markers was 1,231,700 (Table S27). Prior to the map construction, the markers with segregation distortion *(P* < 0.001), integrity (> 75%), or containing abnormal base were filtered. Finally, 226,870 markers were retained.

***6.2.2 Genetic linkage map construction***

The same marker was divided into bin markers by using Perl script. In the end, we obtained 23,735 single nucleotide polymorphism (SNP) markers. All markers were divided into 37 linkage groups using the independent LOD method in Joinmap (version 4.0) [12] software. The independent LOD starts with 20 and ends with 50, and the step is 5. Then, we used Joinmap software to sort the marks in every linkage group and calculate the genetic distance between markers. Finally, 5,795 bin markers were identified the chromosome. Genetic distance was calculated using R/qtl [13](est.map, error.prob = 0.005). The total genetic distance was 2,904.37 cM, with a mean marker density of 0.501 cM per marker (Table S28).

***6.2.3 Chromosome o******rientation***

Upstream and downstream of the marker position, a 100 bp marker sequence was extracted. The marker sequences were aligned to the reference genome using BWA. We constructed a genetic map with 37 autosomes of the CC duck genome and assembled a 1,126.23 Mb genome with a scaffold N50 length of 24.3 Mb using CHROMONMER (version 1.07) [14]. The Z chromosome was constructed by MUMmer (version 3.23) [15] based on sequence similarity using the published duck genome (CAU_duck1.0) as the reference genome. These processes yielded a final draft CC duck genome assembly (CC_duck_1.1) with a total length of 1,126.23 Mb, contig N50 of 3.24 Mb, and scaffold N50 of 73.74 Mb.

***6.2.4 CC duck genome assembly assessment***

We mapped the reads from short-insert length libraries to the CC duck genome with BWA and performed variant calling with SAMtools. With strict quality control and filtering, we obtained a total of 2,992,238 SNPs and noted that the homology rate (9.886×10^-4^) reflected a high single base accuracy. The single-copy orthologs were searched against the assembled genome of CC duck using the BUSCO tool (BUSCO, version 3.0.2) [16], which revealed that 97.7% complete and 1.6% partial of the 2,586 vertebrate BUSCOs are present in this assembly.

**7. CC duck genome annotation**

***7.1 Repeat sequence and annotation***

To identify the repeat sequences in CC duck genome, the homologous-based analysis of sequences and *ab initio* prediction approaches were used to identify repeats sequences in the CC duck genome. The commonly used homolog prediction RepBase database [17] employing RepeatMasker [18] (parameters: -a -nolow -no_is -norna -parallel 1) and its in-house scripts (RepeatProteinMask: parameters: -noLowSimple -pvalue 0.0001 -engine wublast) to extracted repeat regions. *Ab initio* prediction was used to build a *de novo* repetitive elements database by LTR_FINDER [19] (parameters: -C -w 2), PILER [20] (parameter ‘‘-trs’), RepeatScout [21] and RepeatModeler (parameters: -database genome -engine ncbi -pa 15) and then predict repeats by RepeatMasker. Tandem repeat was also extracted using TRF [22] (matching weight = 2, mismatching penalty = 7, INDEL penalty = 7, match probability = 80, INDEL probability = 10, minimum alignment score to report = 50, maximum period size to report = 2000, -d –h), by *ab initio* prediction. *De novo* and RepBase are the predicted TEs in the *de novo* database (predicted by RepeatModeler, RepeatScout, PILER and LTR_FINDER), and RepBase was applied to integrate the information using Uclust software according to 80-80-80 principles, with annotation performed by RepeatMasker. TE proteins are the predicted TEs based on RepBase database identified by RepeatProteinMask. Combined TEs are the combined results after eliminating redundant information. Others represent the repeats can annotation by RepeatMasker, but not including the species above. Unknown represent the repeats no annotated by RepeatMasker. Finally, after removing redundancies, we obtained 133,806,684 bp of repeats, accounting for 11.88% of the genome (Table S29).

***7.2 Structure annotation of protein coding genes***

For gene structure prediction, we employed homology-based, *de novo* based and RNA-seq-base data based to predict genes in the CC duck genome. For homologous comparison, the reference protein sequences of eleven species *Anser cygnoides, Aptenodytes forsteri, Anas platyrhynchos domestica, Coturnix japonica, Columba livia, Egretta garzetta, Gallus gallus, Homo sapiens, Nestor notabilis, Struthio camelus and Taeniopygia guttat*a from the Ensembl database (release 92) and NCBI database were aligned against the duck genome using BLAT search with parameters of an e-value ≤ 1e-5 in the “-F F” option. After filtering low-quality records, all blast hits were concatenated. The sequence of each candidate gene was further extended upstream and downstream by 1,000 bp to represent the whole region of this gene, within which the gene structure was predicted using the GeneWise tool [23]. Homology predictions were denoted as “Homology-set”. In the de novo approach, the packages and software of Augustus [24] (version 2.5.5), GlimmerHMM [25] (version 3.0.1), SNAP [26] (version 1.0), Geneid [27] (version 1.4.4) and GenScan [28] (version 1.0) also was used to predict the gene structure. In addition, we used RNA-seq data of eight tissues (Heart, liver, spleen, lung, kidney, skin, crested and breast muscles) which obtained approximately 47.66 Gb clean data, to predict the structure of transcribed genes using TopHat [29] (version 1.2) and Cufflinks [30] (version 2.2.1). Then, EvidenceModeler [31] (version 1.1.0) was used to combine the set of predicted genes generated from the three approaches into a non-redundant gene set, and PASA [32] (version 2.0.2) was used to annotate the gene structures. Weights for each type of evidence were set as follows: PASA-T-set > Homology-set > Cufflinks set > Augustus > GeneID = SNAP = GlimmerHMM = GeneScan. To obtain the UTRs and alternative splicing variation information, we used PASA to update the gene models. Finally, we successfully generated reference gene structures within the CC duck genome, which is composed of 17,425 protein-coding genes with a mean of 9.58 exons per gene (Table S30). The lengths of genes, coding sequence (CDS), introns, and exons in CC duck were comparable to those of closely-related genomes (Figure S17).

***7.3 Annotation of rRNAs, tRNAs and other non-coding RNAs***

We also predicted gene structures of tRNAs, rRNAs and other non-coding RNAs. A total of 402 tRNAs were predicted using the t-RNAscan-SE tool [33] (Table S31). Because rRNA genes are highly evolutionarily conserved, we chose the human rRNA sequence as references and then predicted 158 rRNA genes using the Blast tool [34] with an e-value of 1e-10. Small nuclear and nucleolar RNAs were annotated by the infernal tool [35] using the Rfam database [36].

***7.4 Functional annotations***

We functionally annotated the predicted proteins within the CC duck genome according to homologous searches against the databases of SwissProt [37] (Bairoch and Apweiler 2000), InterPro [38], NR database (from NCBI) [39] and Kyoto Encyclopedia of Genes and Genomes (KEGG) [40] with E values < 1e-5. The InterproScan tool in coordination with the InterPro database was applied to predict protein function based on the conserved protein domains and functional sites. KEGG pathway and SwissProt database were mainly mapped by the constructed gene set to identify the best match for each gene. In total, we annotated 16,577 (95.1%) genes function (Table S32).

**8. Csp-b duck genome assembly and annotation**

***8.1 Csp-b duck genome assembly***

The Csp-b duck genome was assembled using SOAPdenovo [41], a *de novo* genome assembler based on a *de Bruijn* graph algorithm. A *de Bruijn* graph was built by splitting the reads into *K*-mers, from the short insert size libraries (<1 kb), without making use of pairing information. After a series of graph simplifications, the reads were assembled into contigs. All available paired-end reads were realigned onto the contig sequences to construct the linkage between contigs. The linkage was removed if it was supported by an unreliable weight of paired-end relationships. We used the strategy of subgraph linearization to simplify the contig linkage graph, by extracting unambiguously linear paths. The GapCloser process was iterated in the order of estimated insert size step by step. Finally, for filling the intra-scaffold gaps, a local assembly was performed in order to locate the reads in the gap region, with the other end uniquely mapped to the contig. After filtering the scaffold less than 2,000 bp, we obtain a final Csp-b duck genome assembly with a total length of 1,102.39 Mb, contig N50 of 8.67 kb, scaffold N50 of 675.96 kb (Table S7).

***8.2 Csp-b duck genome annotation and assessment***

We used homologous comparison approaches to predict the protein-coding genes within the Csp-b duck genome. The reference protein sequences were from the Ensembl database (release 92) and NCBI database for the five species and CC duck. After redundancies were removed, the predicted genome of the Csp-b duck contained 15,278 genes, of which approximately 100% had functional annotations (Table S33-S34). In additionally, we also annotated 345 miRNAs, 198 tRNAs, 112 rRNAs and 457 snRNA for Csp-b duck genome (Table S35). Meanwhile, we assessment the Csp-b duck by BUSCO, and the result shown that 91.8% of 2,586 conserved vertebrate genes were assembled to complete (Table S36).

**9. Comparative genomics**

A total of 14 species were used in the gene family analysis, namely, *Anser cygnoides domesticus* (Acyg), *Aptenodytes forsteri* (Afor), *Balearica regulorum* (Breg), *Coturnix japonica* (Cjap), *Columba livia* (Cliv), CC duck, *Egretta garzetta* (Egar), *Gallus gallus* (Ggal), *Gavia stellata* (Gste), *Nestor notabilis* (Nnot), *Opisthocomus hoazin* (Ohoa), *Podiceps cristatus* (Pcri), *Struthio camelus* (Scam) and *Taeniopygia guttata* (Tgut). We filtered the gene sets of each species as follows: (1) when a gene had multiple alternative splicing transcripts, only the longest transcript in the coding region was reserved for further analysis, and (2) genes encoding proteins less than 30 amino acids long were excluded. Gene families were constructed through a hierarchical clustering algorithm and ‘all-against-all’ BLAST [32] (v2.2.26, -p blastp -e 1e-7 -F F). The alignments with high-scoring segment pairs (HSPs) were conjoined for each gene pair by Solar [33] (version 0.9.6). To identify homologous gene pairs, more than 40% coverage of the aligned regions in both homologous genes was required. Finally, homologous genes were clustered into gene families using hcluster_sg [34].

For the phylogenetic analysis, the phylogenetic tree was reconstructed using shared single-copy genes. Protein sequences for these single-copy genes were aligned by MUSCLE [35], and then the protein sequence alignment was transformed back to CDS alignments. We concatenated the CDS alignments of single-copy genes to a ‘supermatrix’. Using this supermatrix, we constructed the phylogenetic tree using the maximum likelihood (ML) algorithm as implemented in RAxML software (version 8.0.19) [36]. The mcmctree function from the PAML package was used for divergence time estimation with 8 calibration points from the TimeTree website (http://www.timetree.org/) [37]. The calibration points list was as follows: *Struthio camelus* and other species (min = 105 Mya, max = 118 Mya), the divergence time of *Taeniopygia guttata* and *Anser cygnoides domesticus* (92-104 Mya); *Aptenodytes forsteri* and *Egretta garzetta* (73-84 Mya); *Nestor notabilis* and *Egretta garzetta* (71-91 Mya); *Nestor notabilis* and *Columba livia* (77-90 Mya); *Aptenodytes forsteri* and *Balearica regulorum* (71-91 Mya); *Coturnix japonica* and *Anser cygnoides domesticus* (74-86 Mya); *Coturnix japonica* and *Gallus gallus* (33-42 Mya).

**10. Experimental validation**

PCR amplification was performed using PCR Gene Amplifier (Bio-Rad, USA) in a total volume of 25 μL, which contained 12 μL of 2 × Taq Master Mix (Dye Plus) (Vazyme, China), 1 μL (10 pmol) of each primer (Table S24), 1 μL of gDNA (100 ng) and 10 μL of ddH2O. After an initial denaturation for 3 min at 95°C, there were 35 amplification cycles (95°C for 15 s, 60°C for 15 s, and 72°C for 60 s) and a final extension for 5 min. PCR products were detected by 1% agarose gel electrophoresis. The PCR products were sequenced by Sangon (Shanghai, China).

Primers for RT-qPCR (Table S24) were designed by Oligo 6. *TAS2R40* was used to measure the expression level. Complementary DNA synthesis from total RNA and one-step quantitative PCR were performed using the Applied Biosystems QuantStudio5 system. The relative expression levels were quantified by SYBR Green-based RT-qPCR using PowerUp™ SYBR™ Green Master Mix (Applied Biosystems) according to the manufacturer's instructions. Three CC ducks of 56 days of age were slaughtered by stunning and exsanguination. Tissues samples including cerebellum, thigh muscle, breast muscle, cerebrum, liver, jejunum, bursa of fabricius, spleen, scalp of crested cushion, rectum, heart, kidney, scalp next to the crested cushion, subcutaneous fat, crested cushion, and abdominal fat (50-100 mg) were rapidly collected and snap-froze in liquid nitrogen and storage at -80°C. The RNA of these samples was used to determine the tissue expression profile of *TAS2R40*. The housekeeping gene, glyceraldehyde 3-phosphate dehydrogenase (*GAPDH*), was used as an endogenous control. All samples were assayed in at least three technical replicates. The collected data were analyzed using the 2^-ΔΔCt^ method.

Fragments of the 5'UTR of *TAS2R40* were cloned and inserted between the NheⅠ and XhoI restriction sites of the pGL-Basic 3.0 vector. The primer sequence information is shown in Table S24. The candidate SNP was mutated from G to A using the Fast Site‐Directed Mutagenesis Kit (Tiangen). All plasmids in this study were verified by DNA sequencing (Tsingke Biotechnology, Nanjing, China). SH-SY5Y cells were cotransfected with the pGL-Basic 3.0 luciferase reporter plasmid, the pGL-Control, pGL-Basic, and pR-TK using Lipofectamine™ 2000 (Invitrogen). Luciferase activity was measured 36 h following transfection using the Dual‐luciferase Reporter System (Promega, Madison, WI). Firefly luciferase activity was normalized to the corresponding Renilla luciferase activity.

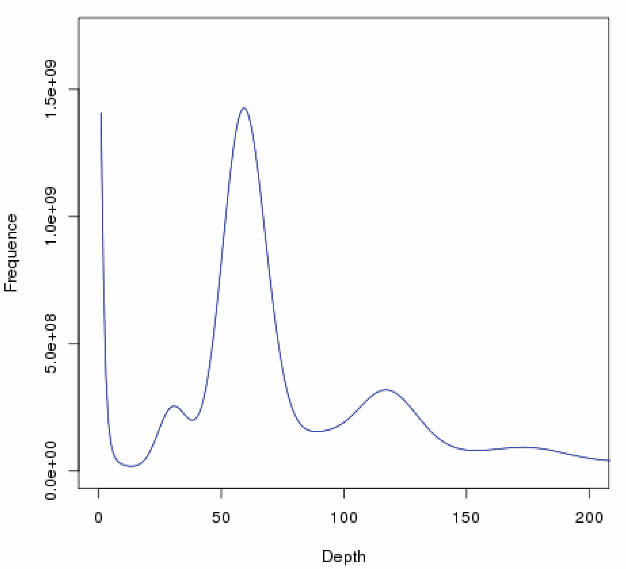
**Supplementary Figures**

**Figure S1.** Genome size estimation of the CC ducks using *K*-mer frequency distribution. The x-axis indicates the depth of each unique 17-mer in the CC duck genome, and the y-axis denotes the percentage of genome for occurrence of unique 17-mer within the sequence dataset. For this analysis, only paired-end reads were used after contamination filtering and base error correction.

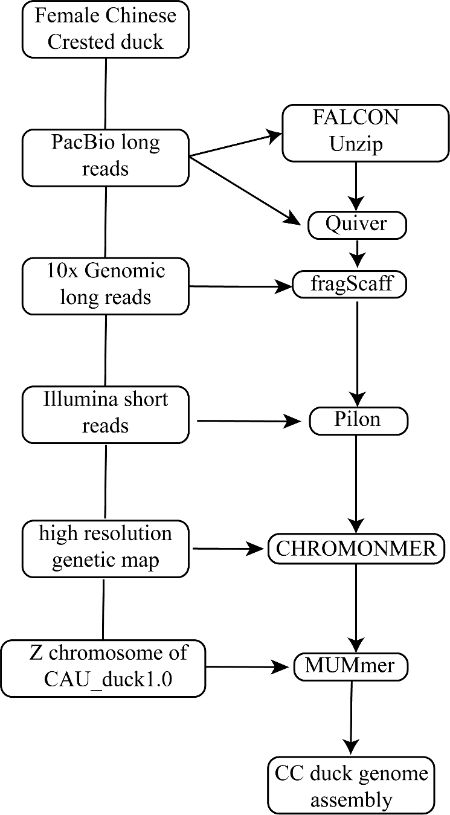

**Figure S2.** Genome assembly workflow.

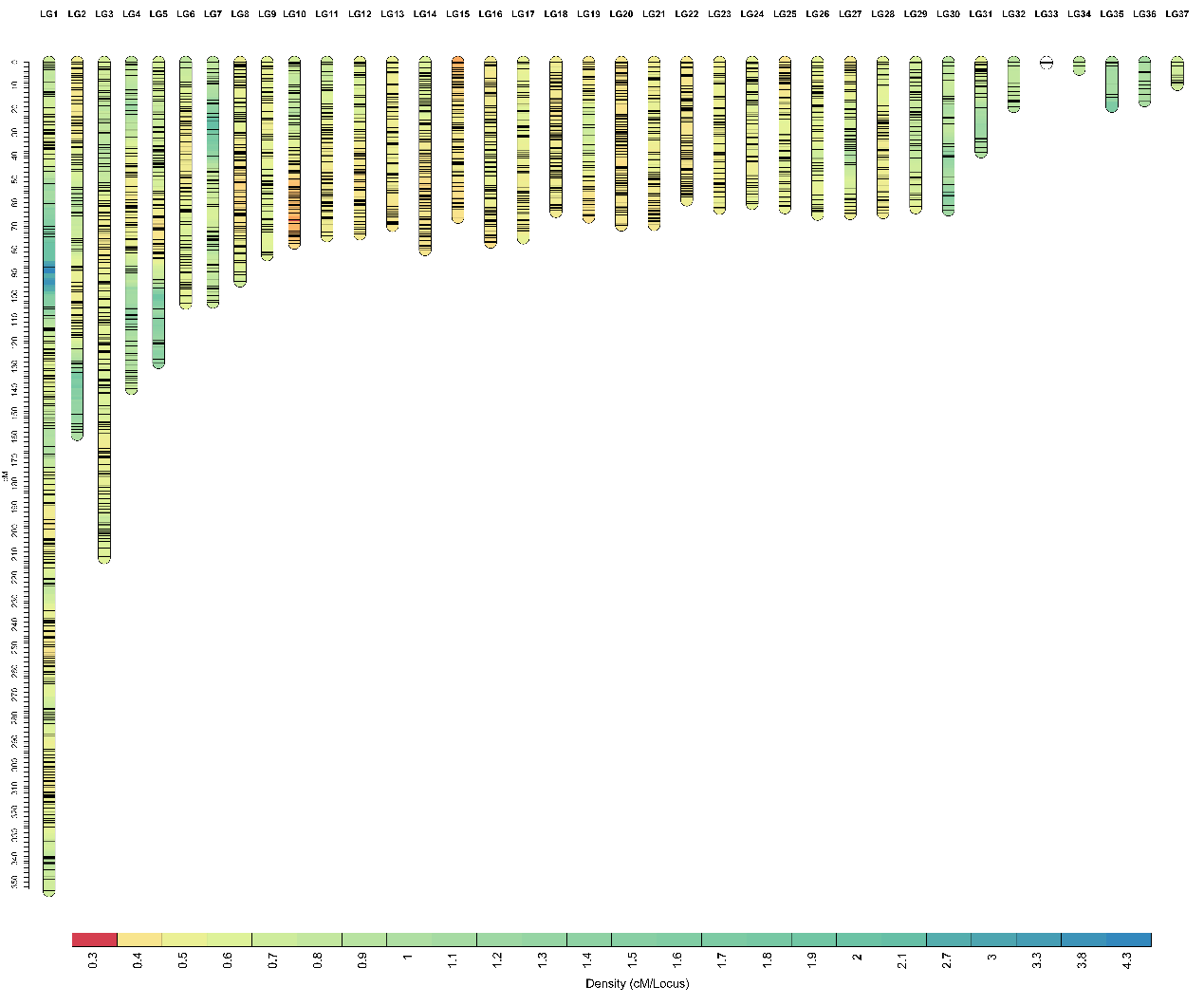

**Figure S3.** High-density linkage map of CC duck. The linkage maps were constructed with a total of 23,735 SNP markers. A total length of 2904.37 cM was mapped with 5,795 bin markers, and the average distance was 0.501 cM per maker.

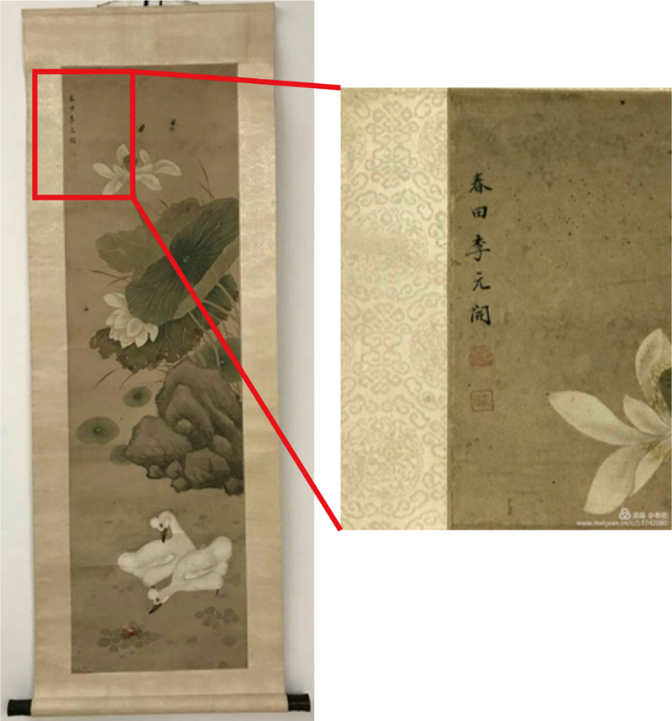

**Figure S4.** Li Yuankai's “Chuntian” ink Painting lotus and ducks.

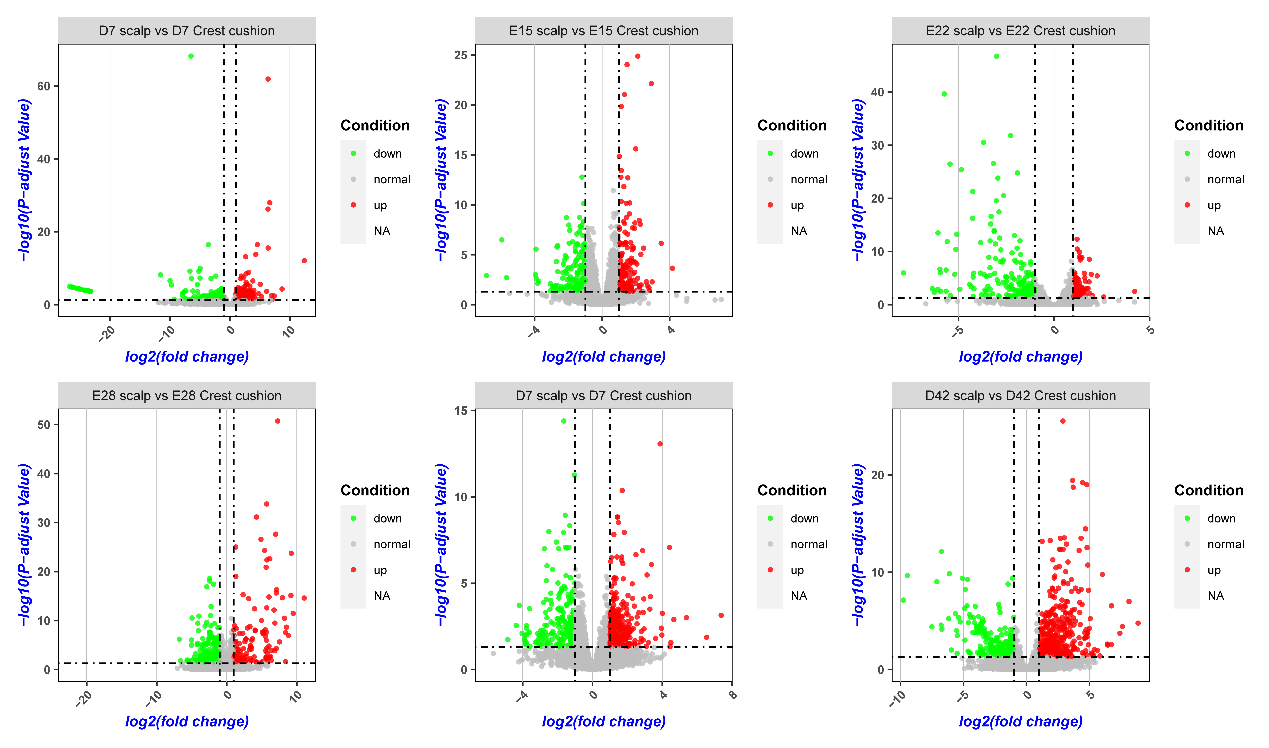

**Figure S5.** The volcano plot of DEGs of each CC duck development stage. Abundance of each gene was normalized as RPKM. DEGs are shown in red (up-regulated) and blue (down-regulated), while gray indicates genes that were not differentially expressed (no-DEGs). We used a false discovery rate ≤ 0.05 and the absolute value of log2Ratio ≥ 1 as the threshold to judge the significance of the differences in gene expression.

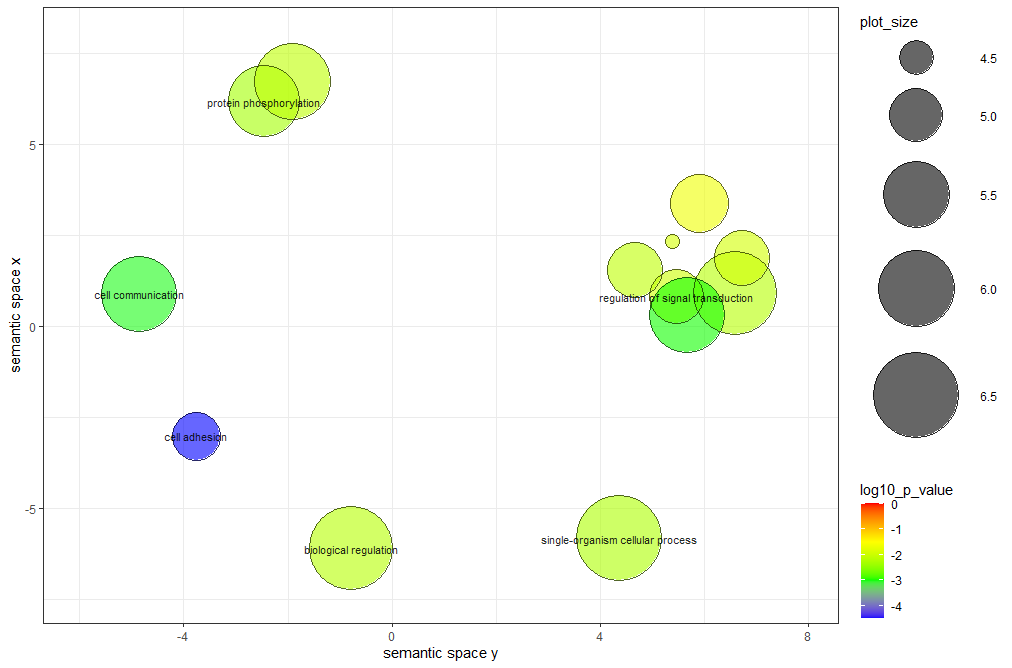

**Figure S6.** GO terms (biological process) of SV relate genes summarized and visualized as a REVIGO scatter plot.

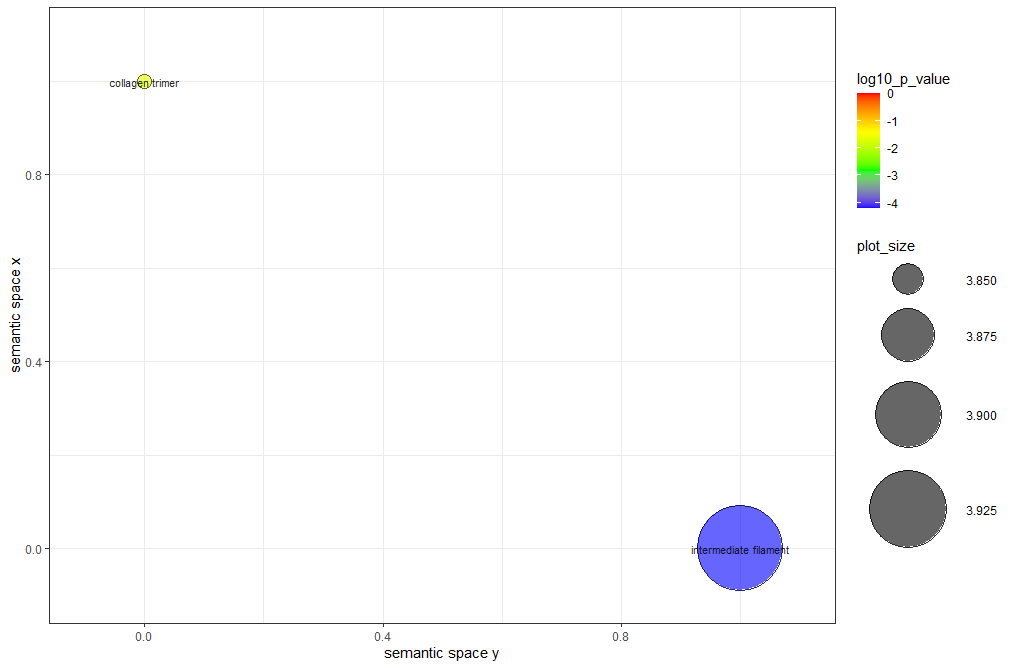

**Figure S7.** GO terms (cellular component) of SV relate genes summarized and visualized as a REVIGO scatter plot.

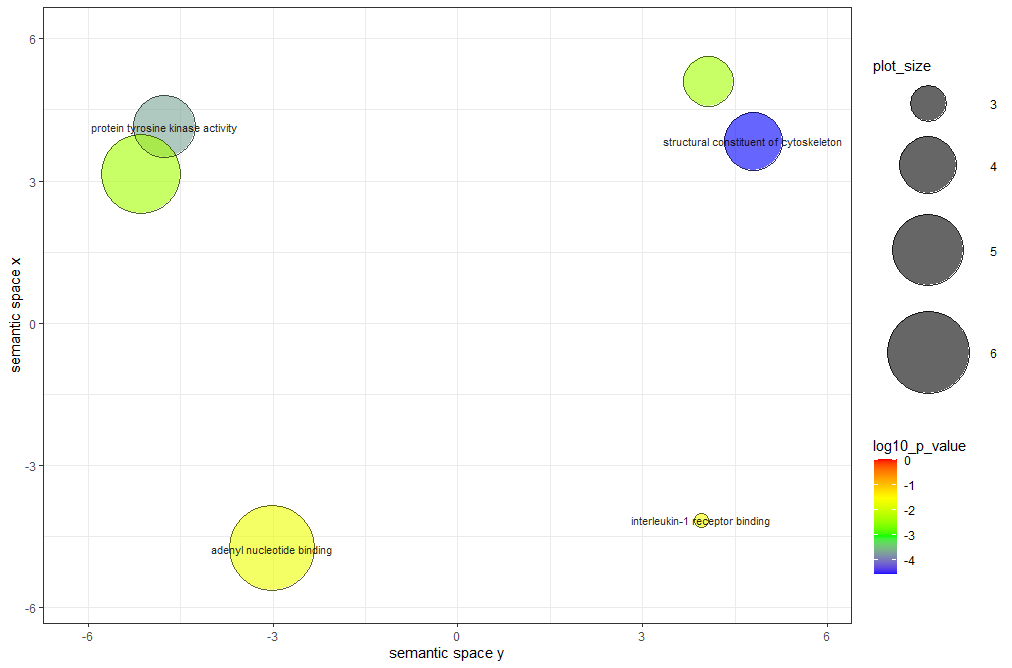

**Figure S8.** GO terms (molecular function) of SV relate genes summarized and visualized as a REVIGO scatter plot.

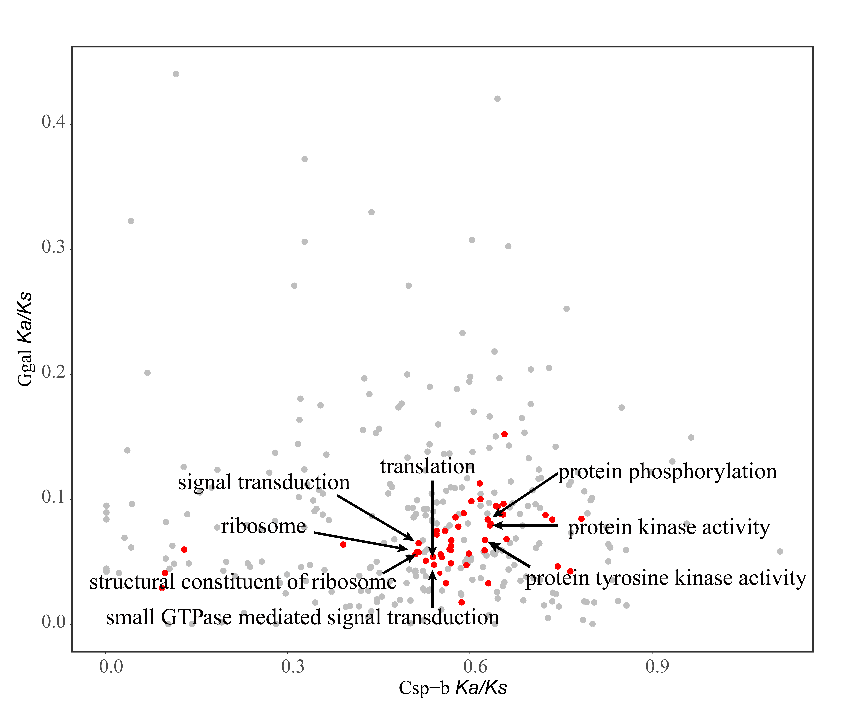

**Figure S9.** Data points represent pairs of Csp−b and *Gallus gallus* mean Ka/Ks ratios by GO category. GO categories with putatively accelerated (binomial test, FDR P value <0.05) nonsynonymous divergence in the Csp−b lineage (red) are highlighted.

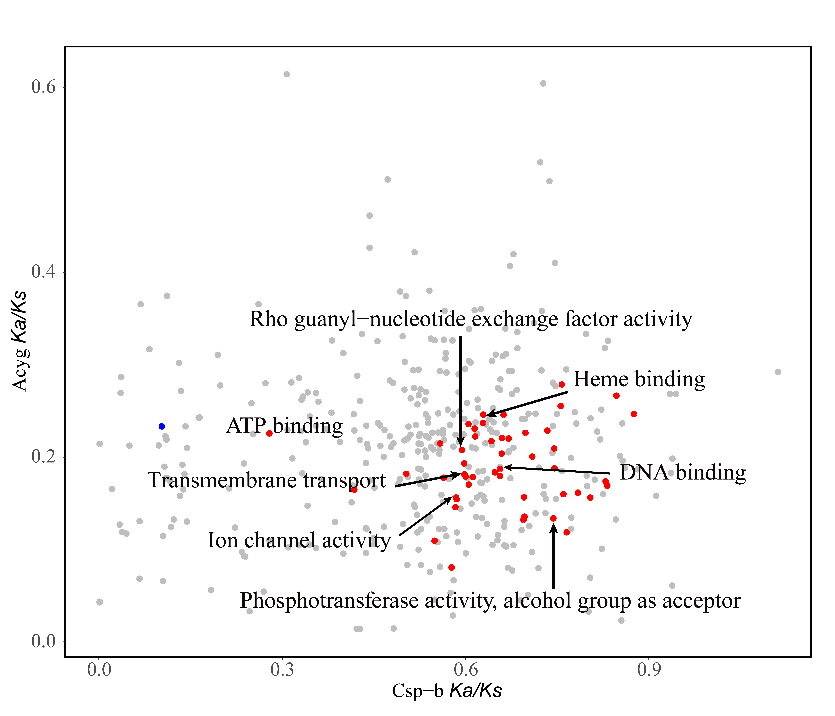

**Figure S10.** Data points represent pairs of Csp−b and *A. cygnoides* mean Ka/Ks ratios by GO category. GO categories with putatively accelerated (binomial test, FDR P value <0.05) nonsynonymous divergence in the Csp−b lineage (red) are highlighted.

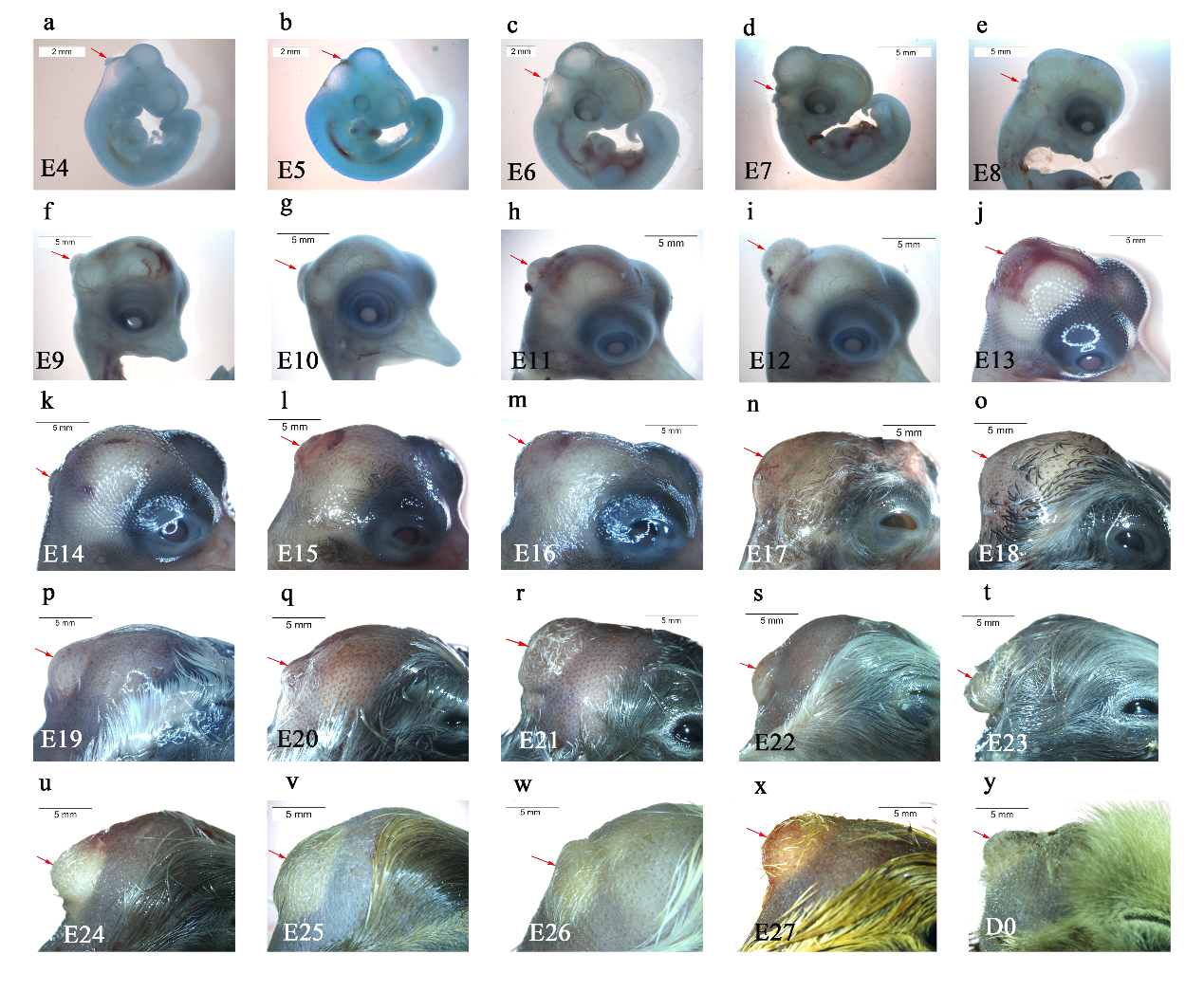

**Figure S11.** Morphology of the crest cushion at different stages of embryonic development. A protuberance of the head integument (arrows) (a: E4, b: E5, c: E6, d: E7, e: E8, f: E9, g: E10, h: E11, i: E12, j: E13, k: E14, l: E15, m: E16, n: E17, o: E18, p: E19, q: E20, r: E21, s: E22, t: E23, u: E24, v: E25, w: E26, x: E27, and y: D0). Scale bar = 2mm (a-c), scale bar = 5mm (d-y).

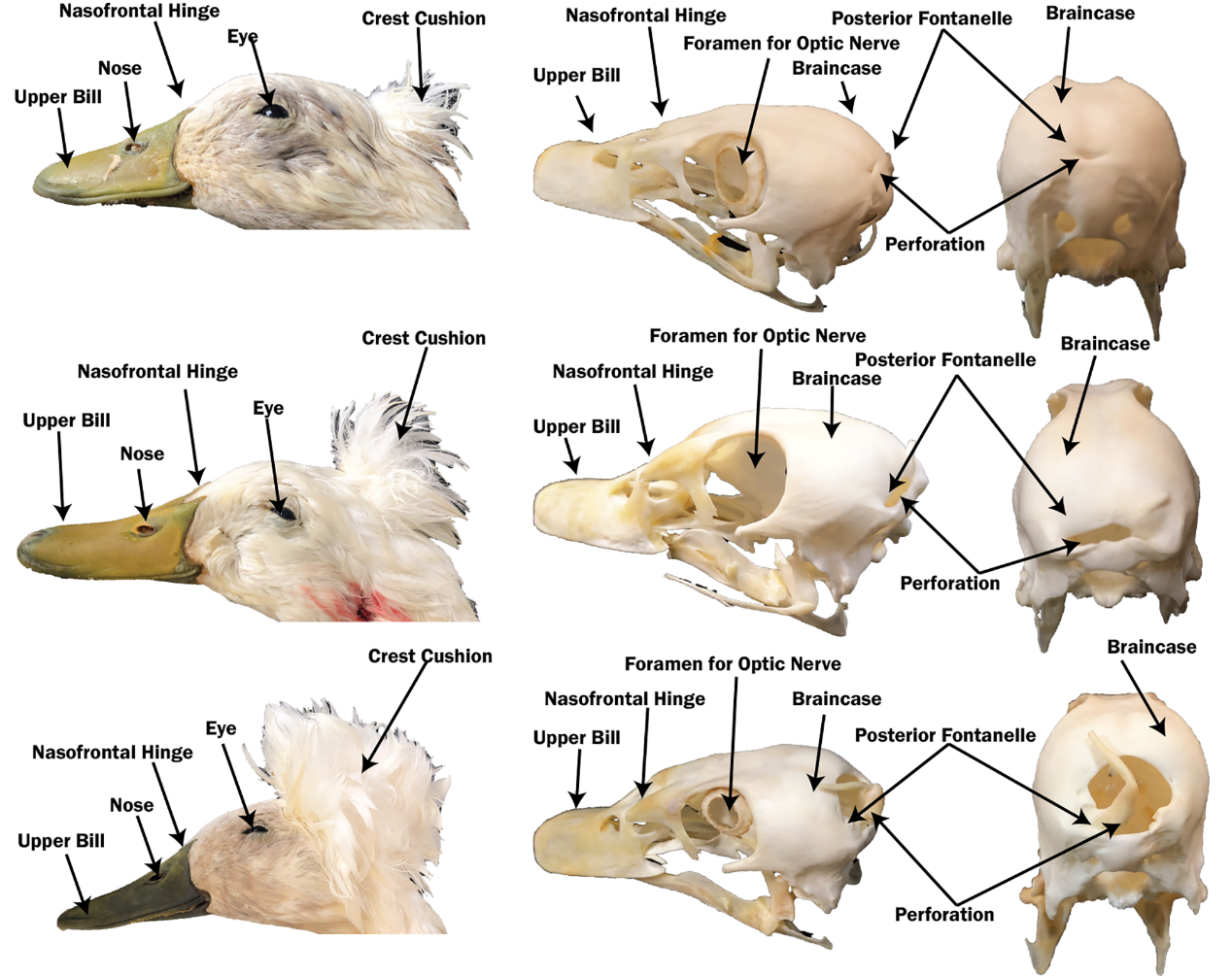

**Figure S12.** The head of CC duck with different size crested cushion. From top to bottom, there are small, medium and large crests.

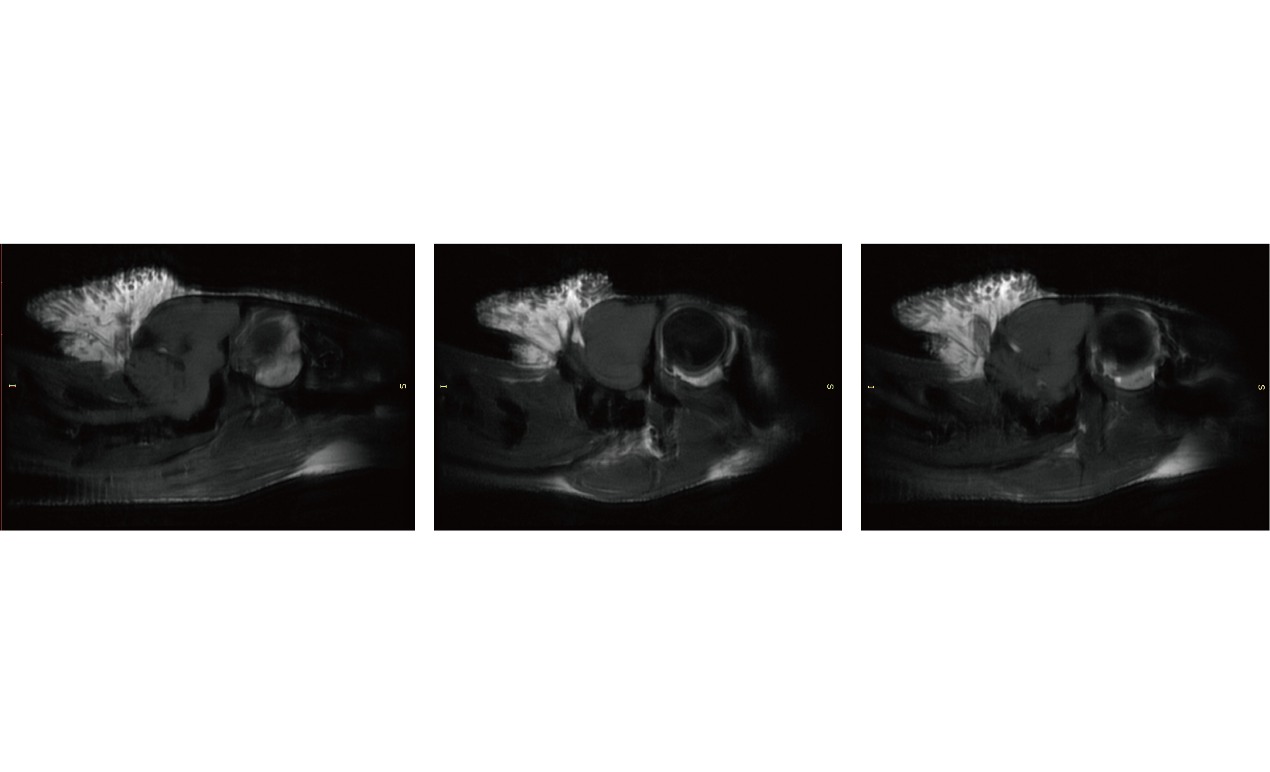

**Figure S13.** MR image of the head of a Crested duck with three sagittal adjacent slices.

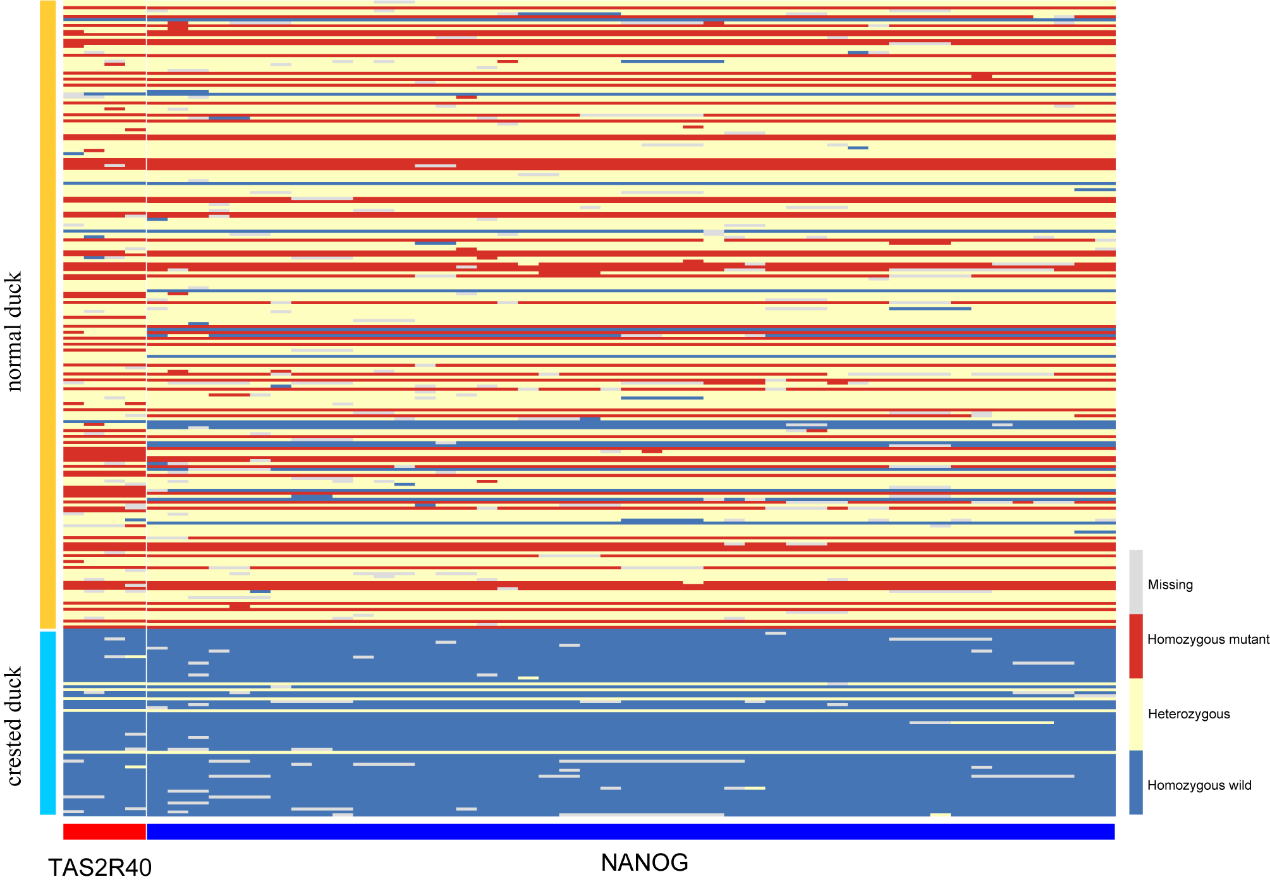

**Figure S14.** Heatmap the genotype of four SNPs in *TAS2R40* and *NANOG*. Blue means wild type; Yellow means heterozygous; Red means homozygous mutation; Gray means missing.

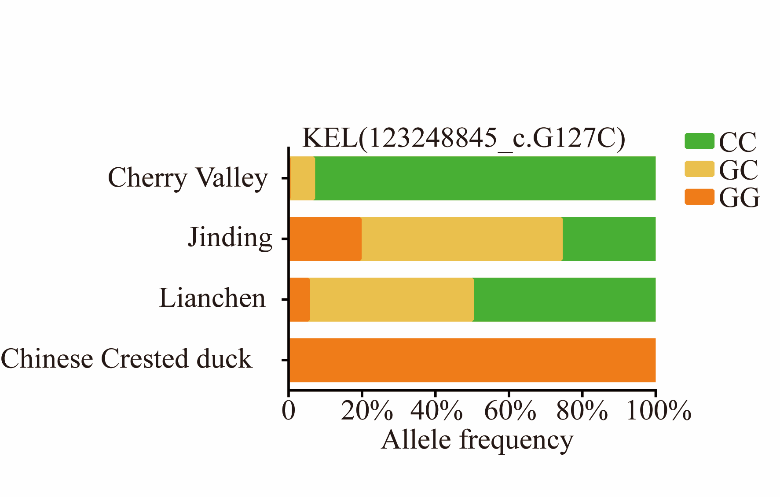

**Figure S15.** Genotype frequency of KEL (123248845_c. G127C).

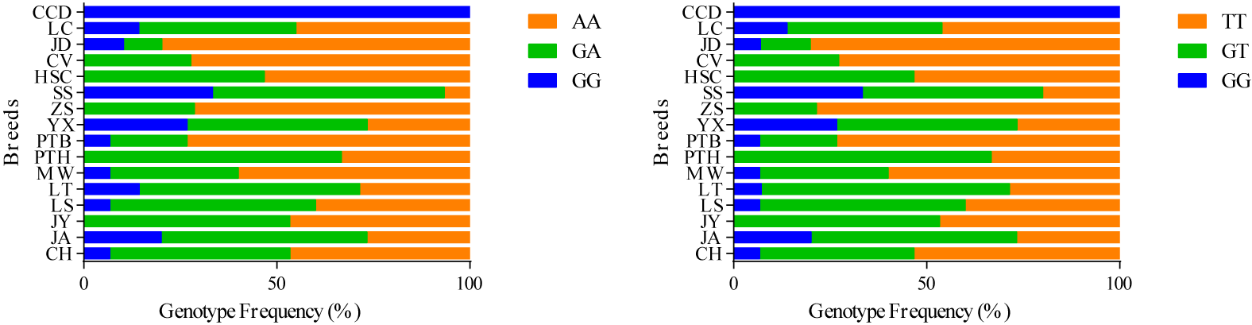

**Figure S16.** Genotype frequency of fourth exon in NANOG (120130992_c. G577A_p. V193M and 120131265_c. G850T: p. A284S). The CCD represent the Chinese creste duck; LC represent the Lianchen duck; JD represent the Jingding duck; HSC represent the Taiwanese Brown Vegetable Duck; SS represent the Sansui duck; ZS represent the Zhongshanma duck; YX represent the Youxianma duck; PTB represent the Putian white duck; PTH represent the Putian black duck; MW represent the Mawang duck; LT represent the mallard; LS represent the Longshencui duck; JY represent the Jingyunma duck; JA represent the Ji’an read feather duck and CH represent the Chaohu duck.

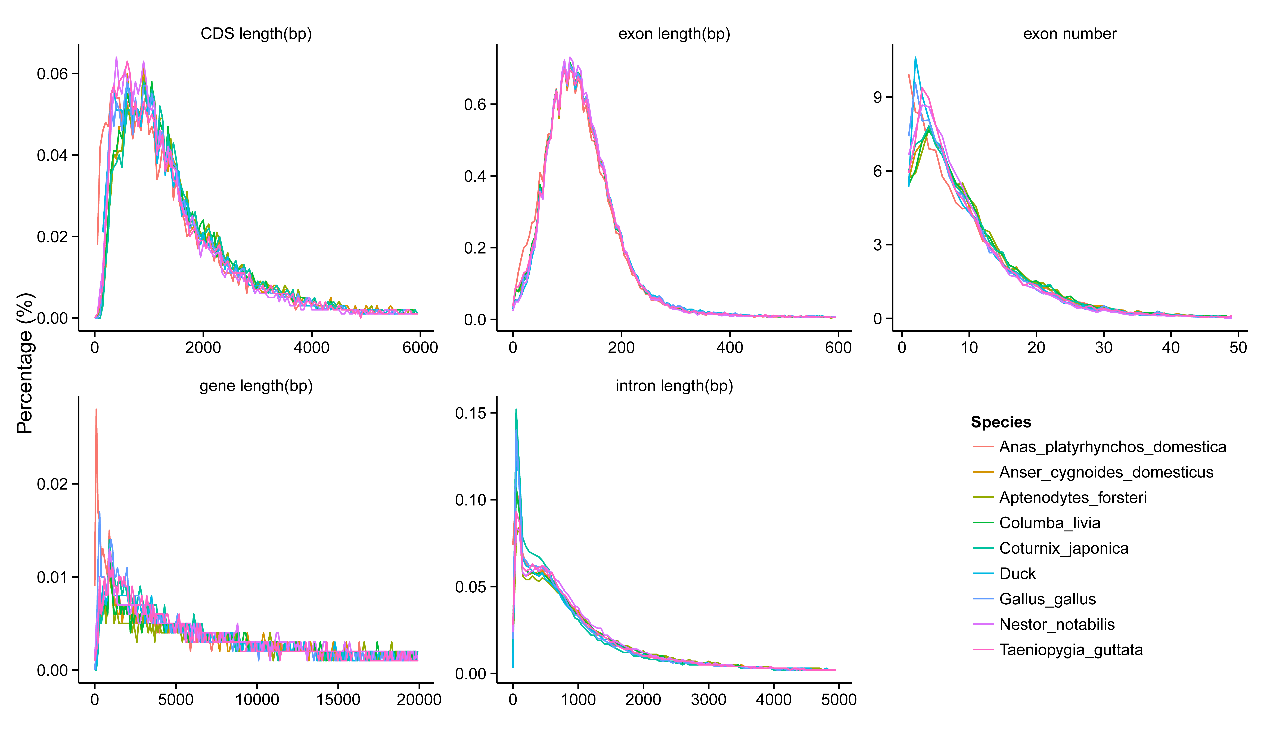

**Figure S17.** Distribution of gene elements in eight genetically related species. The x-axis indicates the length of gene, CDs, exon, intron and number of exon, and the y-axis represents gene number.

**Supplementary Tables**

**Table S1.** Estimation of Chinese crested (CC) duck genome size using *K*-mer analysis.

| K-mer | K-mer number | K-mer Depth | Genome Size (Mbp） | Revised Genome Size (Mbp） | Heterozygous Ratio (%) |
| --- | --- | --- | --- | --- | --- |
| 17 | 74,318,484,504 | 58 | 1,281.35 | 1,257.07 | 0.55 |

**Table S2.** Summary of Genome sequencing data for CC duck genome assembly.

| Pair-end libraries | Insert size(bp) | Total data (G) | Read length (bp) | Sequence coverage (X) |
| --- | --- | --- | --- | --- |
| Pacbio reads | 20K | 85.09 | - | 75.97 |
| 10X Genomics | 600 | 113.19 | 150.00 | 101.06 |
| Illumina reads | 350 | 88.65 | 150.00 | 79.15 |
| Total | - | 286.93 | - | 256.18 |

**Table S3.** Summary of the CC duck genome (CC_duck _v1.0) assembly.

| Sample ID | Length | | Number | |
| --- | --- | --- | --- | --- |
|  | Contig (bp) | Scaffold (bp) | Contig | Scaffold |
| Total | 1,124,559,267 | 1,126,228,291 | 2,027 | 1,216 |
| Max | 14,457,275 | 24,370,317 | - | - |
| ≥ 2 Kb | - | - | 1,984 | 1,173 |
| N50 | 3,244,619 | 7,607,602 | 90 | 46 |
| N60 | 2,453,473 | 6,216,808 | 129 | 62 |
| N70 | 1,709,527 | 4,381,292 | 183 | 84 |
| N80 | 1,070,837 | 2,663,331 | 267 | 117 |
| N90 | 450,014 | 1,155,826 | 423 | 180 |

**Table S4.** Summary of the CC duck genome (CC_duck_1.1) assembly.

| Sample ID | Length | | Number | | |
| --- | --- | --- | --- | --- | --- |
|  | Contig (bp) | Scaffold (bp) | Contig | | Scaffold |
| Total | 1,124,559,048 | 1,126,211,418 | | 2,025 | 863 |
| Max | 14,457,275 | 195,240,008 | - | | - |
| ≥ 2 Kb | - | - | 1,984 | | 821 |
| N50 | 3,244,619 | 73,735,783 | 90 | | 5 |
| N60 | 2,453,473 | 39,495,489 | 129 | | 7 |
| N70 | 1,709,527 | 27,727,581 | 183 | | 11 |
| N80 | 1,070,837 | 20,023,299 | 267 | | 16 |
| N90 | 450,014 | 7,382,291 | 423 | | 24 |

**Table S5** The genomic sequence of CC duck genome (CC_duck_1.1) integrity assessment.

|  |  | Percentage |
| --- | --- | --- |
| Reads | Mapping Rate (%) | 96.68 |
| Genome | Average sequencing Depth | 32.58 |
|  | Coverage (%) | 99.41 |
|  | Coverage at least 4X (%) | 99.17 |
|  | Coverage at least 10X (%) | 97.76 |
|  | Coverage at least 20X (%) | 87.03 |

**Table S6.** Statistics of BUSCO assessment of CC_duck_1.1 assembly.

| Species | Genome BUSCO evaluation results |
| --- | --- |
| CC duck | C:97.7%[S:96.6%,D:1.1%],F:1.6%,M:0.7%,n:2586 |
| Pekin duck | C:94.7%[S:93.7%,D:1.0%],F:1.9%,M:3.4%,n:2586 |

Note: C: Complete BUSCOs;

S: Complete and Single-Copy BUSCOs;

D: Complete and Duplicated BUSCOs;

F: Fragmented BUSCOs;

M: Missing BUSCOs;

n: Total BUSCO groups searched.

**Table S6.** Statistics of predicted protein-coding genes in the CC duck genome.

|  | Gene set | Gene number | Average transcript  length (bp) | Average CDS  length (bp) | Average exons  per gene | Average exon  length (bp) | Average intron  length (bp) |
| --- | --- | --- | --- | --- | --- | --- | --- |
| *De novo* | Augustus | 39,810 | 9,084.77 | 1,139.03 | 4.90 | 232.57 | 2,038.62 |
|  | GlimmerHMM | 206,079 | 4,684.80 | 513.42 | 2.69 | 191.07 | 2,472.49 |
|  | SNAP | 105,418 | 16,475.68 | 719.50 | 4.80 | 149.91 | 4,146.69 |
|  | Geneid | 46,497 | 17,297.08 | 1,227.98 | 5.28 | 232.43 | 3,751.64 |
|  | Genescan | 54,567 | 15,127.91 | 1,426.45 | 7.06 | 201.99 | 2,260.18 |
| Homolog | Acyg | 35,775 | 9,553.75 | 963.32 | 4.86 | 198.08 | 2,223.65 |
|  | Afor | 21,904 | 14,390.24 | 1,335.31 | 6.58 | 202.81 | 2,337.84 |
|  | Apla | 31,985 | 8,983.23 | 833.35 | 4.87 | 171.10 | 2,105.69 |
|  | Cjap | 19,981 | 17,085.92 | 1,432.51 | 7.66 | 186.90 | 2,348.80 |
|  | Cliv | 22,811 | 14,260.60 | 1,159.12 | 6.46 | 179.55 | 2,401.35 |
|  | Egar | 20,075 | 15,761.78 | 1,385.84 | 7.06 | 196.28 | 2,372.10 |
|  | Ggal | 35,950 | 8,891.72 | 819.56 | 4.52 | 181.35 | 2,293.64 |
|  | Hsap | 14,572 | 20,511.97 | 1,573.35 | 8.75 | 179.71 | 2,442.14 |
|  | Nnot | 19,781 | 12,015.58 | 1,257.67 | 6.05 | 207.75 | 2,128.70 |
|  | Scam | 21,095 | 15,057.91 | 1,358.22 | 6.80 | 199.63 | 2,360.45 |
|  | Tgut | 25,334 | 14,204.50 | 1,061.95 | 5.72 | 185.59 | 2,783.22 |
| RNA-seq | PASA | 100,396 | 14,329.69 | 1,055.58 | 6.00 | 175.87 | 2,653.80 |
|  | Cufflinks | 87,156 | 20,594.98 | 3,406.92 | 6.86 | 496.33 | 2,930.99 |
| EVM | | 22,524 | 17,728.93 | 1,322.28 | 7.70 | 171.77 | 2,449.48 |
| PASA-update | | 22,360 | 19,350.63 | 1,338.06 | 7.74 | 172.88 | 2,672.60 |
| Final set | | 17,425 | 24,484.99 | 1,599.03 | 9.58 | 166.86 | 2,666.39 |

Note: Acyg represents for *Anser cygnoides* domesticus; Afor represents for Aptenodytes forsteri; Apla represents for *Anas platyrhynchos* domestica; Cjap represents for *Coturnix japonica*; Cliv represents for *Columba livia*; Egar represents for *Egretta garzetta*; Ggal represents for *Gallus gallus*; Hsap represents for *Homo sapiens*; Nnot represents for *Nestor notabilis*; Scam represents for *Struthio camelus*; Tgut represents for *Taeniopygia guttata*.

**Table S7.** Summary of the Csp-b duck genome assembly.

| Sample ID | length | | number | |
| --- | --- | --- | --- | --- |
|  | Contig | Scaffold | Contig | Scaffold |
| Total | 1,029,904,886 | 1,102,391,966 | 231,908 | 7,534 |
| Max | 107,550 | 4,406,617 | - | - |
| Number>=2000 | - | - | 133,092 | 7,534 |
| N50 | 8,668 | 675,958 | 34,711 | 454 |
| N60 | 6,922 | 530,777 | 48,019 | 637 |
| N70 | 5,381 | 391,923 | 64,881 | 877 |
| N80 | 3,923 | 277,153 | 87,209 | 1,208 |
| N90 | 2,441 | 151,176 | 120,101 | 1,735 |

**Table S8.** The statistic results of gene organization in relative species.

| #Gene set | Number of genes | CDS+ intron length (average) | CDS length (average) | exon length (average) | intron length (average) | Exons per gene (average) |
| --- | --- | --- | --- | --- | --- | --- |
| Acyg | 14,696 | 30,176.71 | 1,763.51 | 162.58 | 2,885.50 | 10.85 |
| Apla | 15,634 | 21,091.75 | 1,507.51 | 148.94 | 2,147.05 | 10.12 |
| Csp-b duck | 15,278 | 22,327.78 | 1,638.79 | 176.41 | 2,495.83 | 9.29 |
| CC duck | 17,425 | 24,484.99 | 1,599.03 | 166.86 | 2,666.39 | 9.58 |
| Ggal | 16,362 | 23,425.95 | 1,603.16 | 165.66 | 2,514.86 | 9.68 |
| Tgut | 16,348 | 25,502.91 | 1,620.25 | 161.88 | 2,650.91 | 10.01 |

Note: Acyg represents for *Anser cygnoides* domesticus; Apla represents for *Anas platyrhynchos* domestica; Ggal represents for *Gallus gallus*; Tgut represents for *Taeniopygia guttata*.

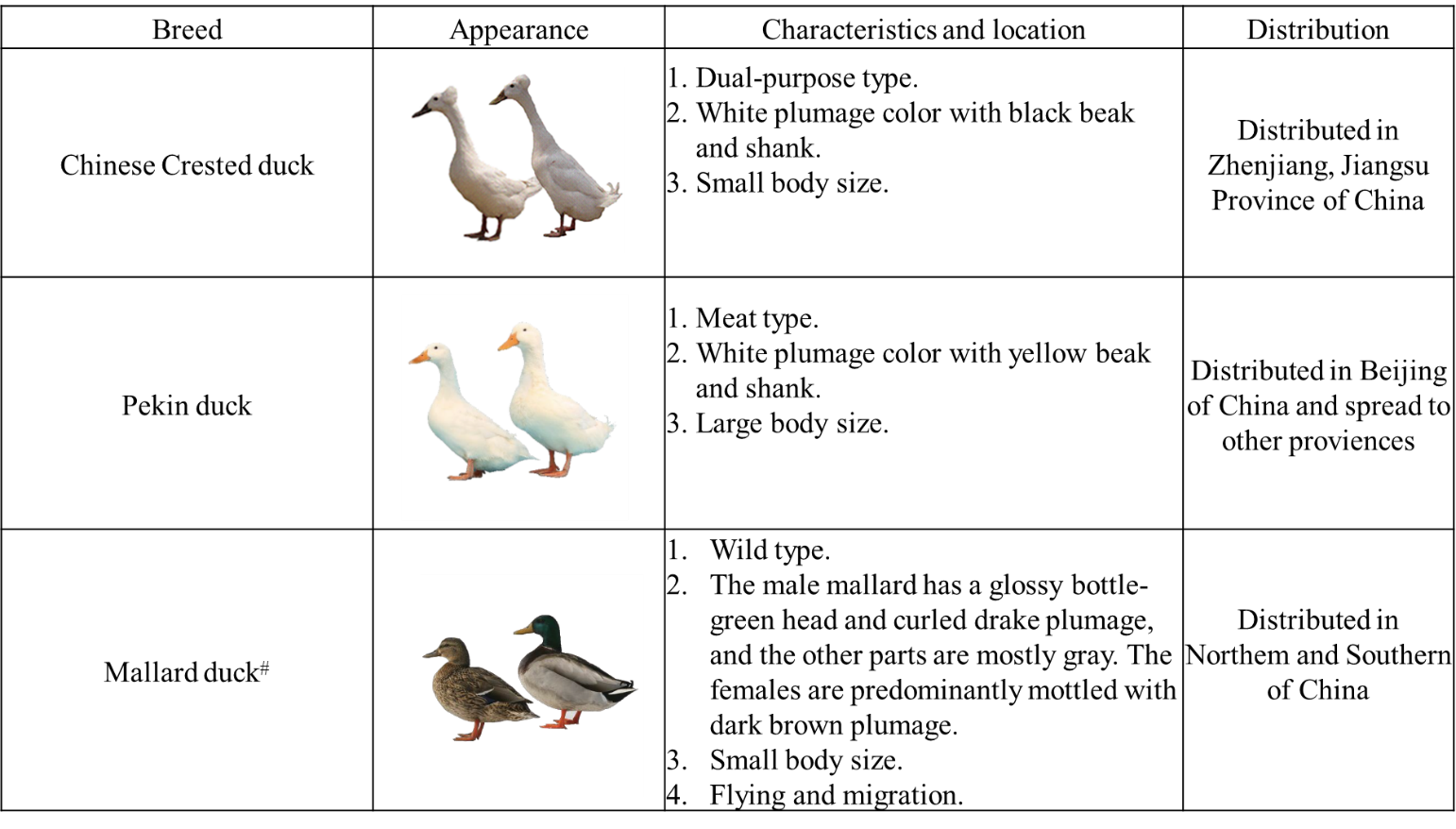
**Table S9.** The information of duck breeds in the present study.

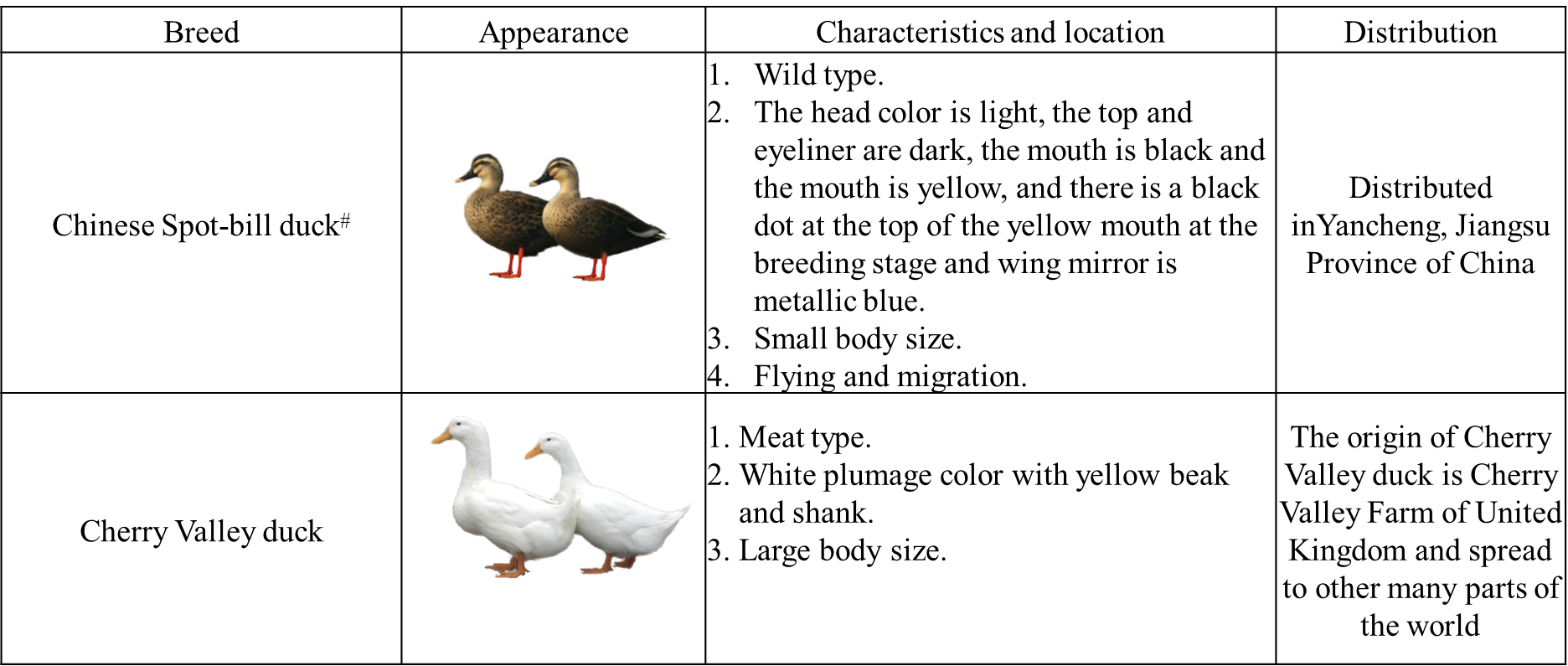

The characteristics, including appearance, location and economic type are list follows. The ^#^ indicates that the male ducks have a glossy green head.

**Table S10.** The information of genome which used to compare genome analysis.

| Species | ID | #gene |
| --- | --- | --- |
| *Anser cygnoides* | Acyg | 14,696 |
| *Aptenodytes forsteri* | Afor | 14,550 |
| *Balearica regulorum* | Breg | 15,000 |
| *Coturnix japonica* | Cjap | 16,024 |
| *Columba livia* | Cliv | 15,562 |
| *Anas platyrhynchos domestica* | Duck | 17,425 |
| *Egretta garzetta* | Egar | 14,123 |
| *Gallus gallus* | Ggal | 18,342 |
| *Gavia stellata* | Gste | 14,661 |
| Nestor notabilis | Nnot | 14,736 |
| *Opisthocomus hoazin* | Ohoa | 13,333 |
| *Podiceps cristatus* | Pcri | 11,707 |
| *Struthio camelus* | Scam | 14,577 |
| *Taeniopygia guttata* | Tgut | 17,472 |
| Total |  | 212,208 |

**Table S11.** The detailed lists of GO terms of expanded gene families.

| GO ID | GO Term | GO Class | Adjusted *P*-value |
| --- | --- | --- | --- |
| GO:0043232 | intracellular non-membrane-bounded organelle | CC | 1.24E-11 |
| GO:0007155 | cell adhesion | BP | 3.03E-07 |
| GO:0006352 | DNA-templated transcription, initiation | BP | 3.28E-07 |
| GO:0044446 | intracellular organelle part | CC | 7.42E-07 |
| GO:0044430 | cytoskeletal part | CC | 1.02E-06 |
| GO:0005856 | cytoskeleton | CC | 1.06E-06 |
| GO:0015630 | microtubule cytoskeleton | CC | 1.61E-06 |
| GO:0007017 | microtubule-based process | BP | 2.06E-06 |
| GO:0042813 | Wnt-activated receptor activity | MF | 3.27E-06 |
| GO:0006351 | transcription, DNA-templated | BP | 5.95E-06 |
| GO:0000786 | nucleosome | CC | 6.35E-06 |
| GO:0034654 | nucleobase-containing compound biosynthetic process | BP | 7.86E-06 |
| GO:0043234 | protein complex | CC | 9.17E-06 |
| GO:0006334 | nucleosome assembly | BP | 1.11E-05 |
| GO:0005200 | structural constituent of cytoskeleton | MF | 2.71E-05 |
| GO:0005874 | microtubule | CC | 3.42E-05 |
| GO:0003677 | DNA binding | MF | 6.29E-05 |
| GO:0022607 | cellular component assembly | BP | 6.32E-05 |
| GO:0005149 | interleukin-1 receptor binding | MF | 9.87E-05 |
| GO:0005488 | binding | MF | 0.000142882 |
| GO:0005871 | kinesin complex | CC | 0.00020003 |
| GO:0006461 | protein complex assembly | BP | 0.000266802 |
| GO:0045296 | cadherin binding | MF | 0.000382788 |
| GO:0032403 | protein complex binding | MF | 0.000396222 |
| GO:0003774 | motor activity | MF | 0.000449488 |
| GO:0008092 | cytoskeletal protein binding | MF | 0.000499352 |
| GO:0008017 | microtubule binding | MF | 0.000543706 |
| GO:0003777 | microtubule motor activity | MF | 0.000758373 |
| GO:0007018 | microtubule-based movement | BP | 0.000921245 |
| GO:0007585 | respiratory gaseous exchange | BP | 0.001060707 |
| GO:0005152 | interleukin-1 receptor antagonist activity | MF | 0.001060707 |
| GO:0016055 | Wnt signaling pathway | BP | 0.001720108 |
| GO:0007156 | homophilic cell adhesion | BP | 0.002264547 |
| GO:0007399 | nervous system development | BP | 0.002363864 |
| GO:0007416 | synapse assembly | BP | 0.002855526 |
| GO:0048013 | ephrin receptor signaling pathway | BP | 0.003224831 |
| GO:0043565 | sequence-specific DNA binding | MF | 0.003708069 |
| GO:0003676 | nucleic acid binding | MF | 0.004379081 |
| GO:0005509 | calcium ion binding | MF | 0.006035406 |
| GO:0006259 | DNA metabolic process | BP | 0.006560727 |
| GO:0006355 | regulation of transcription, DNA-templated | BP | 0.00864148 |
| GO:0003700 | sequence-specific DNA binding transcription factor activity | MF | 0.00864148 |
| GO:0016337 | single organismal cell-cell adhesion | BP | 0.014716038 |
| GO:0016818 | hydrolase activity, acting on acid anhydrides, in phosphorus-containing anhydrides | MF | 0.014749275 |
| GO:0016043 | cellular component organization | BP | 0.015140515 |
| GO:0005515 | protein binding | MF | 0.015880919 |
| GO:0043229 | intracellular organelle | CC | 0.017086721 |
| GO:0019789 | SUMO ligase activity | MF | 0.017307003 |
| GO:0015629 | actin cytoskeleton | CC | 0.017737564 |
| GO:0008146 | sulfotransferase activity | MF | 0.018312533 |
| GO:0005198 | structural molecule activity | MF | 0.036856535 |
| GO:1901363 | heterocyclic compound binding | MF | 0.037064387 |
| GO:0048666 | neuron development | BP | 0.038675779 |
| GO:0097159 | organic cyclic compound binding | MF | 0.03870781 |
| GO:0090129 | positive regulation of synapse maturation | BP | 0.039271302 |
| GO:0005540 | hyaluronic acid binding | MF | 0.042752582 |
| GO:0005615 | extracellular space | CC | 0.045939448 |
| GO:0005044 | scavenger receptor activity | MF | 0.04817562 |

Note: BP for Biological Process, MF for Molecular Function, and CC for Cellular Component.

**Table S12.** The detailed lists of KEGG pathways of expanded gene families.

| Map ID | Map Title | Adjusted *P*-value |
| --- | --- | --- |
| map05130 | Pathogenic Escherichia coli infection | 4.22E-06 |
| map04390 | Hippo signaling pathway | 1.08E-05 |
| map05322 | Systemic lupus erythematosus | 0.000398036 |
| map05034 | Alcoholism | 0.000647261 |
| map04514 | Cell adhesion molecules (CAMs) | 0.002271166 |
| map05200 | Pathways in cancer | 0.003106249 |
| map04540 | Gap junction | 0.003348338 |
| map04916 | Melanogenesis | 0.007751174 |
| map04550 | Signaling pathways regulating pluripotency of stem cells | 0.010514925 |
| map05217 | Basal cell carcinoma | 0.011855086 |
| map04360 | Axon guidance | 0.014518697 |
| map05213 | Endometrial cancer | 0.014518697 |
| map05100 | Bacterial invasion of epithelial cells | 0.020645047 |
| map02010 | ABC transporters | 0.020645047 |
| map04924 | Renin secretion | 0.022652707 |
| map05203 | Viral carcinogenesis | 0.02371033 |
| map00053 | Ascorbate and aldarate metabolism | 0.02928038 |
| map00982 | Drug metabolism - cytochrome P450 | 0.02928038 |
| map00040 | Pentose and glucuronate interconversions | 0.030446281 |
| map00860 | Porphyrin and chlorophyll metabolism | 0.031640755 |
| map00534 | Glycosaminoglycan biosynthesis - heparan sulfate / heparin | 0.03286164 |
| map04145 | Phagosome | 0.036075632 |
| map04930 | Type II diabetes mellitus | 0.039669026 |
| map04913 | Ovarian steroidogenesis | 0.04314923 |
| map04520 | Adherens junction | 0.04314923 |
| map00830 | Retinol metabolism | 0.045242635 |
| map00980 | Metabolism of xenobiotics by cytochrome P450 | 0.046602917 |

**Table S13.** The detailed lists of GO terms of contracted gene families.

| GO ID | GO Term | GO Class | Adjusted *P*-value |
| --- | --- | --- | --- |
| GO:0005200 | structural constituent of cytoskeleton | MF | 0.247227272 |
| GO:0051258 | protein polymerization | BP | 0.247227272 |
| GO:0005874 | microtubule | CC | 0.247227272 |
| GO:0004252 | serine-type endopeptidase activity | MF | 0.247227272 |

Note: BP for Biological Process, MF for Molecular Function, and CC for Cellular Component.

**Table S14.** The detailed lists of KEGG pathways of contracted gene families.

| Map ID | Map Title | Adjusted P-value |
| --- | --- | --- |
| map04670 | Leukocyte transendothelial migration | 0.14744998 |
| map04972 | Pancreatic secretion | 0.14744998 |
| map04924 | Renin secretion | 0.22269293 |
| map05416 | Viral myocarditis | 0.22269293 |
| map05032 | Morphine addiction | 0.24129941 |
| map04723 | Retrograde endocannabinoid signaling | 0.41375538 |
| map04510 | Focal adhesion | 0.44242035 |
| map05150 | Staphylococcus aureus infection | 0.44242035 |
| map05310 | Asthma | 0.44242035 |
| map00601 | Glycosphingolipid biosynthesis - lacto and neolacto series | 0.44242035 |
| map04727 | GABAergic synapse | 0.46275399 |
| map04611 | Platelet activation | 0.50774486 |
| map04514 | Cell adhesion molecules (CAMs) | 0.59788648 |
| map04930 | Type II diabetes mellitus | 0.59788648 |
| map04923 | Regulation of lipolysis in adipocytes | 0.59788648 |
| map05033 | Nicotine addiction | 0.59788648 |

**Table S15.** The detailed lists of GO terms of lineage-specific genes.

| GO ID | GO Term | GO Class | Adjusted P-value |
| --- | --- | --- | --- |
| GO:0005871 | kinesin complex | CC | 0.00791139 |
| GO:0008017 | microtubule binding | MF | 0.00791139 |
| GO:0003777 | microtubule motor activity | MF | 0.00964523 |
| GO:0003774 | motor activity | MF | 0.01022696 |
| GO:0007018 | microtubule-based movement | BP | 0.01022696 |
| GO:0006928 | cellular component movement | BP | 0.01847193 |
| GO:0005875 | microtubule associated complex | CC | 0.02833617 |
| GO:0005509 | calcium ion binding | MF | 0.0303842 |
| GO:0043395 | heparan sulfate proteoglycan binding | MF | 0.03343361 |
| GO:0032403 | protein complex binding | MF | 0.03764866 |
| GO:0007585 | respiratory gaseous exchange | BP | 0.03764866 |
| GO:0019310 | inositol catabolic process | BP | 0.03764866 |
| GO:0050113 | inositol oxygenase activity | MF | 0.03764866 |

Note: BP for Biological Process, MF for Molecular Function, and CC for Cellular Component.

**Table S16.** The detailed lists of KEGG pathways of lineage-specific genes.

| Map ID | Map Title | Adjusted *P*-value |
| --- | --- | --- |
| map05416 | Viral myocarditis | 0.419534933 |
| map05150 | Staphylococcus aureus infection | 0.419534933 |
| map00601 | Glycosphingolipid biosynthesis - lacto and neolacto series | 0.419534933 |
| map05310 | Asthma | 0.419534933 |
| map04514 | Cell adhesion molecules (CAMs) | 0.419534933 |
| map00562 | Inositol phosphate metabolism | 0.463173534 |
| map04810 | Regulation of actin cytoskeleton | 0.463173534 |
| map04670 | Leukocyte transendothelial migration | 0.463173534 |
| map04972 | Pancreatic secretion | 0.556599206 |
| map05032 | Morphine addiction | 0.559041091 |
| map04370 | VEGF signaling pathway | 0.559041091 |
| map04960 | Aldosterone-regulated sodium reabsorption | 0.632440681 |
| map04923 | Regulation of lipolysis in adipocytes | 0.659286786 |
| map04723 | Retrograde endocannabinoid signaling | 0.659286786 |
| map02010 | ABC transporters | 0.659286786 |
| map05219 | Bladder cancer | 0.668406492 |

**Table S17.** The detailed lists of GO terms of positive selection genes.

| GO ID | GO Term | GO Class | Adjusted *P*-value |
| --- | --- | --- | --- |
| GO:0006259 | DNA metabolic process | BP | 0.000241508 |
| GO:0044710 | single-organism metabolic process | BP | 0.000256517 |
| GO:0006974 | cellular response to DNA damage stimulus | BP | 0.000837132 |
| GO:0006278 | RNA-dependent DNA replication | BP | 0.000867071 |
| GO:0007020 | microtubule nucleation | BP | 0.000867071 |
| GO:0015980 | energy derivation by oxidation of organic compounds | BP | 0.001303923 |
| GO:0003824 | catalytic activity | MF | 0.001321106 |
| GO:0016788 | hydrolase activity, acting on ester bonds | MF | 0.001925275 |
| GO:0006281 | DNA repair | BP | 0.002063339 |
| GO:0044711 | single-organism biosynthetic process | BP | 0.00266355 |
| GO:1901566 | organonitrogen compound biosynthetic process | BP | 0.00285784 |
| GO:0006091 | generation of precursor metabolites and energy | BP | 0.002980937 |
| GO:0042558 | pteridine-containing compound metabolic process | BP | 0.003516154 |
| GO:0008757 | S-adenosylmethionine-dependent methyltransferase activity | MF | 0.004885117 |
| GO:0051262 | protein tetramerization | BP | 0.005001275 |
| GO:0006188 | IMP biosynthetic process | BP | 0.005001275 |
| GO:0009067 | aspartate family amino acid biosynthetic process | BP | 0.005001275 |
| GO:0005923 | tight junction | CC | 0.005755734 |
| GO:0016787 | hydrolase activity | MF | 0.006585436 |
| GO:1901607 | alpha-amino acid biosynthetic process | BP | 0.007624777 |
| GO:0065009 | regulation of molecular function | BP | 0.007810055 |
| GO:0006099 | tricarboxylic acid cycle | BP | 0.008173319 |
| GO:0004386 | helicase activity | MF | 0.009017416 |
| GO:0016853 | isomerase activity | MF | 0.0096544 |
| GO:0016879 | ligase activity, forming carbon-nitrogen bonds | MF | 0.010453026 |
| GO:0008152 | metabolic process | BP | 0.011086295 |
| GO:0044237 | cellular metabolic process | BP | 0.011929621 |
| GO:0019842 | vitamin binding | MF | 0.012467748 |
| GO:0006310 | DNA recombination | BP | 0.012712124 |
| GO:0030695 | GTPase regulator activity | MF | 0.0129978 |
| GO:0030554 | adenyl nucleotide binding | MF | 0.014339702 |
| GO:0005815 | microtubule organizing center | CC | 0.014926977 |
| GO:0016874 | ligase activity | MF | 0.015422964 |
| GO:0005977 | glycogen metabolic process | BP | 0.016505149 |
| GO:0005524 | ATP binding | MF | 0.016831962 |
| GO:0044093 | positive regulation of molecular function | BP | 0.017345744 |
| GO:0050790 | regulation of catalytic activity | BP | 0.019843661 |
| GO:0043566 | structure-specific DNA binding | MF | 0.019969926 |
| GO:0016835 | carbon-oxygen lyase activity | MF | 0.019969926 |
| GO:0009396 | folic acid-containing compound biosynthetic process | BP | 0.021581968 |
| GO:0008483 | transaminase activity | MF | 0.021581968 |
| GO:0004221 | ubiquitin thiolesterase activity | MF | 0.024229625 |
| GO:0006260 | DNA replication | BP | 0.025940193 |
| GO:0046394 | carboxylic acid biosynthetic process | BP | 0.025940193 |
| GO:0019752 | carboxylic acid metabolic process | BP | 0.027092353 |
| GO:0006298 | mismatch repair | BP | 0.02721384 |
| GO:0030983 | mismatched DNA binding | MF | 0.02721384 |
| GO:0003993 | acid phosphatase activity | MF | 0.02721384 |
| GO:0000922 | spindle pole | CC | 0.02721384 |
| GO:0007088 | regulation of mitosis | BP | 0.02721384 |
| GO:0004520 | endodeoxyribonuclease activity | MF | 0.02721384 |
| GO:0031301 | integral component of organelle membrane | CC | 0.029079604 |
| GO:0004639 | phosphoribosylaminoimidazolesuccinocarboxamide synthase activity | MF | 0.029488066 |
| GO:0006189 | 'de novo' IMP biosynthetic process | BP | 0.029488066 |
| GO:0034023 | 5-(carboxyamino)imidazole ribonucleotide mutase activity | MF | 0.029488066 |
| GO:0000445 | THO complex part of transcription export complex | CC | 0.029488066 |
| GO:0003721 | telomeric template RNA reverse transcriptase activity | MF | 0.029488066 |
| GO:0004347 | glucose-6-phosphate isomerase activity | MF | 0.029488066 |
| GO:0004726 | non-membrane spanning protein tyrosine phosphatase activity | MF | 0.029488066 |
| GO:0003081 | regulation of systemic arterial blood pressure by renin-angiotensin | BP | 0.029488066 |
| GO:0008705 | methionine synthase activity | MF | 0.029488066 |
| GO:0016608 | growth hormone-releasing hormone activity | MF | 0.029488066 |
| GO:0000930 | gamma-tubulin complex | CC | 0.029488066 |
| GO:0031122 | cytoplasmic microtubule organization | BP | 0.029488066 |
| GO:0004168 | dolichol kinase activity | MF | 0.029488066 |
| GO:0043048 | dolichyl monophosphate biosynthetic process | BP | 0.029488066 |
| GO:0007096 | regulation of exit from mitosis | BP | 0.029488066 |
| GO:0004677 | DNA-dependent protein kinase activity | MF | 0.029488066 |
| GO:0045116 | protein neddylation | BP | 0.029488066 |
| GO:0034364 | high-density lipoprotein particle | CC | 0.029488066 |
| GO:0043691 | reverse cholesterol transport | BP | 0.029488066 |
| GO:0005697 | telomerase holoenzyme complex | CC | 0.029488066 |
| GO:0007004 | telomere maintenance via telomerase | BP | 0.029488066 |
| GO:0051973 | positive regulation of telomerase activity | BP | 0.029488066 |
| GO:0004655 | porphobilinogen synthase activity | MF | 0.029488066 |
| GO:0005784 | Sec61 translocon complex | CC | 0.029488066 |
| GO:0004018 | N6-(1,2-dicarboxyethyl)AMP AMP-lyase (fumarate-forming) activity | MF | 0.029488066 |
| GO:0042138 | meiotic DNA double-strand break formation | BP | 0.029488066 |
| GO:0031177 | phosphopantetheine binding | MF | 0.029488066 |
| GO:0003994 | aconitate hydratase activity | MF | 0.029488066 |
| GO:0004135 | amylo-alpha-1,6-glucosidase activity | MF | 0.029488066 |
| GO:0008140 | cAMP response element binding protein binding | MF | 0.029488066 |
| GO:0032793 | positive regulation of CREB transcription factor activity | BP | 0.029488066 |
| GO:0051289 | protein homotetramerization | BP | 0.029488066 |
| GO:0006919 | activation of cysteine-type endopeptidase activity involved in apoptotic process | BP | 0.029488066 |
| GO:0005083 | small GTPase regulator activity | MF | 0.030498634 |
| GO:0044238 | primary metabolic process | BP | 0.031573997 |
| GO:0004519 | endonuclease activity | MF | 0.032526982 |
| GO:0030176 | integral component of endoplasmic reticulum membrane | CC | 0.033363816 |
| GO:0004725 | protein tyrosine phosphatase activity | MF | 0.034503327 |
| GO:0006725 | cellular aromatic compound metabolic process | BP | 0.035465928 |
| GO:0005096 | GTPase activator activity | MF | 0.03746885 |
| GO:0006302 | double-strand break repair | BP | 0.03999663 |
| GO:0006904 | vesicle docking involved in exocytosis | BP | 0.03999663 |
| GO:0051336 | regulation of hydrolase activity | BP | 0.040069568 |
| GO:0016741 | transferase activity, transferring one-carbon groups | MF | 0.040413856 |
| GO:0004721 | phosphoprotein phosphatase activity | MF | 0.04130608 |
| GO:0006950 | response to stress | BP | 0.042578427 |
| GO:0008276 | protein methyltransferase activity | MF | 0.044080027 |
| GO:0000280 | nuclear division | BP | 0.044080027 |
| GO:0071704 | organic substance metabolic process | BP | 0.044508046 |
| GO:0019899 | enzyme binding | MF | 0.044818697 |
| GO:0016702 | oxidoreductase activity, acting on single donors with incorporation of molecular oxygen, incorporation of two atoms of oxygen | MF | 0.047078635 |
| GO:0006511 | ubiquitin-dependent protein catabolic process | BP | 0.047511947 |
| GO:0004674 | protein serine/threonine kinase activity | MF | 0.04762463 |
| GO:1901360 | organic cyclic compound metabolic process | BP | 0.048647121 |
| GO:0006807 | nitrogen compound metabolic process | BP | 0.049325192 |

Note: BP for Biological Process, MF for Molecular Function, and CC for Cellular Component.

**Table S18.** The detailed lists of KEGG pathways of positive selection genes.

| Map ID | Map Title | Adjusted *P*-value |
| --- | --- | --- |
| map00670 | One carbon pool by folate | 0.003598 |
| map05235 | PD-L1 expression and PD-1 checkpoint pathway in cancer | 0.003877 |
| map00350 | Tyrosine metabolism | 0.003881 |
| map03450 | Non-homologous end-joining | 0.004624 |
| map04610 | Complement and coagulation cascades | 0.008854 |
| map00960 | Tropane, piperidine and pyridine alkaloid biosynthesis | 0.012102 |
| map00400 | Phenylalanine, tyrosine and tryptophan biosynthesis | 0.016615 |
| map01210 | 2-Oxocarboxylic acid metabolism | 0.017485 |
| map00360 | Phenylalanine metabolism | 0.026044 |
| map00401 | Novobiocin biosynthesis | 0.029608 |
| map00250 | Alanine, aspartate and glutamate metabolism | 0.029739 |
| map00520 | Amino sugar and nucleotide sugar metabolism | 0.029857 |
| map05150 | Staphylococcus aureus infection | 0.034806 |
| map05340 | Primary immunodeficiency | 0.034806 |
| map04964 | Proximal tubule bicarbonate reclamation | 0.036466 |

**Table S19.** Genomic regions with selection signals during CC duck domestication.

| Chromosome | Start | End | Number of SNPs© | π© | Number of SNPs® | π® | Log2(π®/π© ratio） |
| --- | --- | --- | --- | --- | --- | --- | --- |
| chr1 | 93710001 | 93730000 | 17 | 7.04716E-05 | 108 | 0.0015318 | 4.442042164 |
| chr1 | 99620001 | 99640000 | 7 | 7.53623E-05 | 82 | 0.00117392 | 3.96134729 |
| chr1 | 99650001 | 99670000 | 1 | 4.16667E-06 | 67 | 0.00108144 | 8.019843066 |
| chr1 | 99690001 | 99710000 | 3 | 5.18116E-05 | 104 | 0.00136815 | 4.722807465 |
| chr1 | 99700001 | 99720000 | 2 | 4.76449E-05 | 89 | 0.00115026 | 4.593494395 |
| chr1 | 119610001 | 119630000 | 1 | 4.16667E-06 | 211 | 0.00315376 | 9.563962317 |
| chr1 | 119710001 | 119730000 | 1 | 4.16667E-06 | 140 | 0.00215426 | 9.014081822 |
| chr1 | 119720001 | 119740000 | 1 | 4.16667E-06 | 100 | 0.00142927 | 8.42216792 |
| chr1 | 120130001 | 120150000 | 3 | 1.63043E-05 | 112 | 0.0014652 | 6.489701293 |
| chr1 | 120140001 | 120160000 | 2 | 1.21377E-05 | 142 | 0.00195301 | 7.330060458 |
| chr1 | 120170001 | 120190000 | 4 | 1.66667E-05 | 106 | 0.00145412 | 6.447034042 |
| chr1 | 120180001 | 120200000 | 4 | 1.66667E-05 | 83 | 0.00112493 | 6.076722941 |
| chr1 | 120610001 | 120630000 | 1 | 4.16667E-06 | 53 | 0.000877041 | 7.717605634 |
| chr1 | 120620001 | 120640000 | 1 | 4.16667E-06 | 71 | 0.0012262 | 8.201083751 |
| chr1 | 120630001 | 120650000 | 1 | 4.16667E-06 | 73 | 0.00115536 | 8.115231894 |
| chr1 | 120640001 | 120660000 | 1 | 4.16667E-06 | 49 | 0.000701906 | 7.396239183 |
| chr1 | 121490001 | 121510000 | 5 | 4.29351E-05 | 79 | 0.00118913 | 4.791605082 |
| chr1 | 121520001 | 121540000 | 3 | 0.0000125 | 72 | 0.000921914 | 6.204632176 |
| chr1 | 121530001 | 121550000 | 2 | 8.33333E-06 | 84 | 0.00111495 | 7.063870187 |
| chr1 | 123210001 | 123230000 | 3 | 1.93842E-05 | 76 | 0.000957165 | 5.625814543 |
| chr1 | 123290001 | 123310000 | 3 | 2.73551E-05 | 86 | 0.00128274 | 5.551275139 |
| chr1 | 123300001 | 123320000 | 4 | 3.53261E-05 | 114 | 0.00177985 | 5.654877366 |
| chr1 | 123310001 | 123330000 | 1 | 7.97101E-06 | 135 | 0.00204622 | 8.003983011 |
| chr1 | 123320001 | 123340000 | 1 | 4.16667E-06 | 177 | 0.00272987 | 9.355721691 |
| chr10 | 4740001 | 4760000 | 3 | 2.28261E-05 | 234 | 0.00361828 | 7.308475859 |
| chr10 | 6170001 | 6190000 | 1 | 7.97101E-06 | 124 | 0.00174154 | 7.771385356 |
| chr10 | 6180001 | 6200000 | 2 | 8.33333E-06 | 90 | 0.00118939 | 7.157113024 |
| chr10 | 6200001 | 6220000 | 1 | 2.59058E-05 | 86 | 0.00122549 | 5.563939765 |
| chr10 | 6210001 | 6230000 | 3 | 3.42391E-05 | 114 | 0.001599 | 5.545381349 |
| chr11 | 7440001 | 7460000 | 1 | 4.16667E-06 | 74 | 0.000853645 | 7.678597577 |
| chr12 | 7730001 | 7750000 | 3 | 0.0000125 | 228 | 0.00327417 | 8.033057326 |
| chr12 | 7740001 | 7760000 | 3 | 0.0000125 | 236 | 0.00329538 | 8.042372931 |
| chr13 | 1230001 | 1250000 | 3 | 5.18116E-05 | 113 | 0.00146411 | 4.820605002 |
| chr17 | 3990001 | 4010000 | 3 | 3.15217E-05 | 216 | 0.00321082 | 6.670452636 |
| chr17 | 4000001 | 4020000 | 2 | 8.33333E-06 | 231 | 0.00344656 | 8.692048304 |
| chr21 | 8880001 | 8900000 | 16 | 0.000111596 | 140 | 0.00195903 | 4.133782269 |
| chr21 | 8930001 | 8950000 | 1 | 4.16667E-06 | 100 | 0.00125331 | 8.232632744 |
| chr3 | 59890001 | 59910000 | 10 | 0.000045471 | 101 | 0.00148214 | 5.026591186 |
| chr3 | 72160001 | 72180000 | 5 | 5.99638E-05 | 121 | 0.00178345 | 4.894435147 |
| chr3 | 72170001 | 72190000 | 4 | 5.57971E-05 | 142 | 0.00207064 | 5.213742798 |
| chr3 | 72180001 | 72200000 | 1 | 4.16667E-06 | 165 | 0.00248806 | 9.221910718 |
| chr4 | 22590001 | 22610000 | 56 | 0.000236777 | 214 | 0.0033189 | 3.809104308 |
| chr4 | 41720001 | 41740000 | 1 | 4.16667E-06 | 47 | 0.000616856 | 7.20989509 |
| chr4 | 41760001 | 41780000 | 1 | 4.16667E-06 | 60 | 0.000773551 | 7.536457758 |
| chr5 | 24260001 | 24280000 | 1 | 4.16667E-06 | 228 | 0.00335862 | 9.654758017 |
| chr6 | 8650001 | 8670000 | 1 | 4.16667E-06 | 127 | 0.00187348 | 8.812610018 |
| chr6 | 8660001 | 8680000 | 15 | 0.0000625 | 138 | 0.00204014 | 5.028668157 |
| chr8 | 22640001 | 22660000 | 1 | 4.16667E-06 | 233 | 0.00344909 | 9.693105216 |
| chr8 | 22670001 | 22690000 | 3 | 0.0000125 | 191 | 0.00312826 | 7.96728852 |
| chr9 | 20850001 | 20870000 | 2 | 2.13768E-05 | 163 | 0.00228928 | 6.742704213 |
| chr9 | 20860001 | 20880000 | 1 | 1.72101E-05 | 138 | 0.00194182 | 6.818010184 |

Note: © means CC duck; ® means MDZ; ©® selective sweep analysis between CC duck and MDZ.

**Table S19.** Genomic regions with selection signals during CC duck domestication.

| Chromosome | Start | End | XP-EHH©® | Z trans©® | *P* value©® | Weighted *F*_ST_©® | Mean *F*_ST_©® | Gene Symbol |
| --- | --- | --- | --- | --- | --- | --- | --- | --- |
| chr1 | 93710001 | 93730000 | 4.614959602 | 2.552936255 | 0.005341 | 0.549034 | 0.364078 | *GRIK1* |
| chr1 | 99620001 | 99640000 | 4.896847048 | 2.864210894 | 0.0020902 | 0.541657 | 0.356505 |  |
| chr1 | 99650001 | 99670000 | 4.451170809 | 2.372072233 | 0.0088443 | 0.577516 | 0.419105 | *CYYR1* |
| chr1 | 99690001 | 99710000 | 4.773650064 | 2.728170438 | 0.0031843 | 0.608604 | 0.372808 |  |
| chr1 | 99700001 | 99720000 | 4.762931989 | 2.716334988 | 0.0033005 | 0.605157 | 0.375028 |  |
| chr1 | 119610001 | 119630000 | 5.854145547 | 3.921309209 | 0.000044035 | 0.52391 | 0.354924 | *HSD3B, MAN1A2* |
| chr1 | 119710001 | 119730000 | 5.489364081 | 3.518498723 | 0.000217 | 0.561434 | 0.410942 |  |
| chr1 | 119720001 | 119740000 | 5.391332488 | 3.410247188 | 0.00032452 | 0.625954 | 0.439889 |  |
| chr1 | 120130001 | 120150000 | 4.781707978 | 2.737068401 | 0.0030995 | 0.626106 | 0.347238 | *NANOG* |
| chr1 | 120140001 | 120160000 | 4.946959253 | 2.919547372 | 0.0017527 | 0.654511 | 0.403916 |  |
| chr1 | 120170001 | 120190000 | 4.79893511 | 2.756091488 | 0.0029248 | 0.569044 | 0.355042 | *APOBEC1, AICDA* |
| chr1 | 120180001 | 120200000 | 4.711934416 | 2.66002084 | 0.0039068 | 0.595229 | 0.379735 |  |
| chr1 | 120610001 | 120630000 | 4.924348104 | 2.894578976 | 0.0018983 | 0.582642 | 0.45329 | *ovostatin* |
| chr1 | 120620001 | 120640000 | 5.222300745 | 3.22359363 | 0.00063296 | 0.477672 | 0.387846 |  |
| chr1 | 120630001 | 120650000 | 5.291869832 | 3.3004154 | 0.00048271 | 0.501661 | 0.36239 |  |
| chr1 | 120640001 | 120660000 | 5.03678381 | 3.018736274 | 0.0012692 | 0.542094 | 0.342917 | *H2A1* |
| chr1 | 121490001 | 121510000 | 5.20725205 | 3.206976087 | 0.00067069 | 0.601087 | 0.383777 | *TRPV5* |
| chr1 | 121520001 | 121540000 | 4.517252223 | 2.445042735 | 0.0072417 | 0.60772 | 0.369978 | *EPHB6* |
| chr1 | 121530001 | 121550000 | 4.61651562 | 2.554654491 | 0.0053147 | 0.61826 | 0.385513 |  |
| chr1 | 123210001 | 123230000 | 4.565888189 | 2.498749073 | 0.0062316 | 0.624462 | 0.359539 | *PEX5, KEL, TAS2R40* |
| chr1 | 123290001 | 123310000 | 4.73941052 | 2.690361369 | 0.0035687 | 0.593641 | 0.408311 | *CD163* |
| chr1 | 123300001 | 123320000 | 4.840131101 | 2.801582224 | 0.0025426 | 0.490058 | 0.359607 |  |
| chr1 | 123310001 | 123330000 | 5.003944083 | 2.982472956 | 0.0014296 | 0.556637 | 0.385726 | *GSTK1* |
| chr1 | 123320001 | 123340000 | 5.10432964 | 3.09332386 | 0.00098964 | 0.553316 | 0.38226 |  |
| chr10 | 4740001 | 4760000 | 4.807164298 | 2.765178581 | 0.0028446 | 0.521633 | 0.356489 | *LRRC31, SAMD7* |
| chr10 | 6170001 | 6190000 | 4.874392265 | 2.839415165 | 0.0022598 | 0.572136 | 0.396231 |  |
| chr10 | 6180001 | 6200000 | 4.525525458 | 2.454178467 | 0.0070603 | 0.598909 | 0.41119 |  |
| chr10 | 6200001 | 6220000 | 4.566340352 | 2.499248375 | 0.0062229 | 0.575252 | 0.404026 | *NAALADL2* |
| chr10 | 6210001 | 6230000 | 4.757445359 | 2.710276368 | 0.0033614 | 0.559528 | 0.366251 |  |
| chr11 | 7440001 | 7460000 | 4.792130938 | 2.74857797 | 0.0029927 | 0.540109 | 0.345487 | *Synpr* |
| chr12 | 7730001 | 7750000 | 4.558058542 | 2.490103174 | 0.0063853 | 0.533013 | 0.339569 | *NTRK3* |
| chr12 | 7740001 | 7760000 | 4.428740676 | 2.347303725 | 0.0094549 | 0.550733 | 0.35281 |  |
| chr13 | 1230001 | 1250000 | 4.527607754 | 2.456477845 | 0.0070153 | 0.555005 | 0.374041 | *TCF25, ATM* |
| chr17 | 3990001 | 4010000 | 4.618830801 | 2.557211032 | 0.0052758 | 0.53919 | 0.371941 |  |
| chr17 | 4000001 | 4020000 | 4.752517044 | 2.704834269 | 0.0034169 | 0.524845 | 0.361803 |  |
| chr21 | 8880001 | 8900000 | 5.519404605 | 3.551671018 | 0.0001914 | 0.583491 | 0.396283 |  |
| chr21 | 8930001 | 8950000 | 5.223263176 | 3.224656397 | 0.00063062 | 0.610886 | 0.411402 |  |
| chr3 | 59890001 | 59910000 | 4.505267547 | 2.431808637 | 0.0075118 | 0.611223 | 0.415467 | *Stx7* |
| chr3 | 72160001 | 72180000 | 4.650712974 | 2.592416971 | 0.0047652 | 0.50288 | 0.354102 |  |
| chr3 | 72170001 | 72190000 | 4.718580258 | 2.667359521 | 0.0038225 | 0.567818 | 0.399021 |  |
| chr3 | 72180001 | 72200000 | 4.7360917 | 2.686696558 | 0.0036081 | 0.535557 | 0.37114 |  |
| chr4 | 22590001 | 22610000 | 4.800999155 | 2.758370712 | 0.0029045 | 0.573005 | 0.433108 | *RFC1* |
| chr4 | 41720001 | 41740000 | 4.679203675 | 2.62387787 | 0.0043467 | 0.587163 | 0.347903 |  |
| chr4 | 41760001 | 41780000 | 4.522632401 | 2.450983804 | 0.0071233 | 0.630026 | 0.368391 |  |
| chr5 | 24260001 | 24280000 | 4.984910919 | 2.961455556 | 0.0015309 | 0.50425 | 0.343862 | *FCF1, Arel1* |
| chr6 | 8650001 | 8670000 | 4.66909461 | 2.61271492 | 0.0044913 | 0.527895 | 0.342137 |  |
| chr6 | 8660001 | 8680000 | 4.695807507 | 2.642212677 | 0.0041183 | 0.561754 | 0.380586 |  |
| chr8 | 22640001 | 22660000 | 5.072790563 | 3.058496786 | 0.0011123 | 0.510208 | 0.34986 | *DNM3* |
| chr8 | 22670001 | 22690000 | 4.768321976 | 2.722286888 | 0.0032416 | 0.54661 | 0.407495 | *PIGC* |
| chr9 | 20850001 | 20870000 | 4.73800882 | 2.68881354 | 0.0035853 | 0.721273 | 0.554048 | *DIP2C* |
| chr9 | 20860001 | 20880000 | 4.917055812 | 2.886526452 | 0.0019476 | 0.644851 | 0.454897 |  |

Note: © means CC duck; ® means mallard in Zhejiang Province (MDZ); ©® selective sweep analysis between CC duck and MDZ.

**Table S20.** The detailed lists of GO terms of SV relate and species-specific genes compared to Pekin duck.

| Structure variants type | GO ID | GO Term | GO Class | Adjusted *P*-value |
| --- | --- | --- | --- | --- |
| SV relate genes | GO:0007155 | cell adhesion | BP | 3.28E-05 |
|  | GO:0022610 | biological adhesion | BP | 3.28E-05 |
|  | GO:0004713 | protein tyrosine kinase activity | MF | 0.000216 |
|  | GO:0023052 | signaling | BP | 0.000576 |
|  | GO:0044700 | single organism signaling | BP | 0.000576 |
|  | GO:0007156 | homophilic cell adhesion | BP | 0.000576 |
|  | GO:0098609 | cell-cell adhesion | BP | 0.000576 |
|  | GO:0007154 | cell communication | BP | 0.000976 |
|  | GO:0007165 | signal transduction | BP | 0.001052 |
|  | GO:0016301 | kinase activity | MF | 0.004411 |
|  | GO:0005201 | extracellular matrix structural constituent | MF | 0.004411 |
|  | GO:0051716 | cellular response to stimulus | BP | 0.004487 |
|  | GO:0004672 | protein kinase activity | MF | 0.004487 |
|  | GO:0006468 | protein phosphorylation | BP | 0.004682 |
|  | GO:0044763 | single-organism cellular process | BP | 0.005758 |
|  | GO:0065007 | biological regulation | BP | 0.006621 |
|  | GO:0050794 | regulation of cellular process | BP | 0.00673 |
|  | GO:0050789 | regulation of biological process | BP | 0.007814 |
|  | GO:0016310 | phosphorylation | BP | 0.007814 |
|  | GO:0010646 | regulation of cell communication | BP | 0.007814 |
|  | GO:0009966 | regulation of signal transduction | BP | 0.010292 |
|  | GO:0023051 | regulation of signaling | BP | 0.011892 |
|  | GO:0007169 | transmembrane receptor protein tyrosine kinase signaling pathway | BP | 0.012947 |
|  | GO:0030554 | adenyl nucleotide binding | MF | 0.020111 |
|  | GO:0016773 | phosphotransferase activity, alcohol group as acceptor | MF | 0.022417 |
|  | GO:0005581 | collagen trimer | CC | 0.022417 |
|  | GO:0048583 | regulation of response to stimulus | BP | 0.022417 |
|  | GO:0005126 | cytokine receptor binding | MF | 0.034571 |
|  | GO:0007167 | enzyme linked receptor protein signaling pathway | BP | 0.034571 |
|  | GO:0005524 | ATP binding | MF | 0.034571 |
|  | GO:0032559 | adenyl ribonucleotide binding | MF | 0.035892 |
| Species-specific genes | GO:0005200 | structural constituent of cytoskeleton | MF | 2.81E-05 |
|  | GO:0005882 | intermediate filament | CC | 6.58E-05 |
|  | GO:0045111 | intermediate filament cytoskeleton | CC | 6.58E-05 |
|  | GO:0005149 | interleukin-1 receptor binding | MF | 0.025075 |

Note: BP for Biological Process, MF for Molecular Function, and CC for Cellular Component.

**Table S21.** The detailed lists of GO terms of the SV and specie specific genes compared to Csp-b duck.

| Structure Variant type | GO ID | GO Term | GO Class | Adjusted *P*-value |
| --- | --- | --- | --- | --- |
| SVs | GO:0051056 | regulation of small GTPase mediated signal transduction | BP | 5.51E-09 |
|  | GO:0005085 | guanyl-nucleotide exchange factor activity | MF | 9.43E-09 |
|  | GO:1902531 | regulation of intracellular signal transduction | BP | 1.36E-08 |
|  | GO:0046578 | regulation of Ras protein signal transduction | BP | 4.10E-06 |
|  | GO:0007265 | Ras protein signal transduction | BP | 1.01E-05 |
|  | GO:0010646 | regulation of cell communication | BP | 1.01E-05 |
|  | GO:0005515 | protein binding | MF | 1.01E-05 |
|  | GO:0009966 | regulation of signal transduction | BP | 1.29E-05 |
|  | GO:0023051 | regulation of signaling | BP | 1.67E-05 |
|  | GO:0030529 | ribonucleoprotein complex | CC | 2.21E-05 |
|  | GO:0005089 | Rho guanyl-nucleotide exchange factor activity | MF | 2.21E-05 |
|  | GO:0035023 | regulation of Rho protein signal transduction | BP | 2.21E-05 |
|  | GO:0005088 | Ras guanyl-nucleotide exchange factor activity | MF | 2.21E-05 |
|  | GO:0007266 | Rho protein signal transduction | BP | 2.21E-05 |
|  | GO:0043167 | ion binding | MF | 2.95E-05 |
|  | GO:0048583 | regulation of response to stimulus | BP | 9.49E-05 |
|  | GO:0044763 | single-organism cellular process | BP | 0.000102 |
|  | GO:0005201 | extracellular matrix structural constituent | MF | 0.000533 |
|  | GO:0003735 | structural constituent of ribosome | MF | 0.00075 |
|  | GO:0005488 | binding | MF | 0.001169 |
|  | GO:0005840 | ribosome | CC | 0.001169 |
|  | GO:0035556 | intracellular signal transduction | BP | 0.001194 |
|  | GO:0055085 | transmembrane transport | BP | 0.001503 |
|  | GO:0007264 | small GTPase mediated signal transduction | BP | 0.001646 |
|  | GO:0006412 | translation | BP | 0.001692 |
|  | GO:0005543 | phospholipid binding | MF | 0.00296 |
|  | GO:0004672 | protein kinase activity | MF | 0.00371 |
|  | GO:0004114 | 3',5'-cyclic-nucleotide phosphodiesterase activity | MF | 0.004419 |
|  | GO:0043168 | anion binding | MF | 0.004419 |
|  | GO:0004713 | protein tyrosine kinase activity | MF | 0.0062 |
|  | GO:0005578 | proteinaceous extracellular matrix | CC | 0.006648 |
|  | GO:0023052 | signaling | BP | 0.007749 |
|  | GO:0044700 | single organism signaling | BP | 0.007749 |
|  | GO:0004112 | cyclic-nucleotide phosphodiesterase activity | MF | 0.008113 |
|  | GO:0030554 | adenyl nucleotide binding | MF | 0.008113 |
|  | GO:0065007 | biological regulation | BP | 0.011467 |
|  | GO:0005216 | ion channel activity | MF | 0.011467 |
|  | GO:0015267 | channel activity | MF | 0.011467 |
|  | GO:0022803 | passive transmembrane transporter activity | MF | 0.011467 |
|  | GO:0022838 | substrate-specific channel activity | MF | 0.011467 |
|  | GO:0050794 | regulation of cellular process | BP | 0.01199 |
|  | GO:0005524 | ATP binding | MF | 0.013964 |
|  | GO:0015085 | calcium ion transmembrane transporter activity | MF | 0.014364 |
|  | GO:0032559 | adenyl ribonucleotide binding | MF | 0.014364 |
|  | GO:0050789 | regulation of biological process | BP | 0.014623 |
|  | GO:0007167 | enzyme linked receptor protein signaling pathway | BP | 0.014623 |
|  | GO:0006468 | protein phosphorylation | BP | 0.015595 |
|  | GO:0044765 | single-organism transport | BP | 0.016342 |
|  | GO:0005581 | collagen trimer | CC | 0.016892 |
|  | GO:0016301 | kinase activity | MF | 0.019124 |
|  | GO:0007165 | signal transduction | BP | 0.019124 |
|  | GO:0007269 | neurotransmitter secretion | BP | 0.022766 |
|  | GO:0051716 | cellular response to stimulus | BP | 0.022766 |
|  | GO:0031012 | extracellular matrix | CC | 0.022766 |
|  | GO:0008081 | phosphoric diester hydrolase activity | MF | 0.025806 |
|  | GO:0005509 | calcium ion binding | MF | 0.025806 |
|  | GO:0023061 | signal release | BP | 0.025806 |
|  | GO:0007154 | cell communication | BP | 0.025806 |
|  | GO:0043169 | cation binding | MF | 0.026076 |
|  | GO:0016310 | phosphorylation | BP | 0.026979 |
|  | GO:0046872 | metal ion binding | MF | 0.027798 |
|  | GO:0016773 | phosphotransferase activity, alcohol group as acceptor | MF | 0.027798 |
|  | GO:0008289 | lipid binding | MF | 0.029614 |
|  | GO:0044699 | single-organism process | BP | 0.035933 |
|  | GO:0005096 | GTPase activator activity | MF | 0.039933 |
|  | GO:0022857 | transmembrane transporter activity | MF | 0.042602 |
|  | GO:0015075 | ion transmembrane transporter activity | MF | 0.046845 |
|  | GO:0005102 | receptor binding | MF | 0.049538 |
|  | GO:0006955 | immune response | BP | 0.049538 |
|  | GO:0017048 | Rho GTPase binding | MF | 0.049538 |
|  | GO:0044463 | cell projection part | CC | 0.049538 |
|  | GO:0016491 | oxidoreductase activity | MF | 0.049538 |
|  | GO:0008528 | G-protein coupled peptide receptor activity | MF | 0.049538 |
|  | GO:0001653 | peptide receptor activity | MF | 0.049538 |
| Species-specific genes | GO:0000786 | nucleosome | CC | 9.94E-23 |
|  | GO:1990104 | DNA bending complex | CC | 9.94E-23 |
|  | GO:0044815 | DNA packaging complex | CC | 2.01E-22 |
|  | GO:0006334 | nucleosome assembly | BP | 6.62E-22 |
|  | GO:0006333 | chromatin assembly or disassembly | BP | 6.62E-22 |
|  | GO:0034728 | nucleosome organization | BP | 6.62E-22 |
|  | GO:0031497 | chromatin assembly | BP | 6.62E-22 |
|  | GO:0000785 | chromatin | CC | 6.62E-22 |
|  | GO:0006323 | DNA packaging | BP | 3.37E-21 |
|  | GO:0032993 | protein-DNA complex | CC | 6.03E-21 |
|  | GO:0065004 | protein-DNA complex assembly | BP | 6.03E-21 |
|  | GO:0071824 | protein-DNA complex subunit organization | BP | 6.03E-21 |
|  | GO:0071103 | DNA conformation change | BP | 4.21E-20 |
|  | GO:0006325 | chromatin organization | BP | 8.31E-18 |
|  | GO:0044427 | chromosomal part | CC | 1.43E-17 |
|  | GO:0034622 | cellular macromolecular complex assembly | BP | 2.23E-17 |
|  | GO:0005694 | chromosome | CC | 3.51E-16 |
|  | GO:0006996 | organelle organization | BP | 1.47E-15 |
|  | GO:0051276 | chromosome organization | BP | 3.77E-15 |
|  | GO:0006461 | protein complex assembly | BP | 2.78E-14 |
|  | GO:0070271 | protein complex biogenesis | BP | 3.11E-14 |
|  | GO:0065003 | macromolecular complex assembly | BP | 8.78E-14 |
|  | GO:0043228 | non-membrane-bounded organelle | CC | 3.03E-13 |
|  | GO:0043232 | intracellular non-membrane-bounded organelle | CC | 3.03E-13 |
|  | GO:0071822 | protein complex subunit organization | BP | 4.99E-13 |
|  | GO:0043933 | macromolecular complex subunit organization | BP | 1.39E-12 |
|  | GO:0022607 | cellular component assembly | BP | 2.53E-12 |
|  | GO:0044085 | cellular component biogenesis | BP | 9.81E-12 |
|  | GO:0044422 | organelle part | CC | 1.47E-11 |
|  | GO:0044446 | intracellular organelle part | CC | 1.47E-11 |
|  | GO:0006259 | DNA metabolic process | BP | 8.89E-10 |
|  | GO:0016043 | cellular component organization | BP | 1.7E-09 |
|  | GO:0071840 | cellular component organization or biogenesis | BP | 2.33E-09 |
|  | GO:0043234 | protein complex | CC | 3.21E-08 |
|  | GO:0032991 | macromolecular complex | CC | 2.2E-06 |
|  | GO:0006352 | DNA-templated transcription, initiation | BP | 0.000478 |
|  | GO:0002376 | immune system process | BP | 0.000606 |
|  | GO:0006955 | immune response | BP | 0.001509 |
|  | GO:0003677 | DNA binding | MF | 0.001992 |
|  | GO:0005871 | kinesin complex | CC | 0.004441 |
|  | GO:0019882 | antigen processing and presentation | BP | 0.011673 |
|  | GO:0003777 | microtubule motor activity | MF | 0.021309 |
|  | GO:0043229 | intracellular organelle | CC | 0.021764 |
|  | GO:0005126 | cytokine receptor binding | MF | 0.021764 |
|  | GO:0043226 | organelle | CC | 0.021764 |
|  | GO:0006928 | cellular component movement | BP | 0.022012 |
|  | GO:0030246 | carbohydrate binding | MF | 0.02287 |
|  | GO:0007018 | microtubule-based movement | BP | 0.023053 |
|  | GO:0008009 | chemokine activity | MF | 0.033732 |
|  | GO:0042379 | chemokine receptor binding | MF | 0.033732 |
|  | GO:0015197 | peptide transporter activity | MF | 0.035693 |
|  | GO:0015833 | peptide transport | BP | 0.035693 |
|  | GO:0005149 | interleukin-1 receptor binding | MF | 0.035693 |
|  | GO:0005875 | microtubule associated complex | CC | 0.0465 |

Note: BP for Biological Process, MF for Molecular Function, and CC for Cellular Component.

**Table S22.** Genomic regions identified as candidate divergent regions (CDRs) of crested cushion.

| Chromosome | Start | End | Number of variants© | π© | Number of variants $ | π$ | log2(π$/π© Ratio) | Weighted *F*_ST_©$ | Mean *F*_ST_©$ |
| --- | --- | --- | --- | --- | --- | --- | --- | --- | --- |
| 1 | 99650001 | 99670000 | 1 | 0.00000417 | 73 | 0.001146 | 8.103899 | 0.39605 | 0.277445 |
| 1 | 99680001 | 99700000 | 1 | 0.00000417 | 73 | 0.000911 | 7.771988 | 0.335273 | 0.186999 |
| 1 | 99690001 | 99710000 | 3 | 0.0000518 | 116 | 0.001481 | 4.836763 | 0.379239 | 0.213689 |
| 1 | 99700001 | 99720000 | 2 | 0.0000476 | 94 | 0.001255 | 4.719187 | 0.387194 | 0.229381 |
| 1 | 99760001 | 99780000 | 7 | 0.000111413 | 74 | 0.00105 | 3.236496 | 0.37696 | 0.238431 |
| 1 | 117490001 | 117510000 | 3 | 0.0000125 | 204 | 0.002679 | 7.743456 | 0.428815 | 0.254057 |
| 1 | 119600001 | 119620000 | 13 | 0.000270478 | 238 | 0.003415 | 3.658488 | 0.365198 | 0.233874 |
| 1 | 119610001 | 119630000 | 1 | 0.00000417 | 219 | 0.003226 | 9.59677 | 0.383849 | 0.251398 |
| 1 | 119710001 | 119730000 | 1 | 0.00000417 | 142 | 0.002252 | 9.077975 | 0.465468 | 0.335629 |
| 1 | 119720001 | 119740000 | 1 | 0.00000417 | 104 | 0.001565 | 8.553347 | 0.517667 | 0.366812 |
| 1 | 120140001 | 120160000 | 2 | 0.0000121 | 156 | 0.00224 | 7.527892 | 0.48086 | 0.316481 |
| 1 | 120170001 | 120190000 | 4 | 0.0000167 | 126 | 0.001646 | 6.626185 | 0.449769 | 0.28583 |
| 1 | 120180001 | 120200000 | 4 | 0.0000167 | 97 | 0.001297 | 6.282011 | 0.470929 | 0.28203 |
| 1 | 120610001 | 120630000 | 1 | 0.00000417 | 55 | 0.000926 | 7.795589 | 0.352487 | 0.209797 |
| 1 | 121490001 | 121510000 | 5 | 0.0000429 | 84 | 0.001141 | 4.731884 | 0.538937 | 0.33581 |
| 1 | 121500001 | 121520000 | 5 | 0.0000239 | 94 | 0.001289 | 5.75276 | 0.459066 | 0.270155 |
| 1 | 122590001 | 122610000 | 11 | 0.00025779 | 271 | 0.003922 | 3.927424 | 0.333098 | 0.218541 |
| 1 | 123210001 | 123230000 | 3 | 0.0000194 | 85 | 0.001075 | 5.793956 | 0.487325 | 0.283651 |
| 1 | 123310001 | 123330000 | 1 | 0.00000797 | 154 | 0.002128 | 8.060391 | 0.432749 | 0.266031 |
| 1 | 123320001 | 123340000 | 1 | 0.00000417 | 193 | 0.002706 | 9.343147 | 0.43426 | 0.272297 |
| 1 | 123330001 | 123350000 | 11 | 0.00023442 | 163 | 0.002133 | 3.185493 | 0.382792 | 0.22851 |
| 1 | 138070001 | 138090000 | 65 | 0.000308877 | 175 | 0.002381 | 2.946657 | 0.341405 | 0.203815 |
| 1 | 138080001 | 138100000 | 55 | 0.000248188 | 137 | 0.001902 | 2.937648 | 0.358867 | 0.225839 |
| 2 | 76060001 | 76080000 | 22 | 0.0000917 | 72 | 0.00092 | 3.327711 | 0.344673 | 0.192218 |
| 4 | 22590001 | 22610000 | 56 | 0.000236777 | 221 | 0.003552 | 3.906832 | 0.39329 | 0.284299 |
| 4 | 22600001 | 22620000 | 47 | 0.000286234 | 167 | 0.002531 | 3.144475 | 0.413748 | 0.286462 |

Note: © means CC duck; $ means the duck breeds without crested cushion; ©$ means selective sweep analysis between CC duck and duck breeds without crested cushion; ℗ means Pekin duck; ® means mallard in Zhejiang Province (MDZ); ©℗ means selective sweep analysis between CC duck and Pekin duck; ©® means selective sweep analysis between CC duck and MDZ.

**Table S22.** Genomic regions identified as CDRs of crested cushion.

| Chromosome | Start | End | XP-EHH©℗ | Z trans©℗ | *P* value©℗ | XP-EHH©® | Z trans©® | *P* value©® | Gene symbol |
| --- | --- | --- | --- | --- | --- | --- | --- | --- | --- |
| 1 | 99650001 | 99670000 | 2.901092 | 2.7022478 | 0.003444 | 4.4511708 | 2.3720722 | 0.008844 | *CYYR1* |
| 1 | 99680001 | 99700000 | 3.4102689 | 3.4275377 | 0.000305 | 4.4780479 | 2.4017513 | 0.008158 |  |
| 1 | 99690001 | 99710000 | 3.2914101 | 3.258231 | 0.000561 | 4.7736501 | 2.7281704 | 0.003184 |  |
| 1 | 99700001 | 99720000 | 2.9619411 | 2.7889234 | 0.002644 | 4.762932 | 2.716335 | 0.003301 |  |
| 1 | 99760001 | 99780000 | 3.503551 | 3.560412 | 0.000185 | 4.6156108 | 2.5536554 | 0.00533 | *APP* |
| 1 | 117490001 | 117510000 | 2.9438583 | 2.7631657 | 0.002862 | 4.722908 | 2.6721384 | 0.003769 |  |
| 1 | 119600001 | 119620000 | 4.1114636 | 4.4263446 | 4.79E-06 | 5.801269 | 3.8629202 | 5.6E-05 | *HSD3B* |
| 1 | 119610001 | 119630000 | 3.8649439 | 4.0751931 | 2.3E-05 | 5.8541455 | 3.9213092 | 4.4E-05 | *MAN1A2* |
| 1 | 119710001 | 119730000 | 3.9195162 | 4.152928 | 1.64E-05 | 5.4893641 | 3.5184987 | 0.000217 |  |
| 1 | 119720001 | 119740000 | 3.786283 | 3.9631457 | 3.7E-05 | 5.3913325 | 3.4102472 | 0.000325 |  |
| 1 | 120140001 | 120160000 | 2.052574 | 1.4935882 | 0.006764 | 4.9469593 | 2.9195474 | 0.001753 | *NANOG* |
| 1 | 120170001 | 120190000 | 2.0551079 | 1.4971977 | 0.006717 | 4.7989351 | 2.7560915 | 0.002925 |  |
| 1 | 120180001 | 120200000 | 2.0173132 | 1.4433616 | 0.074459 | 4.7119344 | 2.6600208 | 0.003907 | *APOBEC1*, *AICDA* |
| 1 | 120610001 | 120630000 | 0.6467571 | -0.508908 | 0.69459 | 4.9243481 | 2.894579 | 0.001898 |  |
| 1 | 121490001 | 121510000 | 3.2416703 | 3.1873798 | 0.000718 | 5.2072521 | 3.2069761 | 0.000671 | *TRPV5* |
| 1 | 121500001 | 121520000 | 3.174076 | 3.0910961 | 0.000997 | 5.3161213 | 3.3271951 | 0.000439 | *EPHB6* |
| 1 | 122590001 | 122610000 | 3.1451307 | 3.0498653 | 0.001145 | 4.5907237 | 2.5261737 | 0.005766 | *SCNN1A*, *VAMP1* |
| 1 | 123210001 | 123230000 | 2.6587552 | 2.3570546 | 0.00921 | 4.5658882 | 2.4987491 | 0.006232 | *PEX5*, *KEL*, TAS2R40 |
| 1 | 123310001 | 123330000 | 2.6575756 | 2.3553742 | 0.009252 | 5.0039441 | 2.982473 | 0.00143 | *CD163* |
| 1 | 123320001 | 123340000 | 3.1372461 | 3.0386342 | 0.001188 | 5.1043296 | 3.0933239 | 0.00099 | *GSTK1* |
| 1 | 123330001 | 123350000 | 3.253112 | 3.2036777 | 0.000678 | 4.8191395 | 2.7784022 | 0.002731 |  |
| 1 | 138070001 | 138090000 | 3.1681649 | 3.0826761 | 0.001026 | 5.2113404 | 3.2114907 | 0.00066 | *SPIC* |
| 1 | 138080001 | 138100000 | 3.5085702 | 3.5675616 | 0.00018 | 4.9801636 | 2.9562133 | 0.001557 |  |
| 2 | 76060001 | 76080000 | 3.1119951 | 3.0026657 | 0.001338 | 4.6305274 | 2.5701271 | 0.005083 |  |
| 4 | 22590001 | 22610000 | 3.7270878 | 3.878826 | 5.25E-05 | 4.8009992 | 2.7583707 | 0.002905 | *RFC1* |
| 4 | 22600001 | 22620000 | 3.5981993 | 3.6952326 | 0.00011 | 4.4274257 | 2.3458517 | 0.009492 |  |

Note: © means CC duck; $ means the duck breeds without crested cushion; ©$ means selective sweep analysis between CC duck and duck breeds without crested cushion; ℗ means Pekin duck; ® means mallard in Zhejiang Province (MDZ); ©℗ means selective sweep analysis between CC duck and Pekin duck; ©® means selective sweep analysis between CC duck and MDZ.

**Table S23.** Genomic selective regions identified of crested cushion.

| Chromosome | Start | End | Number of variants© | π© | Number of variants$ | π$ | log2(π$/π© ratio) | Weighted *F*_ST_ ©$ | Mean *F*_ST_ ©$ |
| --- | --- | --- | --- | --- | --- | --- | --- | --- | --- |
| 1 | 120140001 | 120160000 | 2 | 0.0000121 | 156 | 0.00224 | 7.527892 | 0.48086 | 0.316481 |
| 1 | 120170001 | 120190000 | 4 | 0.0000167 | 126 | 0.001646 | 6.626185 | 0.449769 | 0.28583 |
| 1 | 120180001 | 120200000 | 4 | 0.0000167 | 97 | 0.001297 | 6.282011 | 0.470929 | 0.28203 |
| 1 | 121490001 | 121510000 | 5 | 0.0000429 | 84 | 0.001141 | 4.731884 | 0.538937 | 0.33581 |
| 1 | 121500001 | 121520000 | 5 | 0.0000239 | 94 | 0.001289 | 5.75276 | 0.459066 | 0.270155 |
| 1 | 122590001 | 122610000 | 11 | 0.0002578 | 271 | 0.003922 | 3.927424 | 0.333098 | 0.218541 |
| 1 | 123210001 | 123230000 | 3 | 0.0000194 | 85 | 0.001075 | 5.793956 | 0.487325 | 0.283651 |
| 1 | 123310001 | 123330000 | 1 | 7.97E-06 | 154 | 0.002128 | 8.060391 | 0.432749 | 0.266031 |
| 1 | 123320001 | 123340000 | 1 | 4.17E-06 | 193 | 0.002706 | 9.343147 | 0.43426 | 0.272297 |
| 1 | 123330001 | 123350000 | 11 | 0.0002344 | 163 | 0.002133 | 3.185493 | 0.382792 | 0.22851 |
| 1 | 138070001 | 138090000 | 65 | 0.0003089 | 175 | 0.002381 | 2.946657 | 0.341405 | 0.203815 |
| 1 | 138080001 | 138100000 | 55 | 0.0002482 | 137 | 0.001902 | 2.937648 | 0.358867 | 0.225839 |

Note: © means CC duck; $ means the duck breeds without crested cushion; ©$ means selective sweep analysis between CC duck and duck breeds without crested cushion; ℗ means Pekin duck; ® means mallard in Zhejiang Province (MDZ); ©℗ means selective sweep analysis between CC duck and Pekin duck; ©® means selective sweep analysis between CC duck and MDZ; * means crested duck in F_2_ population; ™ means ducks without crested cushion in F_2_ population; *™ means selective sweep analysis between crested duck and normal duck in F_2_ population.

**Table S23.** Genomic selective regions identified of crested cushion.

| Chromosome | Start | End | XP-EHH©℗ | Z trans©℗ | *P* value©℗ | XP-EHH©® | Z trans©® | *P* value©® |
| --- | --- | --- | --- | --- | --- | --- | --- | --- |
| 1 | 120140001 | 120160000 | 2.052574 | 1.493588 | 0.006764 | 4.946959 | 2.919547 | 0.001753 |
| 1 | 120170001 | 120190000 | 2.055108 | 1.497198 | 0.006717 | 4.798935 | 2.756091 | 0.002925 |
| 1 | 120180001 | 120200000 | 2.017313 | 1.443362 | 0.074459 | 4.711934 | 2.660021 | 0.003907 |
| 1 | 121490001 | 121510000 | 3.24167 | 3.18738 | 0.000718 | 5.207252 | 3.206976 | 0.000671 |
| 1 | 121500001 | 121520000 | 3.174076 | 3.091096 | 0.000997 | 5.316121 | 3.327195 | 0.000439 |
| 1 | 122590001 | 122610000 | 3.145131 | 3.049865 | 0.001145 | 4.590724 | 2.526174 | 0.005766 |
| 1 | 123210001 | 123230000 | 2.658755 | 2.357055 | 0.00921 | 4.565888 | 2.498749 | 0.006232 |
| 1 | 123310001 | 123330000 | 2.657576 | 2.355374 | 0.009252 | 5.003944 | 2.982473 | 0.00143 |
| 1 | 123320001 | 123340000 | 3.137246 | 3.038634 | 0.001188 | 5.10433 | 3.093324 | 0.00099 |
| 1 | 123330001 | 123350000 | 3.253112 | 3.203678 | 0.000678 | 4.81914 | 2.778402 | 0.002731 |
| 1 | 138070001 | 138090000 | 3.168165 | 3.082676 | 0.001026 | 5.21134 | 3.211491 | 0.00066 |
| 1 | 138080001 | 138100000 | 3.50857 | 3.567562 | 0.00018 | 4.980164 | 2.956213 | 0.001557 |

Note: © means CC duck; $ means the duck breeds without crested cushion; ©$ means selective sweep analysis between CC duck and duck breeds without crested cushion; ℗ means Pekin duck; ® means MDZ; ©℗ means selective sweep analysis between CC duck and Pekin duck; ©® means selective sweep analysis between CC duck and MDZ; * means crested duck in F_2_ population; ™ means ducks without crested cushion in F_2_ population; *™ means selective sweep analysis between crested duck and normal duck in F_2_ population.

**Table S23.** Genomic selective regions identified of crested cushion.

| Chromosome | Start | End | Weighted *F*_ST_*™ | Mean *F*_ST_*™ | π* | π™ | log2(π™/π* ratio) | Gene symbol |
| --- | --- | --- | --- | --- | --- | --- | --- | --- |
| 1 | 120140001 | 120160000 | 0.449613 | 0.429441 | 0.000306 | 0.003943 | 3.6862 | *NANOG* |
| 1 | 120170001 | 120190000 | 0.447862 | 0.419648 | 0.000229 | 0.002561 | 3.48178 |  |
| 1 | 120180001 | 120200000 | 0.446388 | 0.442783 | 0.000199 | 0.002553 | 3.684883 | *APOBEC1, AICDA* |
| 1 | 121490001 | 121510000 | 0.352885 | 0.261761 | 0.000439 | 0.003913 | 3.156291 | *TRPV5* |
| 1 | 121500001 | 121520000 | 0.340916 | 0.254526 | 0.000418 | 0.003985 | 3.253655 | *EPHB6* |
| 1 | 122590001 | 122610000 | 0.138596 | 0.132862 | 0.001285 | 0.005385 | 2.067048 | *SCNN1A, VAMP1* |
| 1 | 123210001 | 123230000 | 0.31345 | 0.25939 | 0.000245 | 0.003367 | 3.777666 | *PEX5, KEL, TAS2R40* |
| 1 | 123310001 | 123330000 | 0.482355 | 0.422543 | 0.000267 | 0.00256 | 3.259453 | *CD163* |
| 1 | 123320001 | 123340000 | 0.438095 | 0.401394 | 0.00021 | 0.003485 | 4.056036 | *GSTK1* |
| 1 | 123330001 | 123350000 | 0.332086 | 0.283397 | 0.00081 | 0.003309 | 2.029823 |  |
| 1 | 138070001 | 138090000 | 0.256135 | 0.222041 | 0.000472 | 0.004037 | 3.095785 | *SPIC* |
| 1 | 138080001 | 138100000 | 0.268084 | 0.258634 | 0.000437 | 0.003704 | 3.084839 |  |

Note: © means CC duck; $ means the duck breeds without crested cushion; ©$ means selective sweep analysis between CC duck and duck breeds without crested cushion; ℗ means Pekin duck; ® means mallard in Zhejiang Province (MDZ); ©℗ means selective sweep analysis between CC duck and Pekin duck; ©® means selective sweep analysis between CC duck and MDZ; * means crested duck in F_2_ population; ™ means ducks without crested cushion in F_2_ population; *™ means selective sweep analysis between crested duck and normal duck in F_2_ population.

**Table S25. calibration points of 13 species.**

| Species 1 | Species 2 | Min calibration point (Mya) | Max calibration point(Mya) |
| --- | --- | --- | --- |
| *Struthio camelus* | Other species | 105 | 118 |
| *Taeniopygia guttata* | *Anser cygnoides domesticus* | 92 | 104 |
| *Aptenodytes forsteri* | *Egretta garzetta* | 73 | 84 |
| *Nestor notabilis* | *Egretta garzetta* | 71 | 91 |
| *Nestor notabilis* | *Columba livia* | 77 | 90 |
| *Aptenodytes forsteri* | *Balearica regulorum* | 71 | 91 |
| *Coturnix japonica* | *Anser cygnoides domesticus* | 74 | 86 |
| *Coturnix japonica* | *Gallus gallus* | 33 | 42 |

**Table S25.** Primer used in the experiment in the present study.

| **Gene** | **Primers** | **Sequence (5′-3′)** | **Annealing temperature** | **Function** |
| --- | --- | --- | --- | --- |
| *TAS2R40* | *TAS2R40*-5’utr-F | ATACAAATATAGGTCTATAAAGTGGC | 55℃ | genotyping |
|  | *TAS2R40*-5’utr-R | AGCAATTATTAGGCATATAGCAGA | 55℃ | genotyping |
| *KEL* | *KEL*-intron1-F | TTATGCCTTCCCCTGCACCGTT | 58℃ | genotyping |
|  | *KEL*-intron1-R | AGGCTTTTCCGTTGAAGACATCGTT | 58℃ | genotyping |
| *NANOG* | *NANOG*-exon4-F | GCTGACTGTGCTACTTAAATGCTT | 55℃ | genotyping |
|  | *NANOG*-exon4-R | TTACATGATGCCCCTGAGTTGGAA | 55℃ | genotyping |
| *TAS2R40* | *TAS2R40*-F | CACCAGGCAGATGCAAAATAA | 60℃ | qRT-PCR |
|  | *TAS2R40-R* | TGGCTTCTCCAATGCTGAAA | 60℃ | qRT-PCR |
| *GAPDH* | *GAPDH*-F | GGTTGTCTCCTGCGACTTCA | 60℃ | qRT-PCR |
|  | *GAPDH*-R | TCCTTGGATGCCATGTGGAC | 60℃ | qRT-PCR |

**Table S26.** Statistics of Illumina sequencing data for Csp-b duck genome assembly.

| Pair-end libraries | Insert size (bp) | Total data (G) | Read length (bp) | Sequence coverage (X) |
| --- | --- | --- | --- | --- |
| Illumina reads | 350 | 20.04 | 150 | 15.78 |
| Illumina reads | 250 | 31.53 | 150 | 24.83 |
| Illumina reads | 2000 | 25.88 | 150 | 20.38 |
| Illumina reads | 5000 | 31.53 | 150 | 24.83 |
| Total | - | 108.98 | - | 85.81 |

**Table S27.** Statistics the variation type of CC duck genetic linkage map.

| Maker Type | P1 genotype | P2 genotype | Maker Number | Percentage |
| --- | --- | --- | --- | --- |
| ab x cd | ab | cd | 37 | 0.00% |
| ab x cc | ab | cc | 5204 | 0.07% |
| cc x ab | cc | ab | 5763 | 0.08% |
| ef x eg | ef | eg | 17,547 | 0.24% |
| hk x hk | hk | hk | 1,133,469 | 16.03% |
| nn x np | nn | np | 2,551,429 | 34.39% |
| lm x ll | lm | ll | 2,356,016 | 32.54% |
| aa x bb | aa | bb | 1,231,700 | 16.65% |
| Total |  |  | 7,301,165 | 100% |

**Table S28.** Summary of length and number of markers for each linkage group.

| Chromosome | Length (cM) | SNP number | Average spacing |
| --- | --- | --- | --- |
| 1 | 353.72 | 578 | 0.61 |
| 2 | 158.95 | 282 | 0.57 |
| 3 | 211.57 | 394 | 0.54 |
| 4 | 139.38 | 219 | 0.64 |
| 5 | 128.34 | 207 | 0.62 |
| 6 | 102.97 | 208 | 0.50 |
| 7 | 102.46 | 124 | 0.83 |
| 8 | 93.54 | 227 | 0.41 |
| 9 | 82.30 | 161 | 0.51 |
| 10 | 77.26 | 185 | 0.42 |
| 11 | 74.16 | 162 | 0.46 |
| 12 | 73.38 | 173 | 0.43 |
| 13 | 69.82 | 172 | 0.41 |
| 14 | 80.09 | 195 | 0.41 |
| 15 | 66.35 | 198 | 0.34 |
| 16 | 76.87 | 185 | 0.42 |
| 17 | 75.11 | 166 | 0.46 |
| 18 | 63.80 | 163 | 0.39 |
| 19 | 66.17 | 150 | 0.44 |
| 20 | 69.56 | 201 | 0.35 |
| 21 | 69.35 | 171 | 0.41 |
| 22 | 58.92 | 160 | 0.37 |
| 23 | 62.49 | 145 | 0.43 |
| 24 | 60.43 | 128 | 0.48 |
| 25 | 62.18 | 141 | 0.44 |
| 26 | 65.01 | 129 | 0.51 |
| 27 | 64.68 | 123 | 0.53 |
| 28 | 64.35 | 145 | 0.45 |
| 29 | 62.17 | 99 | 0.63 |
| 30 | 63.09 | 70 | 0.91 |
| 31 | 38.39 | 52 | 0.75 |
| 32 | 18.97 | 25 | 0.79 |
| 33 | 0.53 | 3 | 0.26 |
| 34 | 3.08 | 4 | 1.03 |
| 35 | 18.90 | 18 | 1.11 |
| 36 | 16.46 | 17 | 1.03 |
| 37 | 9.55 | 15 | 0.68 |
| In total | 2,904.38 | 5,795 | 0.50 |

**Table S29.** Classification of repeat elements in the CC duck genome.

|  | *De novo* + Repbase Length (bp) | percentage of Genome （%） | TE proteins Length (bp) | percentage of Genome (%) | Combined TEs Length (bp) | percentage of Genome （%） |
| --- | --- | --- | --- | --- | --- | --- |
| DNA | 1,914,548 | 0.17 | 224,917 | 0.02 | 2,102,402 | 0.19 |
| LINE | 82,564,375 | 7.33 | 47,683,449 | 4.23 | 90,972,533 | 8.08 |
| SINE | 159,179 | 0.01 | 0 | 0 | 159,179 | 0.01 |
| LTR | 30,805,039 | 2.74 | 7,932,962 | 0.70 | 32,906,678 | 2.92 |
| Simple repeat | 1,749,870 | 0.16 | 0 | 0 | 1,749,870 | 0.16 |
| Unknown | 8,822,025 | 0.78 | 0 | 0 | 8,822,025 | 0.78 |
| Total | 118,495,373 | 10.52 | 55,777,743 | 4.95 | 122,152,871 | 10.85 |

**Table S30.** Statistics of predicted protein-coding genes in the Chinese crested (CC) duck genome.

|  | Gene set | Gene number | Average transcript  length (bp) | Average CDS  length (bp) | Average exons  per gene | Average exon  length (bp) | Average intron  length (bp) |
| --- | --- | --- | --- | --- | --- | --- | --- |
| *De novo* | Augustus | 39,810 | 9,084.77 | 1,139.03 | 4.90 | 232.57 | 2,038.62 |
|  | GlimmerHMM | 206,079 | 4,684.80 | 513.42 | 2.69 | 191.07 | 2,472.49 |
|  | SNAP | 105,418 | 16,475.68 | 719.50 | 4.80 | 149.91 | 4,146.69 |
|  | Geneid | 46,497 | 17,297.08 | 1,227.98 | 5.28 | 232.43 | 3,751.64 |
|  | Genscan | 54,567 | 15,127.91 | 1,426.45 | 7.06 | 201.99 | 2,260.18 |
| Homolog | Acyg | 35,775 | 9,553.75 | 963.32 | 4.86 | 198.08 | 2,223.65 |
|  | Afor | 21,904 | 14,390.24 | 1,335.31 | 6.58 | 202.81 | 2,337.84 |
|  | Apla | 31,985 | 8,983.23 | 833.35 | 4.87 | 171.10 | 2,105.69 |
|  | Cjap | 19,981 | 17,085.92 | 1,432.51 | 7.66 | 186.90 | 2,348.80 |
|  | Cliv | 22,811 | 14,260.60 | 1,159.12 | 6.46 | 179.55 | 2,401.35 |
|  | Egar | 20,075 | 15,761.78 | 1,385.84 | 7.06 | 196.28 | 2,372.10 |
|  | Ggal | 35,950 | 8,891.72 | 819.56 | 4.52 | 181.35 | 2,293.64 |
|  | Hsap | 14,572 | 20,511.97 | 1,573.35 | 8.75 | 179.71 | 2,442.14 |
|  | Nnot | 19,781 | 12,015.58 | 1,257.67 | 6.05 | 207.75 | 2,128.70 |
|  | Scam | 21,095 | 15,057.91 | 1,358.22 | 6.80 | 199.63 | 2,360.45 |
|  | Tgut | 25,334 | 14,204.50 | 1,061.95 | 5.72 | 185.59 | 2,783.22 |
| RNA-seq | PASA | 100,396 | 14,329.69 | 1,055.58 | 6.00 | 175.87 | 2,653.80 |
|  | Cufflinks | 87,156 | 20,594.98 | 3,406.92 | 6.86 | 496.33 | 2,930.99 |
| EVM | | 22,524 | 17,728.93 | 1,322.28 | 7.70 | 171.77 | 2,449.48 |
| PASA-update | | 22,360 | 19,350.63 | 1,338.06 | 7.74 | 172.88 | 2,672.60 |
| Final set | | 17,425 | 24,484.99 | 1,599.03 | 9.58 | 166.86 | 2,666.39 |

Note: Acyg represents *Anser cygnoides* domesticus; Afor for *Aptenodytes forsteri*; Apla for *Anas platyrhynchos* domestica; Cjap for *Coturnix japonica*; Cliv for *Columba livia*; Egar for *Egretta garzetta*; Ggal for *Gallus gallus*; Hsap for *Homo sapiens*; Nnot for *Nestor notabilis*; Scam for *Struthio camelus*; Tgut represents *Taeniopygia guttata*.

**Table S31.** Predicted non-coding RNAs in the CC duck genome.

| Type | | Number | Average length (bp) | Total length (bp) | % of genome |
| --- | --- | --- | --- | --- | --- |
| miRNA | | 359 | 87.52 | 31,420 | 0.00279 |
| tRNA | | 402 | 74.54 | 29,964 | 0.002661 |
| rRNA | rRNA | 79 | 297.44 | 23,498 | 0.002086 |
|  | 18S | 14 | 565.29 | 7,914 | 0.000703 |
|  | 28S | 55 | 263.62 | 14,499 | 0.001287 |
|  | 5.8S | 1 | 156 | 156 | 0.000014 |
|  | 5S | 9 | 103.22 | 929 | 0.000082 |
| snRNA | snRNA | 308 | 126.91 | 39,087 | 0.003471 |
|  | CD-box | 114 | 91.52 | 10,433 | 0.000926 |
|  | HACA-box | 82 | 144.94 | 11,885 | 0.001055 |
|  | splicing | 94 | 144.77 | 13,608 | 0.001208 |

**Table S32.** Summary of predicted protein-coding genes from different public databases.

|  | Number | Percent (%) |
| --- | --- | --- |
| Total | 17,425 | - |
| Swiss-Prot | 15,880 | 91.1 |
| NR | 16,547 | 95.0 |
| KEGG | 14,290 | 82.0 |
| InterPro | 15,579 | 89.4 |
| GO | 11,565 | 66.4 |
| Pfam | 14,230 | 81.7 |
| Annotated | 16,577 | 95.1 |
| Unannotated | 848 | 4.9 |

Note: NR represent Non-redundant database; KEGG for Kyoto Encyclopedia of Genes and Genomes database; GO represent Gene Ontology.

**Table S33.** The statistic results of gene organization in relative species.

| Gene set | Number of genes | CDS+ intron length (average) | CDS length (average) | exon length (average) | intron length (average) | Exons per gene (average) |
| --- | --- | --- | --- | --- | --- | --- |
| Acyg | 14,696 | 30,176.71 | 1,763.51 | 162.58 | 2,885.50 | 10.85 |
| Apla | 15,634 | 21,091.75 | 1,507.51 | 148.94 | 2,147.05 | 10.12 |
| Csp-b duck | 15,278 | 22,327.78 | 1,638.79 | 176.41 | 2,495.83 | 9.29 |
| CC duck | 17,425 | 24,484.99 | 1,599.03 | 166.86 | 2,666.39 | 9.58 |
| Ggal | 16,362 | 23,425.95 | 1,603.16 | 165.66 | 2,514.86 | 9.68 |
| Tgut | 16,348 | 25,502.91 | 1,620.25 | 161.88 | 2,650.91 | 10.01 |

Note: Acyg represents *Anser cygnoides* domesticus; Apla for *Anas platyrhynchos* domestica; Ggal for *Gallus gallus*; Tgut represents for *Taeniopygia guttata*.

**Table S34.** The statistical results of Csp-b duck gene function annotation.

| Database | Annotated Number | Annotated Percent (%) |
| --- | --- | --- |
| NR | 15,274 | 100 |
| Swiss-Prot | 14,455 | 94.6 |
| KEGG | 13,913 | 91.1 |
| InterPro | 14,862 | 97.3 |
| GO | 10,992 | 71.9 |
| Annotated | 13,359 | 87.4 |
| Total | 15,278 | - |

Note: NR represent Non-redundant database; KEGG for Kyoto Encyclopedia of Genes and Genomes database; GO represent Gene Ontology.

**Table S35.** The statistical results of Chinese spot-billed (Csp-b) duck non-coding RNA.

| Type | | Number | Average length (bp) | Total length (bp) | percentage of genome (%) |
| --- | --- | --- | --- | --- | --- |
| miRNA | | 345 | 85.0870 | 29,355 | 0.002663 |
| tRNA | | 198 | 74.9242 | 14,835 | 0.001346 |
| rRNA | rRNA | 56 | 146.1607 | 8,185 | 0.000742 |
|  | 18S | 14 | 116.3571 | 1,629 | 0.000148 |
|  | 28S | 39 | 160.9231 | 6,276 | 0.000569 |
|  | 5.8S | 1 | 155.0000 | 155 | 0.000014 |
|  | 5S | 2 | 62.5000 | 125 | 0.000011 |
| snRNA | snRNA | 237 | 117.5949 | 27,870 | 0.002528 |
|  | CD-box | 90 | 84.2111 | 7,579 | 0.000688 |
|  | HACA-box | 72 | 138.0000 | 9,936 | 0.000901 |
|  | splicing | 58 | 129.0517 | 7,485 | 0.000679 |

**Table S36.** Statistics of Benchmarking Universal Single-Copy Orthologs (BUSCO) assessment of Chinese spot-billed (Csp-b) duck genome assembly and gene set prediction.

| Species | Genome BUSCO notation assessment results |
| --- | --- |
| Csp-b duck | C:91.8%[S:91.4%,D:0.4%],F:6.7%,M:1.7%,n:2586 |

Note: C: Complete BUSCOs

S: Complete Single-Copy BUSCOs

D: Complete Duplicated BUSCOs

F: Fragmented BUSCOs

M: Missing BUSCOs

n: Total BUSCO groups searched

Table S38 The list the recently publish genome quality

| Name of Gneome | Sample_ID |  | Total | Max | Number>=2000 | N50 | N60 | N70 | N80 | N90 |
| --- | --- | --- | --- | --- | --- | --- | --- | --- | --- | --- |
| CAU_PK_V1 | length | Contig | 1,080,105,771 | 929,952 | - | 88,037 | 67,764 | 49,890 | 33,462 | 16,380 |
|  |  | Scaffold | 1,136,415,614 | 202,396,406 | - | 74,988,519 | 64,310,432 | 32,166,367 | 20,768,007 | 8,175,025 |
|  | number | Contig | 44,791 | - | 28,723 | 3,342 | 4,741 | 6,598 | 9,225 | 13,710 |
|  |  | Scaffold | 21,240 | - | 9,241 | 5 | 6 | 9 | 14 | 22 |
| CAU_Wild | length | Contig | 1,210,757,325 | 73,505,640 | - | 11,301,519 | 8,373,174 | 6,305,876 | 3,890,484 | 1,365,943 |
|  |  | Scaffold | 1,211,992,756 | 208,326,429 | - | 77,626,585 | 39,543,408 | 26,742,597 | 18,227,546 | 7,574,731 |
|  | number | Contig | 1,983 | - | 1,917 | 34 | 46 | 63 | 87 | 137 |
|  |  | Scaffold | 1,665 | - | 1,599 | 5 | 7 | 10 | 16 | 26 |
| NC_PK | length | Contig | 1,184,302,851 | 28,495,473 | - | 5,679,408 | 4,212,680 | 2,399,784 | 1,418,083 | 528,888 |
|  |  | Scaffold | 1,188,533,289 | 207,238,429 | - | 76,269,206 | 66,856,311 | 26,788,864 | 20,984,523 | 11,966,879 |
|  | number | Contig | 1,661 | - | 1,651 | 57 | 81 | 119 | 182 | 313 |
|  |  | Scaffold | 756 | - | 756 | 5 | 6 | 10 | 15 | 22 |
| CAU_PK_V2 | length | Contig | 1,185,139,460 | 55,383,155 | - | 16,035,322 | 11,859,130 | 8,518,109 | 4,923,319 | 1,728,104 |
|  |  | Scaffold | 1,186,367,508 | 207,246,783 | - | 76,279,691 | 39,447,372 | 26,944,867 | 20,392,265 | 7,681,837 |
|  | number | Contig | 622 | - | 619 | 24 | 33 | 45 | 64 | 101 |
|  |  | Scaffold | 421 | - | 418 | 5 | 7 | 10 | 15 | 24 |
| CAU_Laying | length | Contig | 1,210,592,770 | 53,929,205 | - | 8,556,976 | 5,803,148 | 4,091,236 | 2,746,911 | 1,074,519 |
|  |  | Scaffold | 1,217,695,176 | 212,526,513 | - | 76,919,215 | 39,707,377 | 26,964,064 | 18,828,828 | 7,192,834 |
|  | number | Contig | 859 | - | 852 | 40 | 57 | 81 | 117 | 187 |
|  |  | Scaffold | 294 | - | 288 | 5 | 7 | 10 | 16 | 26 |
